# Mapping Transcriptomic Vector Fields of Single Cells

**DOI:** 10.1101/696724

**Authors:** Xiaojie Qiu, Yan Zhang, Shayan Hosseinzadeh, Dian Yang, Angela N. Pogson, Li Wang, Matt Shurtleff, Ruoshi Yuan, Song Xu, Yian Ma, Joseph M. Replogle, Spyros Darmanis, Ivet Bahar, Jianhua Xing, Jonathan S Weissman

## Abstract

Single-cell RNA-seq, together with RNA velocity and metabolic labeling, reveals cellular states and transitions at unprecedented resolution. Fully exploiting these data, however, requires dynamical models capable of predicting cell fate and unveiling the governing regulatory mechanisms. Here, we introduce *dynamo*, an analytical framework that reconciles intrinsic splicing and labeling kinetics to estimate absolute RNA velocities, reconstructs velocity vector fields that predict future cell fates, and finally employs differential geometry analyses to elucidate the underlying regulatory networks. We applied *dynamo* to a wide range of disparate biological processes including prediction of future states of differentiating hematopoietic stem cell lineages, deconvolution of glucocorticoid responses from orthogonal cell-cycle progression, characterization of regulatory networks driving zebrafish pigmentation, and identification of possible routes of resistance to SARS-CoV-2 infection. Our work thus represents an important step in going from qualitative, metaphorical conceptualizations of differentiation, as exemplified by Waddington’s epigenetic landscape, to quantitative and predictive theories.

## INTRODUCTION

A hallmark of metazoans is the striking ability of a single zygote to differentiate into a multitude of distinct cell types while maintaining the same genomic sequence. To illustrate this process, in the 1950s, Waddington introduced the epigenetic landscape, a metaphor in which differentiation proceeds like a ball sliding downhill into various valleys (Waddington 1957). In this conception, developmental processes are depicted as autonomous and precisely controlled. The epigenetic landscape has been used to intuitively explain cell differentiation (Huang et al. 2007), and more recently transdifferentiation or reprogramming (Cahan et al. 2014); however, a central goal of the field remains to move beyond such a qualitative, metaphorical conceptualization toward more quantitative, predictive models.

Mathematical modeling, especially in conjunction with dynamical systems theories (Brauer and Kribs 2015), provides a powerful tool for gaining mechanistic insights into how gene regulatory networks (GRNs) control various biological processes, including not only the current cell state but also the direction in which it might be headed (Elowitz and Leibler 2000; Alon 2006; Huang 2012). In a dynamical systems formalism, one can represent the state of each cell as a vector (x) in a multi-dimensional gene expression space in which the elements are the instantaneous concentrations of molecules (e.g., copies of mRNAs, proteins, etc. per cell). Ignoring cellular stochasticity, the evolution of the cell state over time, or its velocity (***ẋ***(*t*), is governed by a set of ordinary differential equations (ODEs) determined by the underlying GRN, expressed as ***ẋ***(*t*) = ***f***(***x***(*t*)), where *f* is a function of the instantaneous velocity of cell states. Although some efforts have been made to study structure-function relationships among network motifs such as the toggle-switch network (Huang et al. 2007) central to cell fate decisions (**Fig. 1A**), and to perform whole-cell simulations of bacteria such as *Mycoplasma genitalium* and *Escherichia coli* (Karr et al. 2012; Macklin et al. 2020), it remains a grand challenge to reconstruct the vector field function representing the time evolution of a genome-wide expression state in complex systems such as a mammalian cell from experimental data.

**Figure 1:**
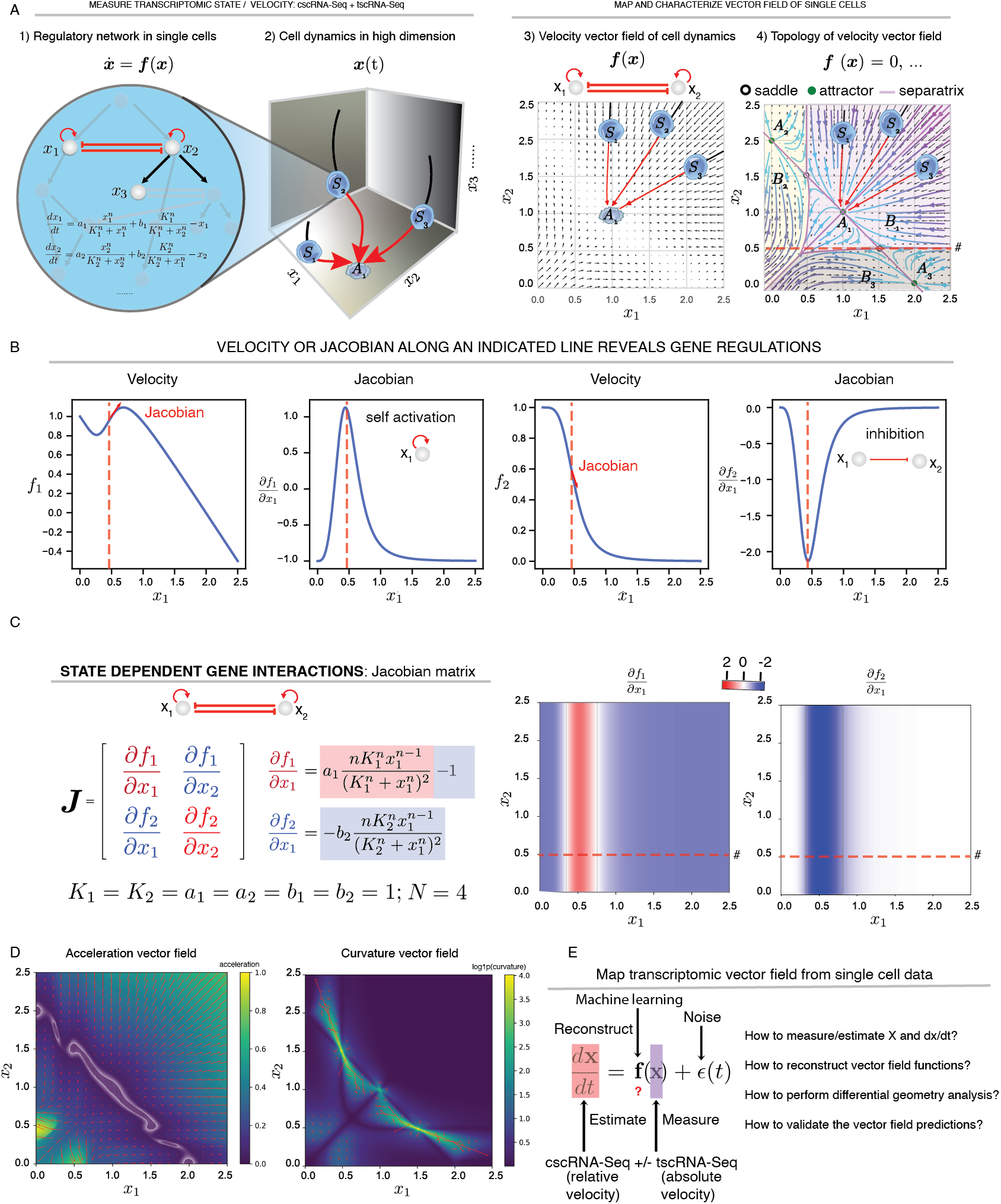
Modeling and Interpreting Single-cell Gene Expression Dynamics using Velocity Vector Field Functions and Differential Geometry Analyses. A. **Cell state transition under dynamical systems framework. 1**) A canonical toggleswitch motif of two genes (whose instantaneous expression levels, e.g. RNA copies in a single cell, are denoted as *x*_1_, *x*_2_) and one of their downstream targets (with expression level of *x*_3_) are embedded in a complex regulatory network. The toggle-switch network motif is used throughout this figure and **Fig. 4. 2**) Cell fate transitions as trajectories in a high-dimensional state space spanned by state descriptors (e.g. expression of RNA or protein). The regulatory network determines the possible trajectories and final fates. A three-dimensional state space captures the dynamics of the highlighted three-gene system from **1**. Any point in this space represents a network state *S*(*t*) = (*x*_1_, *x*_2_, *x*_3_) at time *t*. Three example states *S*_1_, *S*_2_, and *S*_3_, are shown. As in most states, the three states do not necessarily represent stable network states; therefore, they are driven by the network interactions to move along trajectories (**red solid** lines, i.e. streamlines) that converge to the stable attractor state, *A*_1_. Note that the **red** and **black** segments of the trajectories correspond to past and future cell states, respectively. **3**) Global view of cell dynamics via vector field functions. **4**) Topological features of the vector field. Important features include: **steady states** [including attractors (filled circles, *ẋ*(*t*) = 0, *ẍ*(*t*) < 0), and saddle points (unfilled circles, *ẋ*(*t*) = 0, *ẍ*(*t*) > 0], **attractor basins** (domains where all points will be eventually attracted to a particular attractor state, shaded with different colors), **separatrices** (lines or planes that set the boundary between attractor basins, shown as a red line), and **nullclines** (lines or planes where the velocity of one particular dimension is 0). B. **Velocity and Jacobian along the horizontal dashed line indicated in A4.** Calculating the derivative of the velocity of *x*_1_, *f*_1_ (first panel) or that of *x*_2_, *f*_2_ (third panel) along the indicated line gives rise to the Jacobian terms ***J***_11_ or ***J***_21_, which reveals the self-activation of gene *x*_1_, and inhibition of *x*_2_ by *x*_1_, respectively. C. The Jacobian (the matrix **J** = ∇**f**(**x**), where the *ij*-th element is **J_ij_** = *∂**f**_i_*/*∂x_j_* (**left**), of a vector field function reflects state-dependent gene interactions in the state space, represented as a heatmap (**right**). D. Acceleration and curvature vector fields of single-cell gene expression. Color of the heatmaps corresponds to length of the acceleration and curvature vectors at each point in the state space. For **C**/**D**, see more details in **Box 1**. E. Summary of the task of mapping the vector field functions from transcriptomic data, formulated as a machine learning problem, with downstream validations and analyses.

Recent developments in single-cell genomics have enabled profiling of cell states at unprecedented resolution (Macosko et al. 2015; Klein et al. 2015; Cao, O’Day, et al. 2020). These advances provide new opportunities to infer the precise time evolution of cell fate transitions, in a way fundamentally related to vector field reconstruction. Single-cell profilings, of the transcriptome (scRNA-seq) or other genomic features [e.g., chromatin accessibility (Cusanovich et al. 2015; Buenrostro et al. 2015)], multi-omics [transcriptomic, proteomic, or epigenetic state (Stoeckius et al. 2017)], and spatial transcriptomics [e.g., MERFISH (Chen et al. 2015), seqFISH (Lubeck et al. 2014), Slide-seq (Rodriques et al. 2019), and ZipSeq (Hu et al. 2020)] provide complementary transcriptomic, epigenetic, multimodal, and spatial information about single cells. However, due to the destructive nature of these approaches, it is generally impossible to follow the same cell over time. On the other hand, live-cell imaging allows one to track the co-expression of a small number of molecular and cellular features (W. Wang et al. 2020) in many cells over time, but it does not lend itself to visualizing the full transcriptomic states.

Advances in single-cell profiling have fueled the development of computational approaches for inferring cellular dynamics from snapshot measurements. Chief among them are pseudotime-based methods (Trapnell et al. 2014; Saelens et al. 2019) first developed to infer the order of biological progression by learning a graph manifold of single cells based on transcriptome similarity. However, pseudotime ordering is limited to the analysis of central trends of biological progressions rather than the precise dynamics of individual cells over real time, and it is not generally suitable for resolving the directionality of biological processes (Saelens et al. 2019; X. Qiu et al. 2020). A second major advance has been the development of RNA velocity, which predicts the cell RNA expression states in the near future by explicitly exploring the intrinsic splicing kinetics incidentally captured by most scRNA-seq protocols. Efforts have been made to extend “RNA velocity” to “protein velocity” (Gorin, Svensson, and Pachter 2020) or non-stationary states (Bergen et al. 2020). Such methods provide a vantage on the short-term evolution of individual cell states, but have intrinsic limitations (see **RESULTS** and **MATERIAL AND METHODS**) that make it challenging to predict the continuous evolution of historical and future states over a long period of time. Recently, several groups have adapted bulk RNA-seq based on RNA metabolic labeling to single-cell approaches (Erhard et al. 2019; Hendriks et al. 2019; Cao, Zhou, et al. 2020; Battich et al. 2020; Q. Qiu et al. 2020). The ensuing ability to obtain time-resolved scRNA-seq (tscRNA-seq) provides further qualitative measures of cell state and its velocity by distinguishing “new” and “old” RNA molecules in an experimentally programmable manner. Thus, these methods in principle provide the data necessary for accurate reconstruction of transcriptomic vector fields. However, mathematical models and tools for integrating labeling-based tscRNA-seq and splicing-based conventional scRNA-seq (cscRNA-seq) toward properly inferring the RNA turnover rates and estimating RNA velocity remain undeveloped, as do methods for using such information to construct continuous vector fields. Finally, it remains unknown whether it is possible to leverage vector field functions to gain quantitative, predictive, and functional understanding of cell state transitions, and if so, how. Thus, despite striking advances in single-cell profiling, our ability to fully exploit these measurements is limited by the lack of an appropriate analytical framework for interpreting the data and guiding future experiments.

Here, we introduce a comprehensive computational framework for constructing and interpreting single-cell transcriptomic vector fields. The framework addresses three challenges. First, by integrating RNA metabolic labeling and intrinsic splicing kinetics, we build an inclusive model of expression dynamics capable of accurately estimating genome-wide RNA turnover rates, as well as absolute RNA velocities. Second, we develop a general algorithm for robustly reconstructing the transcriptomic vector field functions from sparse, noisy single-cell measurements. Third, we marry the scalability of machine learning-based vector field reconstruction methods with the interpretability of differential geometry analyses, including Jacobian, acceleration, and divergence, to gain further biological insights.

Applications of the newly introduced computational framework to many metabolic labeling–based scRNA-seq datasets that involve various labeling strategies, demonstrate its generalizability and robustness in accurately estimating RNA kinetic rate constants and absolute RNA velocities. Our framework permits us to go beyond discrete velocity samples, to a continuous vector field function in a high-dimensional gene expression space. We validated the vector field trajectory predictions over several days by using clone tracing with DNA barcodes and sequential capture of single cells during neutrophil lineage commitment of HL60 cells as well as during murine hematopoiesis. Differential geometry analyses further proved to provide functional insights into a variety of conventional and metabolic labeling–based scRNA-seq experiments, ranging from zebrafish pigment cell differentiation (Saunders et al. 2019) to murine bidirectional transition between ESC pluripotent cells and a 2C-like totipotent state (Q. Qiu et al. 2020). In addition, we show that our framework facilitates the discovery of potential viral–host interactions and mechanisms of resistance to SARS-Cov-2 infections.

This new framework represents a significant advance from the metaphor of epigenetic landscape to a quantitative and predictive theory of the time evolution of single cell transcriptomics, applicable to many biological systems/processes and at genome-wide scale. We have made the associated computational framework as an open-source software, ***dynamo***, available at https://github.com/aristoteleo/dynamo-release.

## RESULTS

### Differential Geometry Analyses of Velocity Vector Fields Yield Quantitative Information about Gene Regulation

In principle, a velocity vector field (**Box 1**) provides a complete description of the way genes regulate each other. While our discussion of the vector field in this study focuses on transcriptomic space, vector fields can be generally applicable to other spaces, such as proteomic, or metabolic space. As a simple example, consider a two-gene toggle-switch motif (Huang et al. 2007) that appears frequently in cell differentiation such as the *PU.1/Gata-1* motif that controls the bifurcation of hematopoietic progenitors into either the myeloid or erythroid lineage (**Fig. 1A1**). The vector field function for this motif is often formulated as a set of deterministic ODEs (i.e., ignoring biological stochasticity) (**Fig. 1A1**), which model the self-activation and mutual inhibition between *PU.1* and *Gata-1*, specify the instantaneous velocity of a cell at any given expression state, and predict the evolution of the cell state over time (**Fig. 1A2–4**). One can further characterize the topology of this vector field in its gene expression space with separatrices that divide the space into three attractor basins, each containing a stable fixed point (the attractor) corresponding to a stable phenotype (**Fig. 1A4**). We illustrate three representative cells that start from different states of the same attractor basin of attractor *A*_1_, each propagating along a trajectory (streamline) defined by the vector field function to settle at the same attractor state *A*_1_ (**Fig. 1A2–4, Fig. SI1A**). By contrast, saddle points are unstable fixed points located on sepatrices connecting pairs of attractors.

#### Box 1: Differential Geometry of Vector Fields

In this work, we introduced **dynamical systems theory** and **differential geometry** analysis to single-cell genomics. A dynamical system describes the time dependence of a point in a geometrical space, e.g., planetary motion or cell fate transitions, whereas differential geometry uses the techniques of differential/integral calculus and linear/multilinear algebra to study problems in geometry, e.g., the topology or geometric features along a streamline in vector field of the gene expression space. A vector field function ***f***, a fundamental topic of dynamical systems theories, takes spatial coordinate input ***x*** (e.g., single-cell expression in gene state space) in a-dimensional space (each gene corresponds to a dimension) as input and outputs a vector ***υ*** (e.g., corresponds to gene expression velocity vector from a single cell) in the same space, i.e. ***υ*** = ***f***(***x***). In this study, we specifically discuss velocity vector fields that can be used to derive acceleration and curvature vector fields (see **below**). With analytical velocity vector field functions, including the ones that we learned directly from data, we can move beyond velocity to high-order quantities, including the Jacobian, divergence, acceleration, curvature, curl, etc., using theories developed in differential geometry. The discussion of the velocity vector field in this study focuses on transcriptomic space; vector fields, however, can be generally applicable to other spaces, such as morphological, proteomic, or metabolic space.

Because ***f*** is a vector-valued multivariate function, a *d×d* matrix encoding its derivatives, called the *Jacobian*, plays a fundamental role in differential geometry analysis of vector fields:

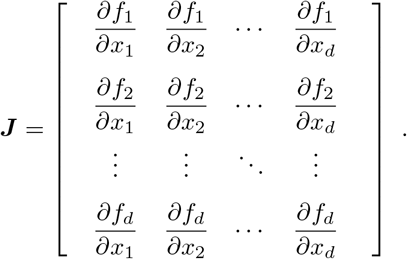

A Jacobian element *∂f_i_*/*∂x_j_* reflects how the velocity of *x_i_* is impacted by changes in *x_j_*.

The trace of the Jacobian is divergence:

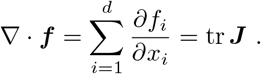

Divergence measures the degree of “outgoingness” at any point, summarized in **Box Fig. 1A**.

**Box Fig. 1.**
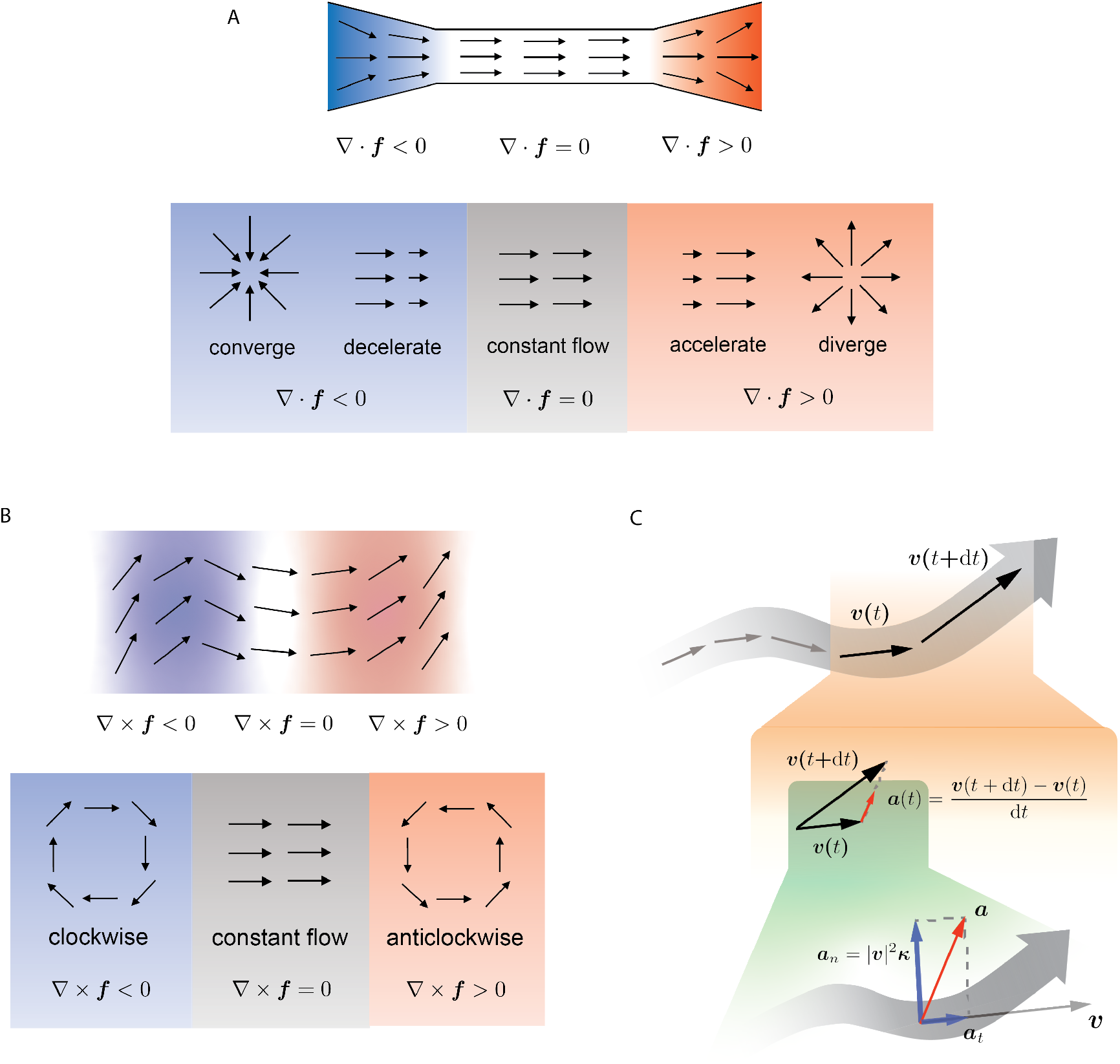
Divergence, curl, acceleration and curvature of vector field.

By definition, an attractor (repulsor) converges (diverges) in any direction. Note that it is possible to have a point where the vectors converge in one direction but diverge in another, a case that is not depicted in the diagram above. This means that although an attractor (repulsor) always has negative (positive) divergence, the opposite does not necessarily hold.

*Curl* is a quantity measuring the degree of rotation at a given point in the vector field. It is well-defined only in two or three dimensions (e.g. two or three reduced principal components or UMAP components):

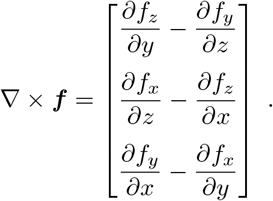

The behavior of curl is summarized in **Box Fig. 1B**.

Many differential geometry quantities are defined on *streamlines*, which are curves everywhere tangent to the vector field. The streamlines can be parametrized with time *t*, denoted ***x***(*t*), as they are essentially trajectories of cells moving in the vector field. In practice, they are often calculated using numerical integration methods, e.g., the Runge–Kutta algorithm. The *acceleration* is the time derivative of the velocity, as shown in **Box Fig. 1C** (orange shade), and can be defined as:

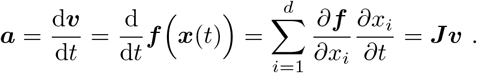

The curvature vector (**Box Fig. 1C**, green shade) of a curve is defined as the derivative of the unit tangent vector 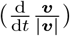, divided by the length of the tangent (|***υ***|):

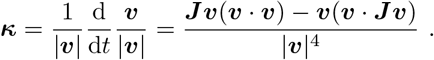

In the context of velocity vector fields and streamlines, the unit tangent vector is the normalized velocity. By definition, acceleration measures the rate of change of velocity in terms of both its magnitude and direction. Curvature, on the other hand, measures only the change in direction, as the velocity vector is normalized. **Box Fig. 1C** (green shade) illustrates how the acceleration can be decomposed into a tangential and a radial component, and the latter is connected to the curvature:

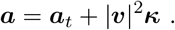

Although acceleration and curvature are mathematically defined on streamlines, the actual calculation, as shown above, can be done pointwise using only the velocity and the Jacobian evaluated at the point of interest. Because the acceleration or the curvature can be calculated for any point in the state space, one obtains the acceleration or curvature vector field.

Other relevant differential geometric analyses, including torsion (applicable to three dimensional vector field), vector Laplacian, etc., can also be computed using vector field functions, although they were not extensively studied in this work.

Analyses of the vector field can also yield deep insights and help generate hypotheses about how genes regulate cell states (**Box 1, Fig. 1B, C**). For example, the Jacobian can be used to investigate the cell state–dependent interactions because it is tightly related to the underlying regulatory network. Specifically, the *ij*—th element **J_ij_** = *∂**f**_i_*/*∂x_j_* of the Jacobian matrix (**J**_*ng×ng*_, *ng* is the number of genes) indicates the change in the velocity of gene *i* with respect to an increase in expression of gene *j* at a given cell state *x*, with a positive (or negative) value for activating (or inhibitory) regulation (**Fig. 1B**). Moreover, the maximum of |*J_ij_*| indicates where gene *j* has the strongest effect (activation or inhibition) on gene *i* (**Fig. 1B, C, Fig. SI1B**). In the toggle-switch model, the Jacobian analysis correctly identifies self-activation and mutual inhibition, with the strongest regulation taking place when *x*_1_ and *x*_2_ are about 0.5 (**Fig. 1B, C, Fig. SI1B**).

A number of additional differential geometric quantities provide complementary information of gene regulation dynamics and cell fate commitment. The **acceleration** field (**Box 1, Fig. 1D**) reveals gene expression subspaces (i.e., hotspots of cells states) where the velocities change dramatically, either in magnitude or direction, due to the alteration of dynamical regulations (e.g., the two symmetric regions in the bottom left corner where the expression level of either *x*_1_ or *x*_2_ increases rapidly). When a cell leaves an unstable state (e.g., a progenitor) and moves toward a stable attractor state (e.g., a mature cell type), its velocity tends to increase before it slows down in the attractor state (**Fig. 1D**). Therefore, it is possible to detect genes that have a large value for acceleration (in magnitude) in progenitor states, making key contributions to cell fate commitment, long before cells exhibit discernible lineage-specific gene expression differences. A related but different quantity is the **curvature** field (**Box 1, Fig. 1D**), which reveals gene expression hotspots where the velocity changes direction abruptly, e.g., in regions around unstable fixed points where one or more genes’ expression changes from induction to repression or *vice versa* (**Fig. 1D**, see especially the regions of the largest curvature, which coincide with the two saddle points). The genes that strongly contribute to the curvature are regulatory genes that steer the cell fate. Curl and divergence (**Box 1, Fig. SI1C**), respectively, characterize the infinitesimal rotation of a cell state in the vector field and the local flux exiting versus entering an infinitesimal region in the expression space – the “outgoingness”. The sources (sinks) of a dynamical system often have strong positive (negative) divergence. Thus, divergence of single cells can be used to identify the possible progenitors or terminal cell types of a differentiation system.

The toggle-switch motif illustrates the significance of vector fields and various differential geometry analyses in studying the dynamics of a regulatory network. However, such simplified motifs are embedded within a complex but unknown genome-wide regulatory network. Thus, instead of constructing explicit kinetic functions that require gene interactions to be known as a prior, it is desirable to apply machine learning methods to reconstruct the transcriptomic vector field functions directly from single-cell measurements. To achieve this overarching goal (**Fig. 1E**), we need 1) the capability to accurately measure the current gene expression state and some local dynamics of cell state, so that we can reliably estimate its velocity vectors; 2) a method that can learn the continuous transcriptomic vector field functions from sparse and noisy singlecell state and velocity measurements; 3) an efficient framework for extracting the differential geometry features (e.g., the Jacobian) and relating them to the underlying mechanism of the biological system. Once those are met, we also need to demonstrate the generality of our method in a set of experimental datasets, as well as to validate our vector field prediction, especially the long-term dynamics of cell states, on well-designed experiments.

### An inclusive Model of RNA Labeling and Expression Kinetics Allows Genome-wide Estimation of RNA Synthesis and Turnover Rates

Various types of single-cell transcriptomic profiling data can be used for RNA velocity estimation. The original RNA velocity method (La Manno et al. 2018) used incidentally captured intron reads from conventional scRNA-seq (cscRNA-seq) data and assumed a universal splicing rate constant. Mainly, the kinetics of RNA transcription, splicing, and degradation obey the following ODEs:

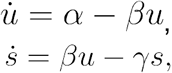

where *u* and *s* are the copies of unspliced and spliced RNAs for a particular gene in the cell, and *α, β*, and *γ* are the respective rate constants for transcriptional, splicing, and degradation (See **SUPPLEMENTARY METHODS** for a discussion on the rate and rate constant and the units of those parameters). Assuming cells with extreme high expressions of unspliced and spliced RNA expressions (top right corner of the *u, s* phase plane) are at pseudo-steady state 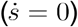, and using the substitution 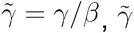 can be estimated with a linear regression of cells at steady states. Thus, the conventional RNA velocity as defined in the original study (La Manno et al. 2018) is given by:

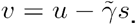

The resultant degradation rate constants and velocities from conventional RNA velocity method are therefore relative, and scaled by the gene-specific splicing rate constant *β* (See **MATERIAL AND METHODS**). We reason that such limitations can be relaxed with scRNA-seq augmented with RNA metabolic labeling (time-resolved scRNA-seq or tscRNA-seq), which measures RNA turnover dynamics in a controllable, less biased, and time-resolved fashion (See **MATERIAL AND METHODS**). The first tscRNA-seq study, scSLAM-seq (Erhard et al. 2019), introduced a new form of RNA velocity, NTR (new to total ratio) velocity, by replacing unspliced and spliced counts with new and total RNA to analyze the response of mouse fibroblast cells to cytomegalovirus (CMV) infection. However, a comprehensive estimation framework that allows genome-wide estimation of RNA synthesis and turnover rates for both cscRNA-seq and tscRNA-seq has not yet been established.

To develop a unified framework for extracting RNA dynamics information from cscRNA-seq and tscRNA-seq datasets, we constructed an inclusive model (**Fig. 2A**) that considers on-off switching of promoter state, RNA metabolic labeling (when using tscRNA-seq data), RNA splicing and degradation, and even translation and protein degradation when transcriptomic– proteomic coassay data are available. To account for different data types and experiments, we further implement three reduced models: **Model 1** considers RNA transcription, splicing and degradation, but not RNA metabolic labeling, and is tailored for cscRNA-seq, whereas both **Models 2** and **3** are tailored for tscRNA-seq with metabolic labeling, with the difference that only **Model 3** considers RNA splicing (**Fig. SI2A**).

**Figure 2:**
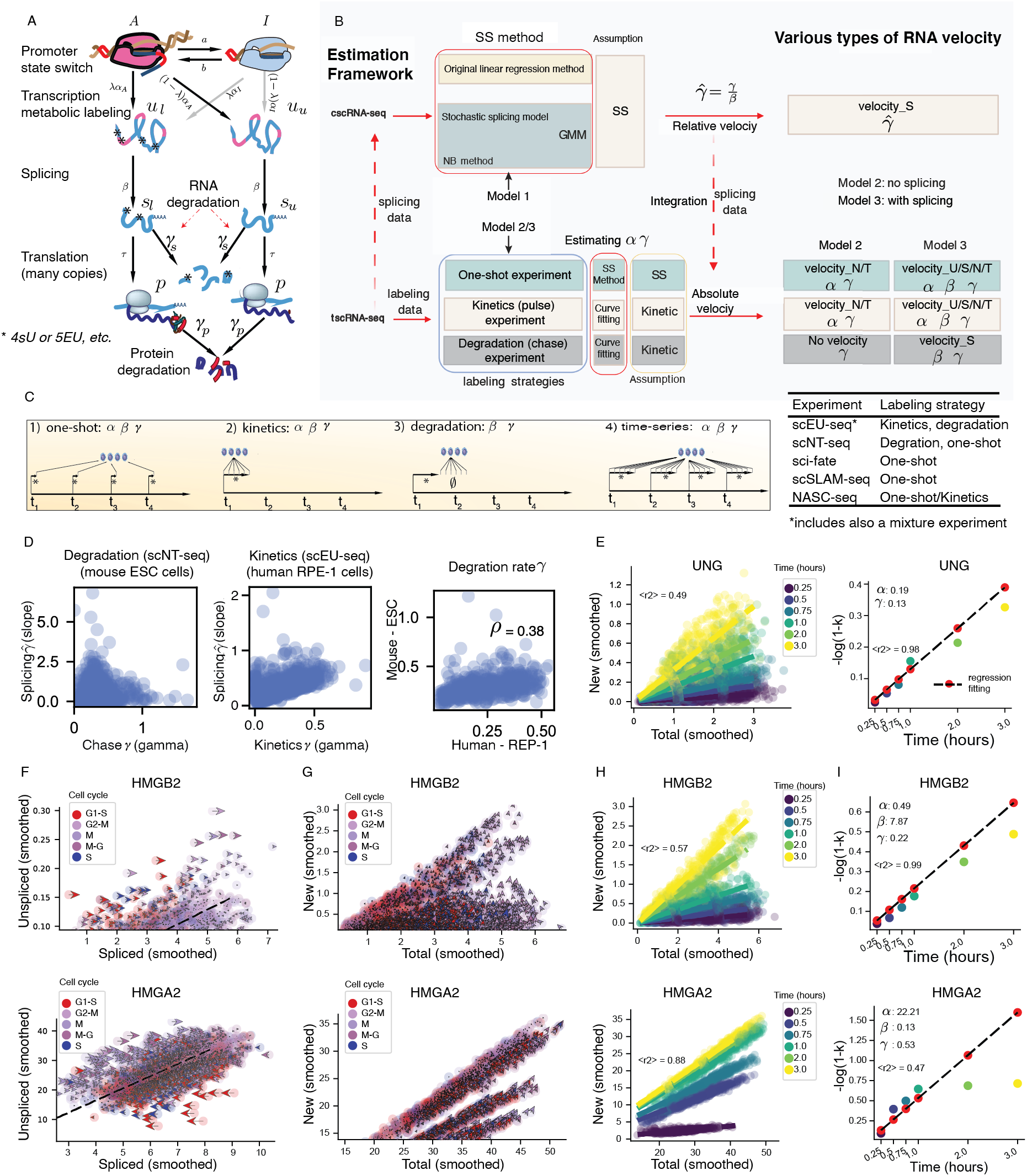
Inclusive Model of Expression Dynamics Incorporates RNA Metabolic Labeling in Single Cells. A. A comprehensive model of gene expression kinetics that includes promoter state switch, RNA metabolic labeling, RNA transcription, splicing, translation, and RNA/protein degradation. *u_u_, u_l_, s_u_*, and *s_l_* are respectively unspliced unlabeled, unspliced labeled, spliced unlabeled, and spliced labeled RNA. Note that unspliced (*u*) and spliced (*s*) RNA are the sums of *u_u_, u_l_* and *s_u_, s_l_*, respectively. Similarly, labeled RNA (*o*) and total *t* RNA are the sums of *u_l_, s_l_* and *u, s*, respectively. B. ***Dynamo***’s comprehensive estimation framework of kinetic parameters for time-resolved metabolic labeling–based scRNA-seq or tscRNA-seq experiments, and conventional scRNA-seq experiments without metabolic labeling (cscRNA-seq). GMM: generalized methods of moments; NB: negative binomial; SS: steady state; velocity_N: new RNA velocity; velocity_T: total RNA velocity; velocity_U: unspliced RNA velocity; velocity_S: spliced RNA velocity. Description of **Model 1/2/3** can be found in **Fig. SI2A**. C. Typical RNA metabolic labeling strategies and their application in published tscRNA-seq studies. On the left, **One-shot** labeling experiments (an experiments with a single RNA labeling period): estimating *α, γ* with labeling data and *β* when combining labeling and splicing data; **kinetics** labeling experiments (a time-series of 4sU or other nucleotide analog treatment): same as **one-shot labeling**; **degradation** labeling experiments (a time-series with an extended 4sU or other nucleotide analog treatment period, followed by chase at multiple time points): estimating *γ* with labeling data and *β* when combining labeling and splicing data; **Multi-time-series** labeling experiments (single cell samples are collected at multiple time points, each with a kinetics experiment): same as **one-shot labeling**. The table on the right summarizes the main labeling strategies used in all published tscRNA-seq studies. D. Comparing degradation rate constants (*γ*) calculated from tscRNA-seq data and the scaled degradation rate constants 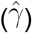 from the corresponding splicing data, and those from human cells or mouse cells. Scatter plot of the absolute degradation rate constants based on labeling data and the scaled degradation rate constants based on the corresponding splicing data in mouse ESC cells from scNT-seq study (**left**) or the human RPE-1 cells from the scEU-seq study (**middle**). Degradation rate constants from murine ESC cells tend to show a global increase compared to that from the human RPE-1 cells (**right**). E. Two-step method (see **MATERIAL AND METHODS**) of the kinetics experiment [data from scEU-seq study (Battich et al. 2020)]: **1**) A strong linearity in the new–total RNA phase plane of gene *UNG* with ascending slope for longer labeling times; **2**) A strong linearity between −*log*/(1 − *k*) and labeling time period *t* for the *UNG* gene. “Smoothed” means that the expression was locally averaged based on a *k*-nearest neighbor graph in the reduced principal component (PC) space across cells, the same applies to the **F**, **G**, **H**. F. Phase portraits of spliced-unspliced RNA planes of *HMGB2* and *HMGA2*, genes for which splicing is faster or slower, respectively, than degradation. G. Same as above but for the phase portraits of total–new RNA planes. H. Strong linearity in the new–total RNA phase planes of *HMGB2* and *HMGA2*, with ascending slope *k* for longer labeling time periods. I. Strong linearity between −*log*(1 − *k*) and labeling time period *t* for *HMGB2* and *HMGA2*. F-I all used the kinetics experiment dataset from scEU-seq study (Battich et al. 2020)

When only cscRNA-seq data are available, or when one needs to use splicing data from tscRNA-seq experiments (which also capture intrinsic splicing kinetics), ***dynamo*** can be used to estimate the relative degradation rate constant 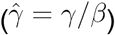 and relative spliced RNA velocity (**Fig. 2B, top**). The estimation methods built upon **Model 1** from **Fig. SI2A** include both the original method (La Manno et al. 2018) and the generalized method of moments (GMM). The GMM, in turn, consists of the stochastic splicing method, which relies on a master equation formulation of RNA kinetics (see **MATERIAL AND METHODS**) and is equivalent to the stochastic method developed recently (Bergen et al. 2020), and a new approach, the negative binomial (NB) method, which additionally models the gene expression at steady state as a negative binomial (NB) distribution, in the same vein as in previous studies (Grün, Kester, and van Oudenaarden 2014).

By comparison, from a tscRNA-seq experiment, one can estimate the absolute kinetic parameters (*α, β, γ*) and calculate unspliced, spliced, new, or total RNA velocity, depending on the exact labeling strategy and underlying model used (**Fig. 2B, bottom**). We suggest three general labeling strategies, namely one-shot, kinetics/pulse, and degradation/chase experiments, which cover most labeling methods in published tscRNA-seq experiments, aimed at estimating different RNA kinetic parameters (**Fig. 2C**). It is possible to extend or combine these three general labeling strategies to more complicated labeling schemes, e.g., the fourth type in (**Fig. 2C**), which consists of a time-series of multiple kinetics experiments, or a mixture experiment as in scEU-seq study (Battich et al. 2020), in which a variable initial kinetics experiment and later accompanying degradation experiment are conducted but kept in a fixed time period.

Estimating the parameters and RNA velocities with labeling data involves some technical subtleties, which we took into account when developing the corresponding algorithms tailored for each labeling strategy. Overall, we estimate absolute splicing and degradation constants (*β, γ*) by first estimating the degradation rates from labeling data and then the scaled degradation rate constant 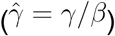 from splicing data, followed by obtaining a confident splicing rate constant 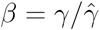 (See **MATERIAL AND METHODS** and **SUPPLEMENTARY METHODS** for details).

To demonstrate the effectiveness of our approach, we applied our framework to two previously reported datasets: a degradation dataset obtained by scNT-seq of murine ESCs (Q. Qiu et al. 2020) and a kinetics dataset obtained by scEU-seq of RPE-1 cells (Battich et al. 2020) (**Fig. 2D–I and Fig. SI2B–I**). In both datasets, the values of *γ* estimated from the degradation experiment, or those from the kinetics experiment using the two-step method (see **MATERIAL AND METHODS**), show no apparent correlation with 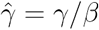. This indicates that the universal splicing rate *β* for all genes, as assumed in the original velocity method, does not generally hold (**Fig. 2D, left** and **middle**). Unsurprisingly, the splicing rates are generally much higher than the degradation rates, indicating that splicing kinetics are faster overall than degradation kinetics (**Fig. SI2B left** and **middle, C**). Still, certain genes have extremely fast degradation rates (**Fig. SI2B left** and **middle, C**). For example, *Slc25a32* degrades quickly, with a half life (*t*_1/2_ = ln 2/*γ*) of just 14 minutes, 81 times faster than *Ank2* (*t*_1/2_ of 18.6 hours) (**Fig. SI2D**). Housekeeping genes tend to be spliced quickly and degraded more slowly than other genes (**Fig. SI2E**).

In the scEU-seq cell-cycle data (Battich et al. 2020), genes with either fast splicing or fast degradation rates were enriched in cell-cycle–related pathways (**Fig. SI2F**). Interestingly, splicing and degradation rates of mouse genes are correlated with, but generally higher than, those of their human orthologs (**Fig. 2D right, Fig. SI 2B right**). This result complements the conclusion of (Matsuda et al. 2020; Rayon et al. 2020), which was based on a small set of human proteins (Battich et al. 2020) that have half-lives roughly twice as long as their mouse counterparts. Log-likelihood and R-square values (see **MATERIAL AND METHODS**) calculated based on observed data and estimated values from the curve-fitting and linear regression, respectively, confirm that parameter estimations confidently fit the data for the degradation and kinetics experiments (**Fig. 2E, H, I and Fig. SI2D**). In particular, the new and total RNAs show the expected strong linear relationship, with slope increasing with the labeling time during the kinetics experiment (**Fig. 2E, G–I;** see also **MATERIAL AND METHODS**). Analysis of the transcription and degradation rates for the mixture experiment (Battich et al. 2020) (**Fig. SI2G–I**) revealed that the genes with the highest transcription rates are all mitochondrially encoded (*ATP8, MT-RNR2, ND2, TRMT2B*) (**Fig. SI2H**), likely reflecting a high demand for energy production and the presence of multiple copies of the mitochondrial genome in a single cell.

The spliced RNA velocity reflects splicing and degradation kinetics, whereas labeling-based total RNA velocity reflects transcription and degradation kinetics (**Fig. 2B**, bottom; see also **MATERIAL AND METHODS**). Thus, for a kinetics experiment, we can plot the unspliced/spliced velocity on the “phase plane” (La Manno et al. 2018) of spliced and unspliced RNAs, as well as the new/total velocity on the “phase plane” of total and new RNAs. For example, the splicing rate of *HMGB2* is greater than its degradation rate (**Fig. 2F, G**), and across cells its unspliced RNA is less abundant than its spliced RNA (**Fig. 2F, G**). By contrast, *HMGA2* exhibits the opposite dynamic: Spliced RNA velocities of both genes are close to zero near the line fitted by the steady-state model, and positive above the line (**Fig. 2F**). The new RNA velocities are always non-negative, as the levels of labeled RNAs generally increase during a short labeling experiment. In comparison, total RNA velocities do not exhibit any particular structure on the total-new RNA phase plane (**Fig. 2G**). Thus, these results reveal the relationship between the ratio of unspliced and spliced RNA levels and the ratio of the splicing and degradation rates, as well as some subtleties distinguishing different types of velocities in the phase plane.

The analysis described above illustrates the generality of our inclusive estimation framework across multiple platforms, labeling strategies and biological systems, which also reconciles the programmable labeling kinetics and intrinsic splicing kinetics to achieve accurate quantification of RNA synthesis and turnover rates, enabling functional biological discoveries.

### RNA Metabolic Labeling Improves Accuracy of RNA Velocity Estimation

scSLAM-seq studies demonstrated that NTR-velocity outperforms splicing-based RNA velocity in revealing the directionality of velocity (Erhard et al. 2019). Reanalysis of those data with ***dynamo*** yielded good separation of CMV-infected and mock murine fibroblast cells in the UMAP space (**Fig. 3A**). To determine which data type offers more consistent vector flow, and is therefore numerically more stable for vector field reconstruction, we use the cosine correlation between the velocity vectors of each individual cell and those of its nearest neighbors in the PCA space to quantify velocity vector consistency (**Fig. 3B**). Considering only the scSLAM-seq data, velocity vectors were more consistent when using labeling data than when using the corresponding splicing data, implying that unbiased measurement of new RNA by RNA metabolic labeling is a superior approach because it does not depend on the “mis-priming” of introns in conventional scRNA-seq (**Fig. 3B**). Interestingly, the velocity flow from droplet based scRNAseq with 10x Chromium platform was even more consistent than that from the labeling data of the scSLAM-seq experiment. Downsampling suggested that this result arose due to the improved statistical power gained by using a much larger number of cells (~1000 cells in 10x vs. ~ 100 cells in the splicing data), and because the PCR-duplicates-removed unique molecular identifier (UMI) counts instead of read counts enabled by the 10x Chromium platform (**Fig. 3B**). These analyses suggest that moving forward large-scale, UMI-based metabolic labeling datasets will be optimal for velocity analysis and vector field reconstruction.

**Figure 3:**
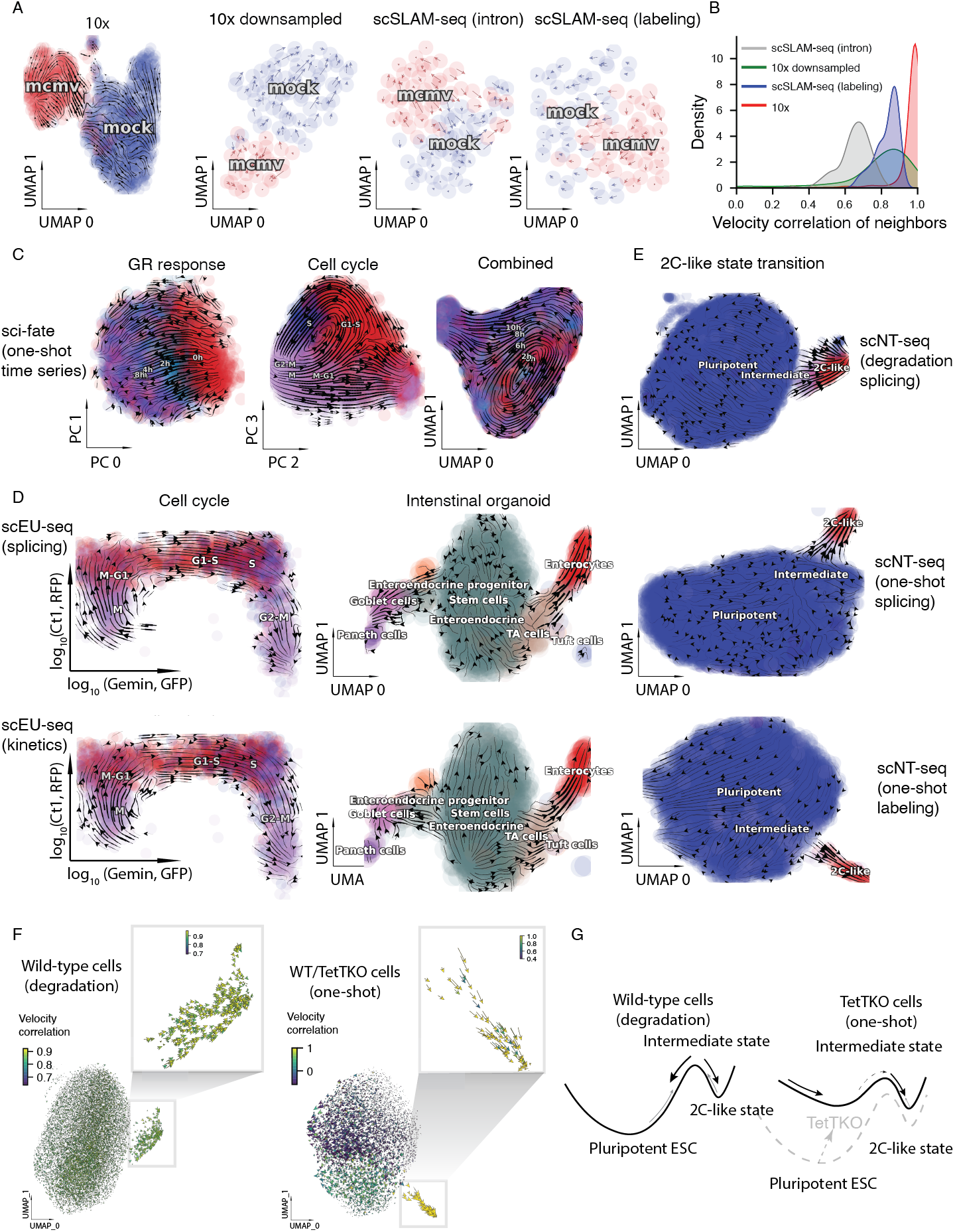
Metabolic Labeling Experiments Improve and Generalize RNA Velocity Estimation. A. RNA velocity streamline plots on the UMAP space for conventional and **one-shot** labelingbased scRNA-seq datasets from mouse fibroblast cells of the 2-h CMV (cytomegalovirus) infection scSLAM-seq experiment. From left to right: streamline plots of the full 10x dataset, the downsampled 10x dataset [same cell number (96) as the rest two panels], the intronexon dataset from the scSLAM-seq experiment, and the labeling based dataset from the scSLAM-seq experiment. Cells are colored by status of infection (mcmv: infected by MCMV; mock: not infected). B. Density plot of velocity correlation of neighbor cells, indicative of local velocity consistency, across the four datasets shown in **A**. Unbiased nascent RNA labeling, and large-scale, UMI-based scRNA-seq improves local RNA velocity consistency. C. RNA velocity streamline plots of **one-shot** labeling dataset from (Cao, Zhou, et al. 2020) reveal two orthogonal processes: GR response and cell cycle for the dexamethasone treated A549 cells. From left to right: streamline plot on the first two principal components (PCs), the second two PCs, and the first two UMAP components that are reduced from the four PCs, respectively. D. Conventional (**top**) and **kinetics** labeling (**bottom**) RNA velocity analysis of the RPE1-FUCCI cells (**left**) and murine intestinal organoid system (**right**) of the scEU-seq study. E. Conventional (**top**, **middle**) and **degradation** labeling (**bottom**) RNA velocity analysis of the TET-dependent stepwise pluripotent–2C bidirectional transition of murine ESC in the scNT-seq study. F. Cells committed to 2C-like totipotent states showing strong local velocity consistency. Top right boxes are magnified versions of the corresponding insets in the bottom. G. A bistable switch model of ESC pluripotent and 2C-like totipotent cell states explains the increased commitment of TetTKO cells to the 2C-like totipotent state cells through the intermediate cell state. Y-axis corresponds to the metaphorical “potential” (the global stability) of cell states.

To assess ***dynamo***’s ability to deconvolve orthogonal cellular processes, we analyzed datasets from sci-fate in which cell cycle progression and glucocorticoid receptor activation are explored (Cao, Zhou, et al. 2020). In that study, Cao *et. al* also proposed an interesting linkage analysis (See **DISCUSSION**) in which they observed the cell-cycle transition from G1 phase to S phase, and then the transition to G2/M phase, as well as the transition from no GR (glucocorticoid receptor) response to a high–GR activity state, whereas they did not observe such patterns from either splicing-based or NTR-based RNA velocity analysis (Cao, Zhou, et al. 2020). We revisited those transitions and performed time-resolved total RNA velocity analysis on combined or individual set(s) of cell-cycle and GR response genes detected by the original study. A streamline plot of the first two PCA components of the combined analysis (or the UMAP space of the separated analysis with only GR-response-related genes) revealed a smooth sequential transition from untreated cells at time point 0 to 2, 4, 6, and 8 hours after the initial DEX (dexamethasone) treatment, indicating progressive activation of the GR response (**Fig. 3C GR response, Fig. SI3A**). Similarly, an acyclic loop matching the cell-cycle progression from G1 to S, from S to G2M, and from M to G1 was revealed from the second two PCA components of the combined analysis (or the UMAP space of the separated analysis with only cell-cycle-related genes) (**Fig. 3C Cell cycle, Fig. SI3A**). Interestingly, when projecting cells onto the UMAP space, combined analysis revealed both a linear progression of the GR response and a circular loop indicative of cell cycle (**Fig. 3C combined)**. Next, we analyzed datasets from the scEU-seq study (Battich et al. 2020) and observed a sequential transition from M to M-G1, G1-S, S, G2-M for the RPE1-FUCCI cells (shown from left to right in **Fig. 3D, left column**) as well as a bifurcation (**Fig. 3D right column**) from intestinal stem cells into the secretory lineage of enteroendocrine, goblet, and paneth cells (**left**) and the enterocyte lineage of TA, tuft and enterocyte cells (**right**) for the intestinal organoid data.

Next, we investigated the mESC dataset from the scNT-seq study (Q. Qiu et al. 2020). We first analyzed the mESC degradation experiment and detected a transition from the intermediate cell state into either the dominant pluripotent ESCs or rare 2C-like totipotent cells that manifest as a tip in the UMAP embedding from our velocity results (**Fig. 3E top**). We then analyzed the *Tet 1/2/3* triple knockout (TetTKO) dataset. Both splicing-based and labeling-based velocity revealed an outgoing flow from the pluripotent ESCs, with a considerable fraction of cells moving towards the intermediate and ultimately the 2C-like totipotent cell state in the tip (**Fig. 3E middle** and **bottom, F, Fig. SI3B, C**). To unify these results, we proposed a bistable switch model consisting of ESC pluripotent and 2C-like totipotent cell states (**Fig. 3F, G**). In wild-type cells, the intermediate cell state is an unstable transition state (saddle point) that can automatically transition into the totipotent state or the pluripotent state, the latter of which occupies the predominant attractor basin. On the other hand, in TetTKO cells, pluripotent cells are perturbed, lowering the barrier between the pluripotent and intermediate cell states and easing the transition from pluripotent to totipotent cells.

In summary, these findings reveal an optimal experimental strategy for recovering RNA velocity information and demonstrate the generalizability and accuracy of our approach in resolving the time-resolved RNA velocity across various single-cell protocols, labeling strategies, and biological systems. A similar analysis can be also applied to cscRNA-seq datasets for robust RNA velocity analysis (**Fig. SI3D–H**).

### Robust Reconstruction of Vector Field Functions of Single Cells

Proceeding from improved single-cell velocity estimates, we will now describe how to go beyond discrete, sparse, and local measures of velocity samples to continuous vector field functions in the full gene expression state space. We start with a theoretical discussion of the recoverability of vector field functions (See **Fig. SI4A** and **SUPPLEMENTARY METHODS**) ((Y Kim et al. 2000; Weinreb et al. 2018). Successful reconstruction of the vector field function from transcriptomic data depends on whether the input datasets capture sufficient dynamical information and whether hidden variables such as proteomic and epigenetic states are redundant in specifying cell dynamics. To test this, we examined a dataset (Kimmerling et al. 2016) in which sisters/cousins from primary activated murine CD8+ T cells were captured and measured using a specifically designed microfluidics platform (Fig. SI4B). Because sister or cousin cells are generated from the same cell through one or two cell divisions, respectively, they should explore the expression space in a similar manner (Fig. SI4C). Indeed, the transcriptomic distances between sisters and cousins are both significantly lower than those of random cell pairs (Fig. **SI**4D). Moreover, the distances between transcriptome-wide spliced RNA states of cells are highly correlated with those of estimated RNA velocity, and even more so for the unspliced RNA states (**Fig. SI4E**). In addition, cells close in transcriptome state shared similar RNA velocity vectors, and neighbor cells that also happened to be sisters or cousins did not exhibit higher similarity (**Fig. SI4F**). These results indicate when hidden variable effects are not apparent in the system, as in this case, one may predict velocity ***υ*** via a vector field function ***f*** once the the transcriptomic state *x* is known, namely, ***υ*** = ***f***(***x***).

To construct the vector field function (**Fig. 1A**), we adopted a machine learning approach that takes advantage of recent advances in vector-valued function approximation to scalably, efficiently, and robustly learn the transcriptomic vector field (see **Box 2**) from noisy and sparse samples of single-cell states and velocity estimates (**Fig. 4A**). The framework employs sparseVFC (sparse approximation of Vector Field Consensus) (J. Ma et al. 2013), which uses a vector-valued kernel method built on RKHS (reproducing kernel Hilbert space) to learn the vector field, which is expressed analytically as a weighted linear combination of a set of vector-valued kernel basis functions (**Fig. 4A Output**). The learning process relies on sparse approximation to estimate the coefficients (weights) of a selected number of basis functions, each associated with a control point, that is often much smaller than the number of data points, without loss of accuracy (**Fig. 4A Output**). With sparse approximation, the vector field reconstruction scales linearly with the number of data points in both computational time and memory requirements (J. Ma et al. 2013), allowing accurate reconstruction of a high dimensional vector field, e.g. in 30 principal components (PCs), using only a few hundred control points for more than 100,000 cells in a matter of minutes with a modern workstation. To account for the noise and outliers of velocity measurements, sparseVFC relies on an **EM algorithm** to iteratively optimize the set of inliers as well as the optimized coefficient set for each basis function corresponding to each control point (**Fig. 4A**), further improving the robustness of vector field reconstruction. With the continuous vector field function, we also derived analytical formulas of its Jacobian, acceleration, curvature, divergence, curl, etc. Although we used the two-gene system (See **below**) to illustrate the vector field reconstruction procedure, our reconstruction works in either high-dimensional PCA space or lower space after nonlinear dimension reduction (like UMAP), or directly in the full gene-expression space. The vector field reconstructed in low-dimensional space can then be projected back to the full transcriptomic space for gene-specific velocity and differential geometry analyses.

**Figure 4.**
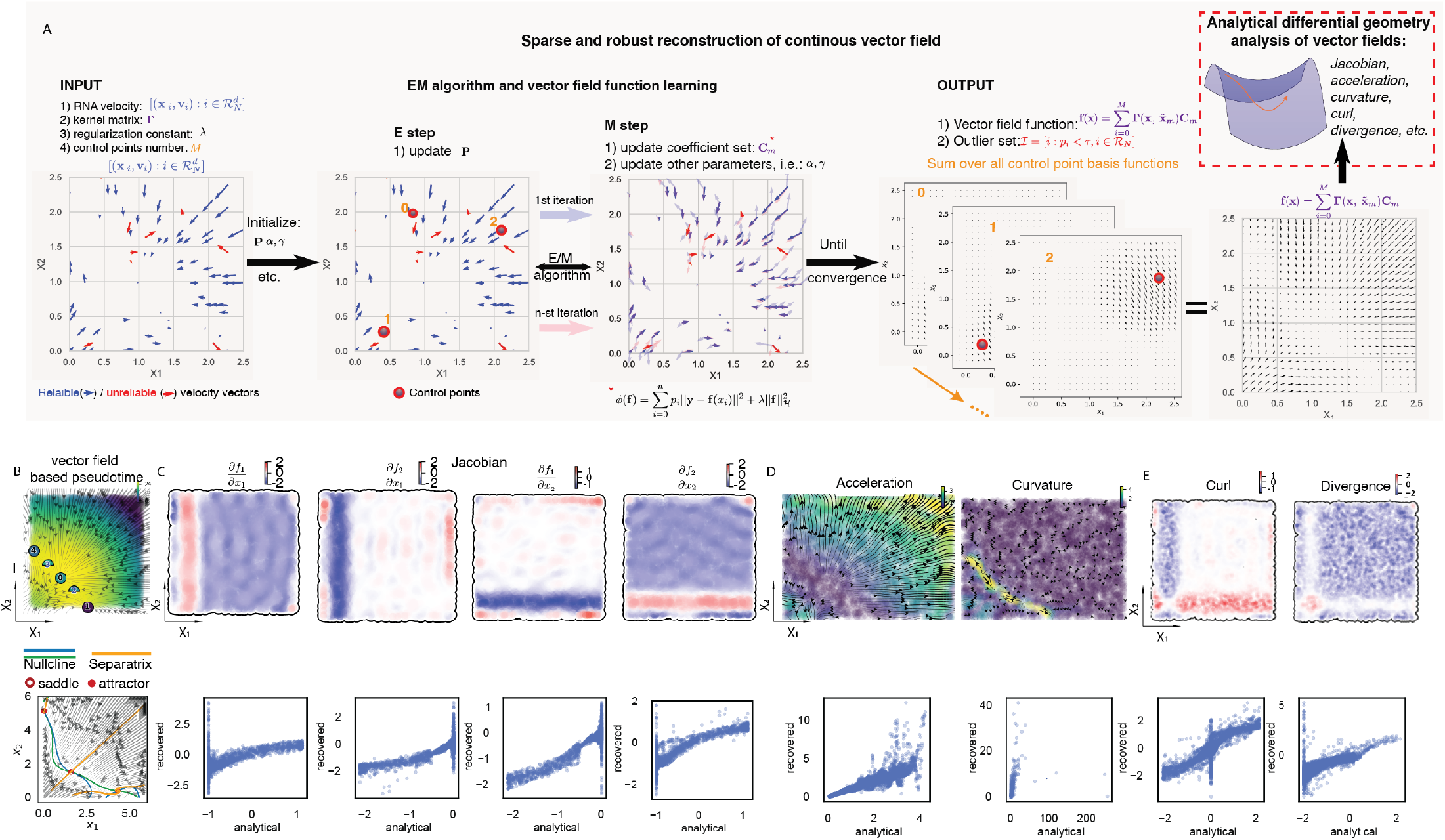
Mapping the Transcriptomic Vector Field, Quantifying its Topography, and Moving towards Differential Geometry of Single Cells. A. Functional reconstruction of the continuous and analytical velocity vector field from sparse, noisy single cell velocity measurements with sparseVFC (J. Ma et al. 2013) (see details in **MATERIAL and METHODS** and **SUPPLEMENTARY METHODS**). B. Reconstructed vector field and topological features of the simulated toggle-switch system. **Top**: Scatterplots of cells (x/y-axis: expression of *x*_1_/*x*_2_) that are colored by vector-field based pseudotime, calculated via Hodge decomposition on simplicial complexes (a sparse directional graph) constructed based on the learned vector field function (Maehara and Ohkawa 2019). Full cycle nodes correspond to attractors while half-cycle saddle points. Streamline plot of the reconstructed vector field is superimposed on top of the scatterplot of cells. **Bottom**: x/y-nucline and separatrix, plotted on top the streamline plot of the reconstructed vector field. C. Scatterplots of cells (x/y-axis: expression of *x*_1_/*x*_2_) with a frontier representing the expression boundary of sample cells (**top**). Cells are colored by the estimated values of the indicated Jacobian elements. Scatterplots of the estimated values of indicated Jacobian elements and corresponding analytical values across cells. In general, recovered Jacobian closely matches the ground truth but deteriorates at the boundary of the sample data points. D. Same as in **C** but for the recovered acceleration and curvature. Since acceleration and curvature are vectors, the streamlines of the recovered acceleration and curvature vector field are visualized. Cells are colored by the length of acceleration or curvature vectors across cells. E. Same as in **C** but for the recovered curl and curvature.

#### Box 2: Vector Field Function Learning in Reproducing Kernel Hilbert Space

The overall goal of vector field function learning is to find a vector-valued function ***f*** in the function space 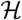 consisting of all possible vector field functions, such that, trained by a sparse set of coordinate–velocity data pairs 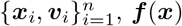 gives the velocity ***υ*** at an arbitrary coordinate ***x*** as schemed in **Box Fig. 2A**.

**Box Fig. 2.**
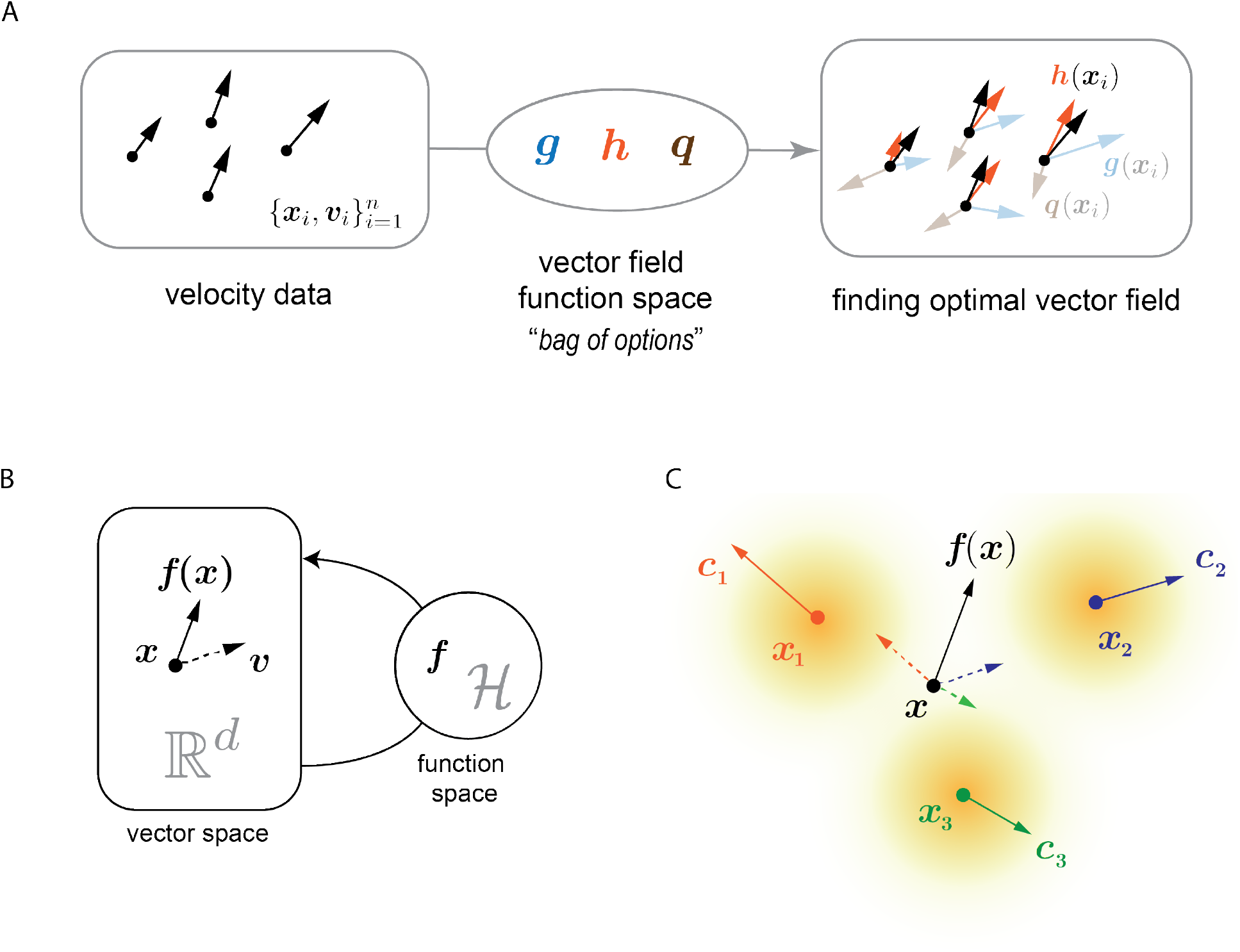
Learning vector field function that is expressed a summation of a set of basis functions in the function space.

The coordinates ***x**_i_* in the gene expression space are fed into vector field functions (***g, h***, and ***q***) in the function space 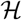, which output a vector, also in the gene expression space, for each coordinate. To distinguish the output vectors from the velocity vectors from the data, these vectors from the vector field functions are called “*evaluations*”. As shown in the rightmost panel in **Box Fig. 2A**, intuitively ***h*** is best when one compares its evaluations ***h***(***x**_i_*) to the velocity data ***υ**_i_*. This comparison can be formally evaluated with a loss *functional* (a function of functions) Φ(***f***) that measures how close the evaluations of the vector field functions and the velocity data are.

In general, a function space may have an infinite number of functions, and the learning procedure involves singling out one function through variational analysis,

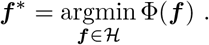

One example of the loss functional assumes a sum-of-squares form with a regularization term to reduce the risk of overfitting,

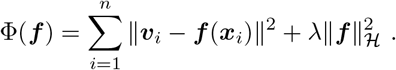

The last *L*_2_ regularization term is an abstractly defined inner product of ***f*** in the function space 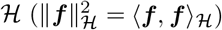. It is nontrivial to minimize the above loss functional computationally with respect to functions in the function space 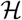. Note that ***f*** is an object defined in the function space, whereas ***f***(***x***), the evaluation of ***f*** at point ***x***, is an object in the gene expression space, an 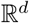 vector space, the same space in which the velocity vectors ***υ*** lives. The diagram in **Box Fig. 2B** outlines the relationships of the vector space and the function space. We need a mathematical tool to 1) represent the vector field function ***f*** in a way that its evaluation ***f***(***x***) can be computed analytically, so that the loss function minimization can be performed computationally; and 2) compute 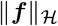 without actually calculating the inner product in the function space, as this is often hard to achieve.

One such tool is the reproducing kernel Hilbert space (RKHS)· “Hilbert space” is effectively the function space (see **MATERIAL AND METHODS**), and the “reproducing kernel” is a function that takes in two coordinates from the 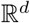 vector space and outputs a *d* × *d* matrix. A key elegant feature of the reproducing kernel is that it both encodes all the “options” in the function space and determines how the functional inner product is performed.

In RKHS, one evaluation of the vector field function ***f*** is given by,

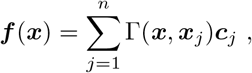

where ***c*** is a combination coefficient vector in the 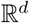 space, and Γ(·, ·) the reproducing kernel. A simple but effective choice of the reproducing kernel is the Gaussian kernel:

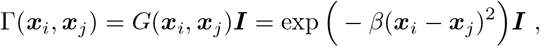

where ***I*** is the identity matrix, and *β* a width parameter of the Gaussian function *G*(·, ·). This reduces the representation to a simpler form:

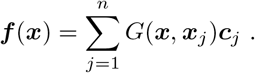

Because *G*(***x, x***_*j*_) is only a scalar, the evaluation becomes a superposition of coefficient vectors ***c***_*j*_’s, whose weights decay as their distances to the point of interest ***x*** increase, as shown in the **Box Fig. 2C**. The three data points each have an associated coefficient vector, and the evaluation at ***x*** is a combination of them, weighted by the Gaussian kernel (yellow radial gradient).

Also, the functional inner product in RKHS can be computed as:

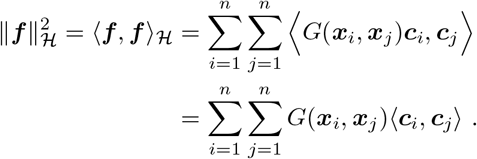

Note that the inner product 〈·, ·〉 without the subscript is defined on the 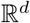 vector space, and a common choice is the dot product, i.e. 〈***u, v***〉 = ***u***^T^ ***υ***. With both the functional form of the vector field and its norm explicitly defined, the loss function can be optimized with respect to the coefficient vectors ***c***. For further steps involved in minimizing the loss function, see the corresponding sections in **MATERIAL AND METHODS** and **SUPPLEMENTARY METHODS**.

To demonstrate the power of the vector field reconstruction, we first tested the efficacy of our reconstruction on a simulation dataset with 5000 randomly sampled points on the state space of the toggle-switch model introduced in **Fig. 1**. The estimated streamlines of the reconstructed vector field, as well as the fixed points, nullcline, etc., were nearly indistinguishable from the analytical ones (**Fig. 4B**). Moreover, we could accurately recover the Jacobian matrix across the state space, although the agreement deteriorated at boundary regions with far fewer sample data points (**Fig. 4C**). The estimated higher-order vector calculus quantities closely matched the true analytically computed quantities, but with diminishing consistency (**Fig. 4D, E, Fig. SI4G**). The analytical formulae of vector calculus that we derived provide a tremendous increase in speed, enabling analysis that was nearly 1000x faster than highly-optimized state-of-the-art numerical approaches such as [numdifftools](https://pypi.org/project/numdifftools/) (**Fig. SI4H**).

Pseudotime analysis has been routinely used to recover a sequence of events from single-cell genomics data, e.g., how progenitor cells differentiate into a terminal cell state (**Fig. SI4D**). Because a vector field function can be decomposed into a gradient or curl part, one can use the gradient part to define a scalar potential (canonically defined only for closed physical systems) even if the vector field system is open, e.g., a biological system (Ao 2009). Hence, we tested the idea of using the scalar potential from a reconstructed vector field through the Hodge decomposition as a new type of pseudotime analysis based on a vector field (Maehara and Ohkawa 2019). Because this method utilizes velocity fields that consist of the direction and magnitude of cell dynamics, it is intrinsically directional and arguably more relevant to real time than other pseudotime methods. As expected, the vector field–based pseudotime revealed a smooth cell state transition from states far from attractor states (**Fig. 4B bottom**). Similar analysis for the wild-type and TetTKO murine ESCs further confirmed an automatic transition from the intermediate cell states to pluripotent and 2C-like totipotent cell states, as well as an increased transition rate from pluripotent cells to the intermediate and 2C-like totipotent cell states in TetTKO cells (**Fig. SI4I**). We further performed vector field reconstruction analysis on a variety of published cscRNA-seq datasets, demonstrating the robustness, generalizability of our vector field reconstruction as well as the downstream differential geometry analyses (**Fig. SI4J–N**).

### Vector Field Trajectory Predictions are Concordant with Sequential Clone Tracing of HL60 Neutrophil Lineage Commitment and Murine Hematopoiesis

Once a vector field is learned, one immediate application is to predict the historical or future state of a cell in a manner analogous to Newtonian mechanics, i.e., if one knows the position of a cell in the state space and the function of the state evolution, one can predict where it is and how fast it is moving through the state space, here the gene expression space, at any point in time (**SUPPLEMENTARY ANIMATION**). We tested the accuracy and limitation of this prediction by comparing the single-cell trajectory prediction with gene expression in clonal cells (cells arising from the same progenitor through cell division) measured sequentially, which approximates the dynamics of a single cell over time (**Fig. 5A**).

**Figure 5.**
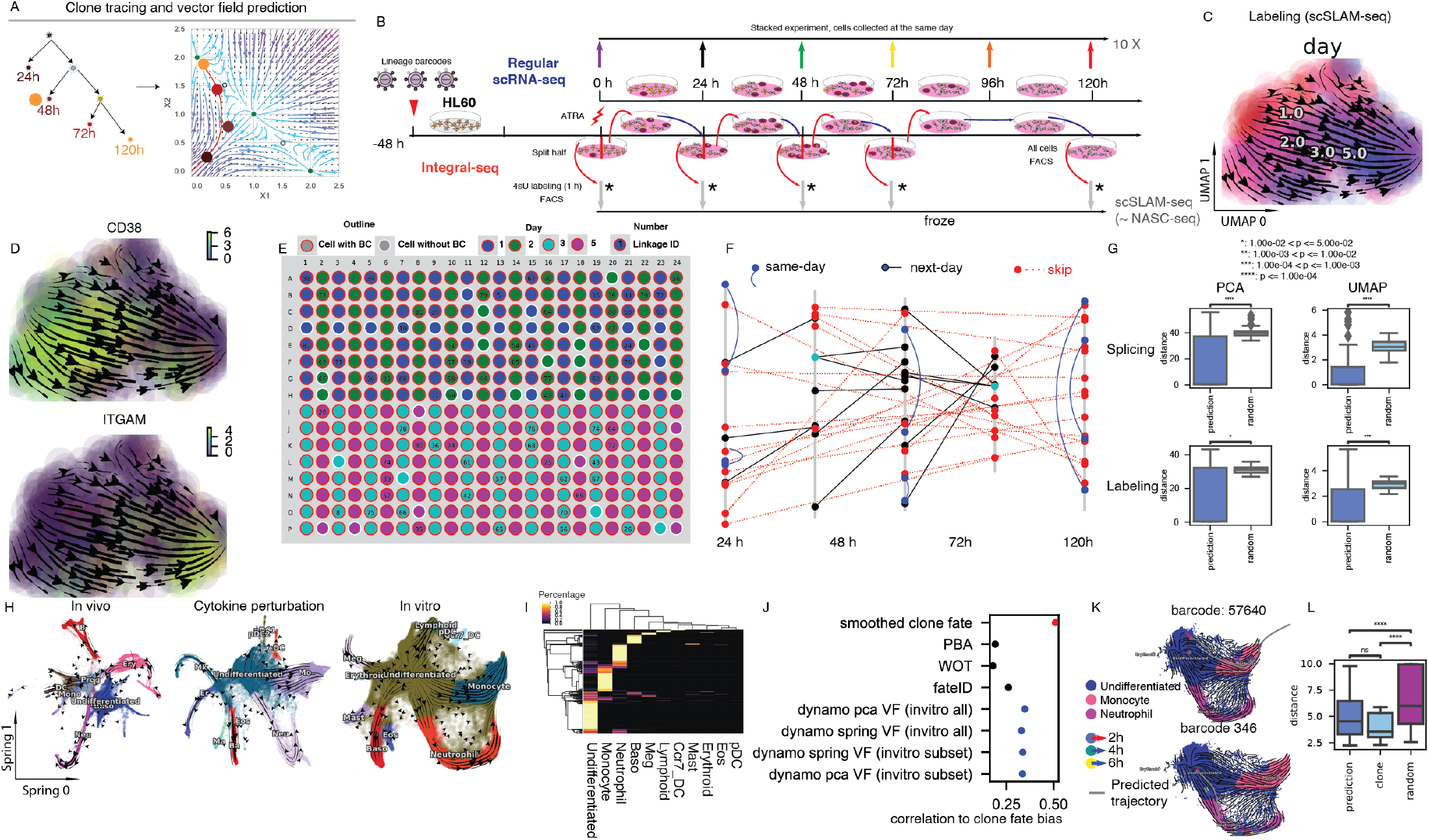
Vector Field–based Cell Fate Predictions are Validated by Clonally related Single Cells. A. Clonity of cells, which were sequentially sampled at different time points, is inferred based on the static barcodes (**left**) and is used to validate the vector field prediction (the red line, **right**). B. Experimental schemes of conventional 10x Chromium–based scRNA-seq (**top**) and plate-based metabolic labeling scRNA-seq (scSLAM-seq or NASC-seq) coupled with sequential clonal cell tracking via lentivirus lineage barcodes (**bottom**) for neutrophil fate commitment of HL60 cells under ATRA treatment. C. RNA velocity streamline plot of UMAP embedding (same as in **D**) based on labeling data from the scRNA-seq experiment reveals neutrophil-lineage commitment over time. D. Progenitor marker (*CD38*) and neutrophil marker (*CD11b* or *ITGAM*) expression on UMAP space and streamline plot confirm the directionality of neutrophil lineage differentiation. E. Layout of an example 384-well plate and locations of clonally related cells. F. Forty confident clone-related linkages across different days among 944 cells from the integral-seq experiment. G. Boxplot of minimal distance of clone cells in later time points to the vector field prediction trajectory and the distance of random cells from the same day. Mann–Whitney–Wilcoxon test (two-sided) was used to calculate the p-value between groups. ****: *p* < = 1*e* – 04. H. RNA velocity streamline plot of SPRING embedding (same as in **K**, embedding is from (Weinreb et al. 2020)) reveals lineage hierarchy from murine hematopoietic stem cells to myeloid (megakaryocytes, erythroids, mast cells, basophil, eosphil, neutrophil, monocytes, dendritic cells, etc.) and lymphoid lineages for *in vivo* and *in vitro* systems. Cells are colored by the cell type identity. I. A majority of cells from the same clone are biased towards a specific lineage or remain in the undifferentiated cell state. In the heatmap, row, column, and color correspond to a particular clone of cells, a particular cell lineage, and the probability that cells eventually commit to a cell lineage (or maintain the undifferentiated cell state). J. Comparing vector field–based lineage fate predictions with other state-of-art methods. Smoothed clone fate (red dot) is the prediction based on clone barcodes. PBA, WOT, and fateID (black dots) are predictions based on other state-of-art algorithms that do not use velocity information. Those first four methods are from the original study (Weinreb et al. 2020). Predictions for all cells (fifth and sixth items) or the same subset of cells (seventh and eighth items) from the original studies are presented. Both PCA embedding, and SPRING embedding from the vectors fields based on the original study were used for prediction. K. Predictions with vector fields tolerate noise from estimated single-cell velocities. L. Boxplot of minimal distances of day 6 clone cells to the vector field prediction trajectory, the distance between cells from the same clones, and the distances of random cells from day 6. Same statistical test is used as in **G**.

To validate single-cell trajectory prediction, we integrated metabolic labeling based scRNA-seq and sequential capture of uniquely barcoded clone cells over time (**Fig. 5B**). We applied this strategy to study ATRA (all-trans-retinoic acid)-induced neutrophil lineage commitment of a human leukemia cell line (HL60). To provide a baseline for the differentiation protocol and the vector field reconstruction, we also performed a 10x scRNA-seq experiment in which differentiation was initialized at different days, so that we could harvest all samples on the same day (**Fig. 5B**). For the tscRNA-seq experiment, we infected cells with the GBC (gene barcode) library originally designed for Perturb-seq (Adamson et al. 2016), so that randomly synthesized barcode sequences were genomically integrated to barcode cells and each of their descendants. Importantly, the DNA barcodes are transcribed so they can be measured during scRNA-seq. After infection, cells were rested for 3 days before the neutrophil differentiation process was initialized by administration of ATRA. On days 0, 1, 2, 3, and 5, we split the cells into two halves: one half was subjected to 4sU labeling, followed by flow sorting of cells with barcodes expressing the blue fluorescent protein (BFP) reporter into 96-well plates; the other half continued differentiation until the next round of splitting or were collected at the final time point for sorting. We then used a protocol modified from scSLAM-seq/NASC-seq (Erhard et al. 2019; Hendriks et al. 2019) to obtain a tscRNA-seq dataset (see **MATERIAL AND METHODS**).

Both of the splicing-based RNA velocities from the 10x and clone-traced tscRNA-seq dataset, as well as the labeling-based RNA velocity from the same tscRNA-seq experiment, revealed a smooth transition from day 1 to day 5, although as expected it was less smooth in the spliced-based RNA velocity from the tscRNA-seq experiment (**Fig. 5C, Fig. SI5A, B**). FACS analysis of CD11b (*ITGAM*) and CD14 revealed that at day 1, > 6% of cells were already CD11b+, whereas more than 83% cells were CD11b+ at day 5, indicating that ATRA treatment induces differentiation of HL60 cells with high efficacy (**Fig. SI5C**). The neutrophil markers *ITGAM* or *Fut4* were progressively turned on after ATRA treatment, whereas the progenitor markers *CD38, FGR*, and *LCP2* were turned off (**Fig. 5D, Fig. SI5A-D**).

We used customized scripts to process the clone barcodes in single cells, first assigning each cell to a set of clone barcodes, next removing spurious clonal linkages processed at nearby wells on the plate (**Fig. 5E**), and finally building a “cell linkage” graph (**Fig. 5F**). In the end, we obtained 41 pairs of linkages among 944 cells, of which 6 were same-day linkages (cells that are clonally related but appears from the day), 14 were next-day linkages (cells that are clonally related but appear from two consecutive days) and 21 were skip-day linkages (cells that are clonally related but appear from two different days that are more than one day apart) (**Fig. 5F**). Next, we took cells at earlier time points from confident next-day or skip-day linkages and predicted their cell states over time in either the spliced- or labeling-based vector field. We then calculated the minimal distance (prediction distance) from clone cells at later time points to the vector field–predicted trajectory, and the distance between random cell pairs on the corresponding day (random-cell distance). We found that the predicted distances were significantly smaller than the random-cell distance (**Fig. 5G**). These results indicated that our vector field could predict cell trajectory reasonably well over several days in the HL60 neutrophil differentiation system.

To further test the general applicability and potential limitations of our vector field trajectory predictions, we applied our method to data from a recently published study in which the clonal fate of barcoded hematopoietic stem cells (HSCs) was tracked by sequential profiling of the cell population over time (Weinreb et al. 2020). This study includes three major experiments: *in vitro*, cytokine perturbation, and *in vivo* studies (See **SUPPLEMENTARY METHODS**). For each experiment, although roughly 100,000 cells were sequenced, the sequencing depth was shallow (only 600 genes were captured per cell on average). To improve RNA velocity estimation for such shallowly sequenced datasets and correct unexpected backward flow, we developed a heuristic approach that relies on a gene-wise velocity confidence metric based on the pattern of a gene in the phase plane according to broad cell type hierarchy priors [(**Fig. SI5E**), see more discussion in **SI**]. After removal of genes with less confident velocity estimations, RNA velocity flow from ***dynamo*** revealed a smooth hierarchical transition from murine HSCs to each of the expected lineages in all three experiments (**Fig. 5H**). The erythroid lineage from the *in vivo* study was still mixed with the wrong backward flow, indicating possible batch effects between different animals or possible *in vivo* micro-environment differences across cells belonging to the same clone (i.e., hidden variables). We also applied the same correction approach to another study of endothelial to hematopoietic transition (Zhu et al. 2020), and found that it could readily correct the backflow from the pre-HE cell type to the HE bottleneck (**Fig. SI5E**).

We next used the corrected RNA velocity estimates to reconstruct the vector field and predict the fate of individual cells over a long period during hematopoiesis, and to compare the fate with the clone barcode information. After estimating the cell fate probability distribution of cells with the same clone barcodes on day 6, we found that the majority of clones were biased to a specific lineage, whereas a considerable number of clones remained in the undifferentiated state even at day 6 (**Fig. 5I**). Focusing on clones whose day 0 cells were undifferentiated but day 6 cells were differentiated, we numerically integrated the trajectories of these cells in the vector field based on either the PCA space, or the SPRING embedding from the original study. By comparing the distribution of fates assigned by the predicted single-cell trajectories (see **SUPPLEMENTARY METHOD**) to observed cells in each clone and the gold-standard clone barcode-based fate distribution, we found that our vector field–based prediction outperformed the methods used in the original study (PBA, WOT, and fateID) (**Fig. 5J**). Our prediction worked similarly well across vector fields based on SPRING or PCA space, and just the vector field based on only part of undifferentiated and the neutrophil/monocyte cells, as used in the original study or the full cell fates of the entire *in vitro* dataset (**Fig. 5J, Fig. SI5F**). However, the predictions were significantly worse for the cytokine dataset and even more so for the *in vivo* dataset (**Fig. SI5F**). These results suggested that leveraging both spliced and unspliced data, instead of merely total RNA information, allows better predictions of cell state dynamics than using only total RNA information, especially in conjunction with our vector field approach. In addition, perturbations by cytokines may overshadow intrinsic cell fate dynamics. Lastly, from the *in vivo* experiment, we concluded that the unmeasured environmental cues or other hidden variables that each cell experiences makes it questionable to assume a time-invariant vector field and to predict cell fates (See **SUPPLEMENTARY METHODS**).

To demonstrate the robustness of vector field trajectory prediction, we investigated the velocity vectors of two randomly chosen progenitors and its predicted trajectories for the *in vitro* data on the SPRING space. Interestingly, although the velocity vectors of those progenitors pointed in the wrong direction, our vector field prediction could still provide the correct cell fate predictions, as evidenced by the location of day 6 cells on the SPRING space (**Fig. 5K**). This may be because our vector field reconstruction relies on information from the entire dataset, and thus tolerates random noise from the input measurements. Lastly, the minimum distances from clone cells at later time points to the vector field–predicted trajectories were similar to the distances between cells from the same clones at later time points but significantly smaller than the distances of random cells from the same later time point, again confirming our ability to predict predicting single-cell trajectories with vector field functions reasonably well over several days (**Fig. 5L**).

Together, these results demonstrate that the vector field predictions can reliably predict the single cell fate trajectories over several days, allowing us to infer the life history and future fate of single cells from transcriptomic data.

### Vector Field Analysis Reveals Topological and Geometric Properties across a Variety of Dynamical Processes

Having demonstrated the validity of single-cell trajectory prediction of the vector field function, we next applied differential geometric analyses (**Fig. 4A, 6A**) to a variety of csc- or tscRNA-seq data to reveal predictive information about gene regulation underlying various biological processes. In general, we performed differential analyses and gene-set enrichment analyses based on top-ranked acceleration or curvature genes, as well as the top-ranked genes with the strongest self-interactions, top-ranked regulators/targets, or top-ranked interactions for each gene in individual cell types or across all cell types, with either raw or absolute values (**Fig. 6A**). Integrating that ranking information, we can build regulatory networks across different cell types, which can then be visualized with ArcPlot, CircosPlot, or other tools (See example of ArcPlot and CircosPlot in **Fig. 6A**).

**Figure 6:**
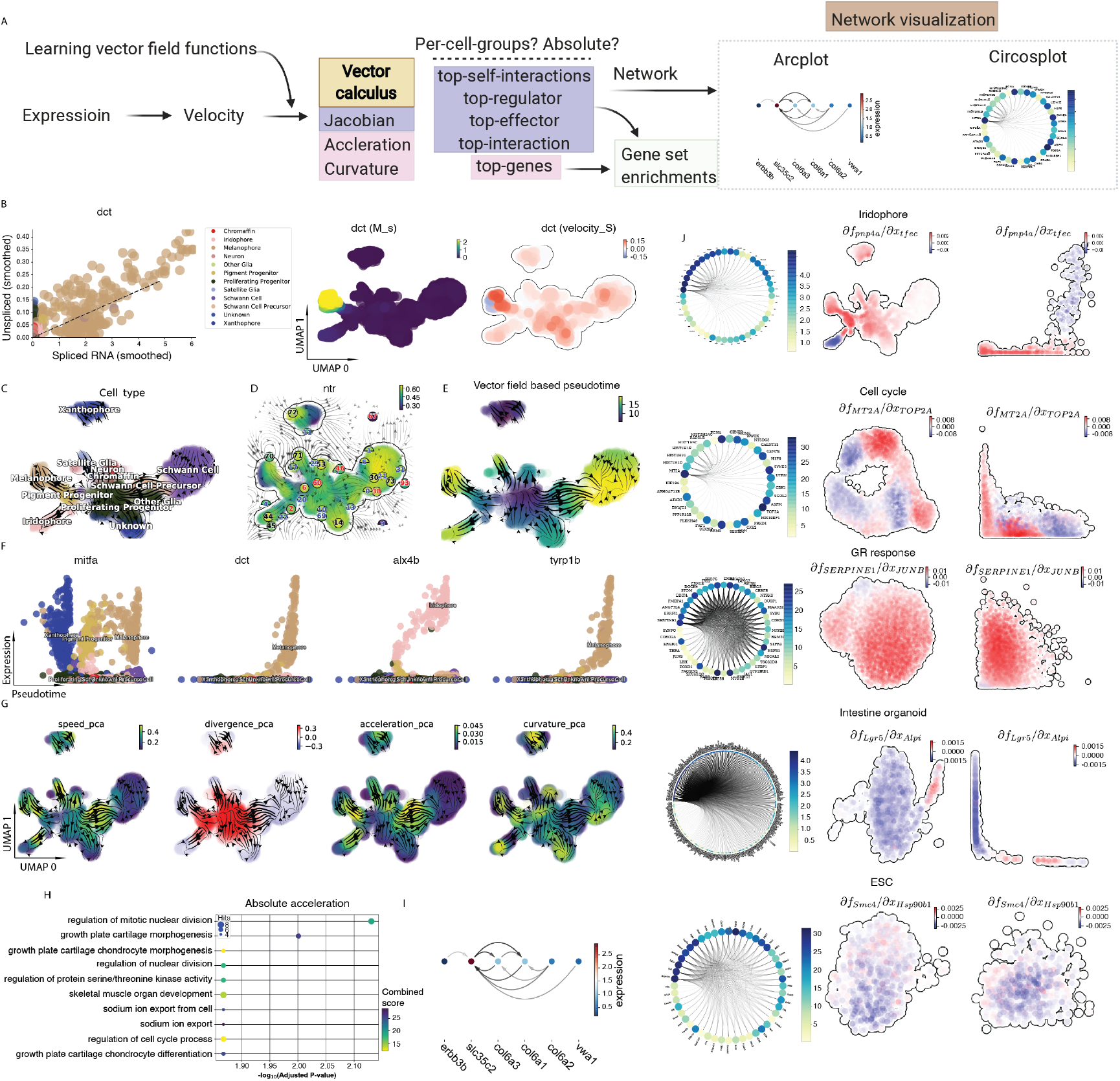
Vector Field and Differential Geometry Analysis of Zebrafish Pigmentation and other Dynamical Processes. A. Scheme for using Jacobian, acceleration, and curvature information to rank genes (either in raw or absolute values) across all cells or in each cell group, followed by gene set enrichment and regulatory network construction and visualization. B. Phase-portrait, conventional RNA velocity fitting (dashed line) of locally smoothed spliced and unspliced RNA (**left**), expression (**middle**), and velocity (**right**) scatterplots of the zebrafish melanophore marker *dct* on UMAP embedding (same as in **C–E, G, J**). C. Streamline plot of the estimated RNA velocity projected onto UMAP space reveals that multipotent pigment cell progenitors differentiate into pigment cells, peripheral neurons, Schwann cells, chromaffin cells, and other cell types. D. Reconstructed vector field and identified stable and unstable steady state points pinpoint the topological domains of progenitors, metastable states, and terminal cell types. The color of digits in each node is related to the type of fixed points (**Fig. 5C**): **black**: absorbing fixed points; **red**: emitting fixed points; **blue**: unstable fixed points. E. Pseudotime derived from the vector field function corroborates the arrow of time of zebrafish pigmentation. F Expression of the progenitor marker *mitfa* decreases from progenitors to terminal cell types, whereas expression of mature cell type marker, *dct* (melanophore), *tyrp1b* (melanophore), and *alx4b* (Iridophore) increases in the corresponding lineages with the pseudotime. G. Acceleration, curvature, divergence, and curl analyses reveal, respectively, key cell-fate commitment–related hotspots, decision points, sources, and sinks, as well as regions with processes orthogonal to cell differentiation (e.g., the cell cycle). H. GO enrichment analysis of genes with top absolute acceleration in a previously “unknown” cell cluster reveals its potential chondrocyte origin. I. Key gene regulatory network in the “unknown” cell cluster, derived from the estimated cellwise Jacobian matrices of chondrocyte related genes with the reconstructed vector field function. Network is visualized as an Arcplot. J. Jacobian matrix analysis is generally applicable to reveal context-specific gene regulatory networks across different cell types or technologies: from top to bottom, iriphore cells (cscRNA-seq), cell-cycle response (sci-fate), GR response (sci-fate), cell-cycle (scEU-seq), intestinal organoid (scEU-seq), ESC (scNT-seq). Circosplots of top gene interactions identified via Jacobian analysis for each datasets are shown on the **left**. Jacobian values between an example regulator and its target across cells on UMAP embedding (**middle**) or their gene expression space (x-axis: regulator, y-axis: effector, **right**) are also shown for each system.

For the main demonstration, we chose a high-quality cscRNA-seq dataset of zebrafish post-embryonic pigment cell differentiation because it involves a multitude of cell states and decision points (Saunders et al. 2019). The phase plane of the melanophore cell marker *dct* exhibited both high spliced and unspliced gene expression, as well as strong positive velocity only in melanophore cells (**Fig. 6B**). The RNA velocity on the UMAP embedding revealed a smooth transition from proliferating progenitors to pigment progenitors, which then ramified into various differentiated cell types (**Fig. 6C**).

To further analyze the cell fate transitions during pigment cell differentiation, we learned the vector field function (in the top 30 PCA or UMAP space), and then characterized the topological structure of the 2D UMAP vector field (**Fig. 6D**). We identified a number of fixed points with varying stability, with half circles indicating saddle points and full circles indicating stable fixed points (see **MATERIAL AND METHODS**). Node 6 is an emitting fixed point representing a destabilized progenitor state, whereas nodes 44, 70, 72, and 14 are absorbing (i.e., stable) fixed points corresponding respectively to the iridophore, melanophore, and xanthophore terminal cell types and a cell type not identified in the original study. Lastly, nodes 20 and 29 are unstable fixed points (saddle points) corresponding to the bifurcation point of the iridophore and melanophore lineages or that of the neuron and satellite glia lineages. The vector-field-based pseudotime (**Fig. 6E, F**) revealed that the progenitor marker *mitfa*, melanophores markers, *dct* and *tryp1b*, and iridophore marker *alx4b* all processively turned off or switched on along pseudotime, as expected.

Differential geometry analyses provided complementary and in-depth mechanistic insights about pigment cell differentiation (**Fig. 6G**). Analyses of cell speed (the length of velocity vectors) and acceleration magnitude (the length of acceleration vectors) revealed that progenitors generally have low speed, reflecting a metastable cell state, whereas transitions of pigment progenitors and proliferating progenitors speed up when committing to a particular lineage, e.g., iridophore/ melanophore/Schwann lineage, etc. Cell divergence analysis indicated that those pigment progenitors and proliferating progenitors function as sources with high positive divergence, whereas differentiated cell types such as melanophores, iridophores, chromaffin, and Schwann cells function as sinks with strong negative divergence. Analysis of cell curvature magnitude (length of the curvature vectors), which measures the deviation of a single cell’s trajectory from being straight in the gene expression space at any point on this trajectory, revealed that large curvature emerges when a cell makes a fate decision (e.g., at the bifurcation point of the iridophore and melanophore lineages or of the neuron and satellite glia lineages).

Differential geometry analyses also lead to functional predictions. Genes with the highest acceleration in early progenitors tended to be highly expressed in terminal cell types (**Fig. 6G**). Interestingly, the top-ranked genes with the highest absolute acceleration from the previously unknown cell type were enriched in chondrocyte-related pathways, indicative of a potential chondrocytic origin (**Fig. 6H**). Furthermore, Jacobian analysis revealed potential regulation of the chondrocyte marker *slc36c2* by the pigment regulator *erbb3*, consistent with previous reports that *EGFR* (*erbb3*) signaling is critical for maintaining the chondrocyte lineage (Fisher et al. 2007). In addition, this analysis revealed a strong connection between chondrocyte-specific markers *col6a3 col6a, col6a2*, and *vwa1* (**Fig. 6I**). In iridophore cells, *pnp4a, fhl12a, crip2*, etc., are hub regulators (**Fig. 6J first row, left**), and *pnp4a* was potentially activated by *tfec* in the progenitors of iridophore lineage (**Fig. 6J first row, middle**) (Petratou et al. 2021), with possible repression occurring when *tfec* expression level was high in the mature iridophore cells (**Fig. 6J first row, right**).

Next, we extended our Jacobian analysis to other cell types or other published tsc- or cscRNA-seq datasets in order to identify key regulators and cell type (state)-specific regulatory networks (**Fig. 6J**, **Fig. SI6A–G**). In the cell-cycle-related vector field analysis of the sci-fate dataset, we found that the cell cycle-related gene *MT2A* (Lim et al. 2009) is a hub regulator whose expression is potentially repressed by *TOP2A* at the S stage when *TOP2A* expression is high (**Fig. 6J second row)**. Furthermore, based on datasets from either the sci-fate or the scEU-seq study, *E2F7* appears to repress *BRCA1* expression around the G1-S stage (**Fig. SI6D, F**) (Westendorp et al. 2012). For the GR response-related vector field analysis of the sci-fate dataset, we found that the GR response-related gene *SERPINE1* is a hub gene whose expression is potentially activated by *JUNB (Sundqvist et al. 2018)*, especially at the later time points of the GR response, although this activation saturates when *JUNB* expression is high (**Fig. 6J third row, right**). Strong putative repression from *GTF2IRD1* to *CEBPB* was also observed at a later stage of the GR response (**Fig. SI6E**). Our Jacobian analysis formally validated that there were no strong interactions between the cell-cycle progression and GR response in the sci-fate dataset, although there were strong interactions among cell cycle– related components (**Fig. SI6A-C**). For the intestinal organoid dataset, we found that *Lgals4*, *Dmbt1, Krt8*, etc., are hub genes of the Jacobian-based regulatory network (**Fig. 6J fourth row, right**). *Alphi* also appears to repress the intestinal stem cell marker *Lgr5* (Clevers 2017) when expressed at a low level, but starts to activate *Lgr5* expression at moderate expression, ultimately saturating when *Alphi* expression is high. From the mESC dataset, we found that the heat shock gene *Hsp90b1* is a hub gene of the mESC-specific regulatory network. *Hsp90b1* appears to weakly activate *Smc4* (**Fig. 6J fifth row**) (Bradley et al. 2012). Finally, analyses of a pancreatic endocrinogenesis dataset (Bastidas-Ponce et al. 2019) (**Fig. SI6G**) revealed differentiation dynamics and key gene interactions. The acceleration and divergence accurately highlight hotspots, including a saddle point in ductal cells (negative divergence), exit from this state to early endocrine progenitors (high positive acceleration), the bifurcation point for late progenitors to differentiate into stable cell types (high acceleration and positive divergence), and stable cell types (negative divergence). Jacobian analyses of several key genes agreed with previous experimental findings: 1) *Ngn3* activates *Pax4* in progenitors to initiate pancreatic endocrinogenesis (Arda, Benitez, and Kim 2013), 2) the toggle switch formed by the mutual inhibition between *Pax4* and *Arx* at the bifurcation point (Arda, Benitez, and Kim 2013), and 3) the activation of *Ins2* in beta cells by *Pdx1* (*Arda, Benitez, and Kim 2013*).

Our extensive analyses demonstrate that vector field reconstruction and differential geometric analysis enable quantitative discovery of regulatory mechanisms across various single-cell technologies, conditions, and biological systems.

### Vector Field Analyses Reveal Potential Host-virus Interaction and Resistance Mechanism of SARS-CoV-2 Infection

The ongoing COVID-19 (coronavirus disease 2019) pandemic, resulting from infection with SARS-CoV-2 (severe acute respiratory syndrome coronavirus 2), has prompted intense interest in the mechanisms of host-virus interactions and host resistance to viral infection. We wondered whether the vector field analysis we developed provides insights into the pathogenicity of SARS-CoV-2. To explore this possibility, we reanalyzed data from a study that investigated the single-cell response of SARS-CoV-1/2 infection in the permissive human epithelial cell line Calu-3 (Emanuel et al. 2020). Our analysis of the scRNA-seq data confirmed previous reports of transient early interferon signaling followed by a secondary nuclear factor-kappa B (*NF-kB*) response in cells infected by either virus, with SARS-CoV-2 triggering a stronger response (**Fig. SI7A-F**). Our vector field analysis also revealed host genes (*NFKB1A, HSP90AA1, PLAU*, etc.) and pathways (viral gene expression and interferon-gamma related pathways, etc.) related to virus infection in a time-dependent manner (**Fig. SI7G-J**).

The SARS-CoV-2 virus does not have introns. Hence, to detect putative interactions between host and virus genes, we designed a strategy for estimating viral RNA velocity by treating the *3’UTR* level as the “unspliced RNA” for all viral genes, and the amount of each viral gene as “spliced RNA” (See **MATERIAL AND METHODS**). Using this approach, we observed typical activation patterns as seen in regular RNA velocity phase plots for both structural (**Fig. 7A, B**) and nonstructural genes (**Fig. SI7K**). With the reconstructed composite host–virus vector field, we next investigated the top-ranked “regulators” and “effectors” from the host genome for each viral gene across time (**Table SI1**). We identified *LSM8, RAB21, RBPJ (TNFSF10, NFKBIA, PARP14*) as the top negative (positive) regulators (effectors) shared by viral genes in cells that are infected by SARS-CoV-2 for 12 hours (12-h cells) (**Fig. 7C, Fig. SI7L**). By compiling the top regulators and effectors of each viral gene/feature, we built networks of interactions between host and virus genes at each time point (**Fig. 7D, Fig. SI7M, Table SI1**). The majority of those genes were reported previously to be associated with SARS-CoV-2 infection (**Fig. 7E, Fig. SI7M, Table SI1**). For example, the *MX1* antiviral effector is activated after SARS-CoV-2 infection (Bizzotto et al. 2020), and the ADP-ribose binding sites *of14* of *PARP* share sequence homology with SARS-CoV-2 ADRP (Webb and Saad 2020). Enriched GO terms for those genes were associated with viral life cycle, regulation of tumor necrosis factor secretion, response to cytokines, etc. (**Fig. 7E, Fig. SI7N**). Those results collectively support the validity of our approach for identifying interactions between host and virus genes.

**Figure 7:**
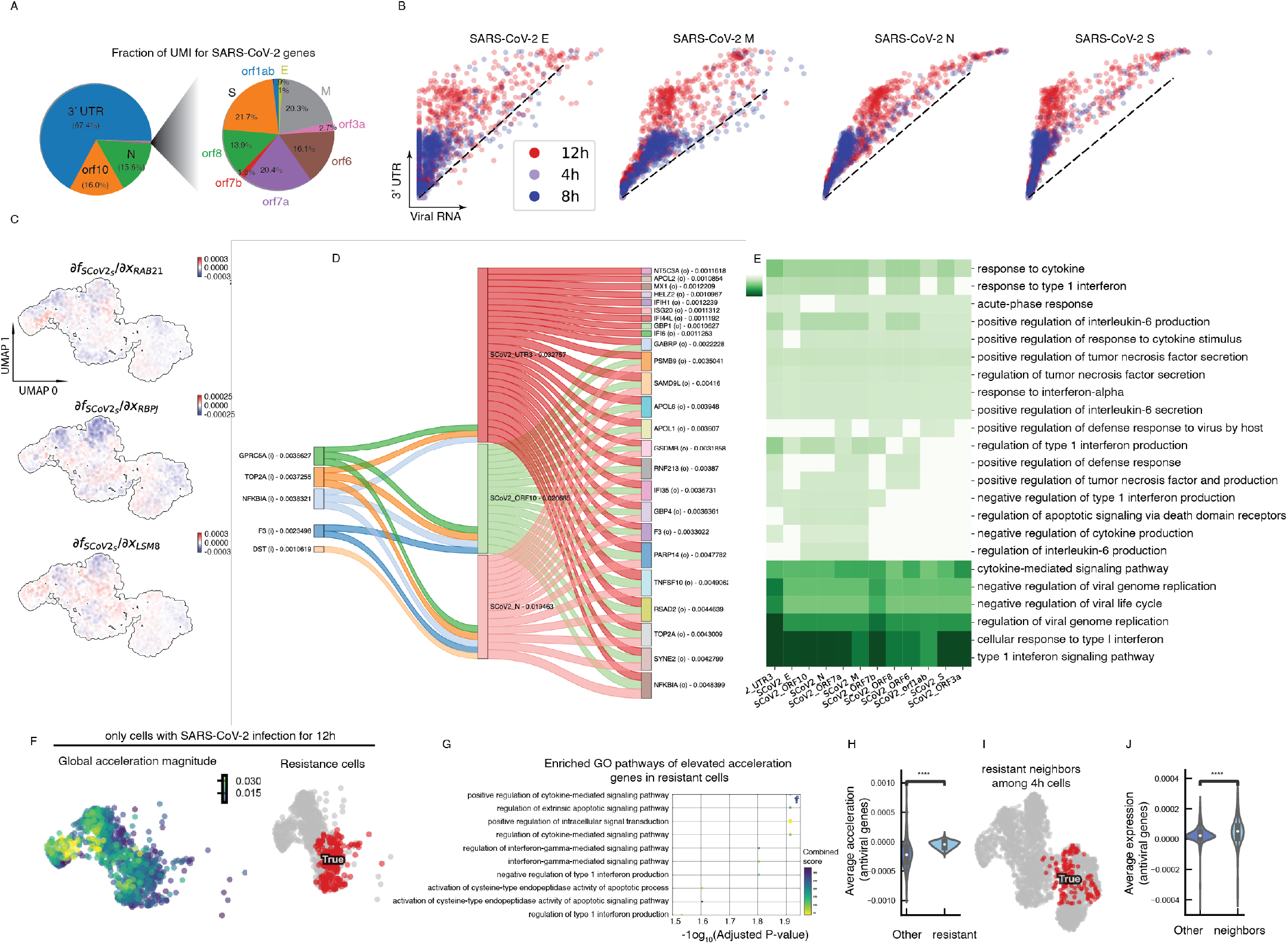
Vector Field and Differential Geometry Analyses of SARS-CoV-2 Infection Reveals Virus–host Interaction as well as Host Resistance Mechanisms. A. Pie chart of the UMI (unique molecular identifier) fraction of SARS-CoV-2 viral features/ genes among total viral UMI counts. *3’UTR, orf10*, and *N* of SARS-CoV-2 occupy the majority of viral UMI counts (**left**). The remaining genes are visible upon magnifying the corresponding small portion of the pie chart (**right**). When making the pie chart, UMIs counts for each feature/gene were aggregated across all cells. B. Treating the *3’UTR*, an indicator of viral load, as “unspliced RNA” and each structural gene [envelope (*E*), membrane (*M*), nucleocapsid (*N*), spike (*S*)] as “spliced RNA” reveals most cells are in the induction phase for all structural genes each with distinct kinetics (see phase plots for nonstructural genes in **Fig. SI7K**). C. Cell-wise Jacobian values of representative negative regulators (*LSM8, RAB21*, and *RBPJ*) of the SARS-CoV-2 *S* gene in the host cell, from 12-h cells, on the UMAP embedding (same as in **I**). D. Sankey diagram to visualize the putative host–virus gene interactions in 12-h cells. Interactions with average Jacobian values less than 1.05e-3 were filtered out in order to simplify the diagram. Numbers after gene names indicate computed average Jacobian values for each associated interaction or sum of all relevant interactions. “i”: input or regulator; “o”: output or effector based on the Jacobian analysis. E. Top effectors of each SARS-CoV-2 feature/gene from 12-h cells share commonly enriched GO pathways related to viral infections. For **C, D,** and **E,** see similar analysis for other time points and genes in **Fig. SI7L, M, N**. F. Scatterplots of global acceleration values of 12-h cells (**left**) and the location (**right**) of putative SARS-CoV-2–resistant cells on the UMAP embedding (see also **Fig. SI7O, P**). G. Differential acceleration analysis of putative resistant cells vs. background 12-h cells reveals that genes with elevated acceleration in resistant cells are enriched in antiviral pathways. Mann–Whitney–Wilcoxon test (two-sided) was used to calculate the p-value between two groups: other (non-resistant cells) and resistant (resistant cells). ****: *p* < = 1*e* – 04. H. Distribution of mean accelerations of genes related to antiviral pathways is up-shifted in putative resistant cells. I. Nearest neighbors of putative resistant cells among 4-h cells are enriched near the gap next to the regions dominated by 4-h cells on the UMAP embedding. J. Nearest neighbors of resistant cells among 4-h cells show elevated average expression of genes related to antiviral related pathways. The same test as in **G** was used, but for two groups, other (non-neighboring cells) and neighbors (neighbors of resistant cells among 4-h cells). ****: *p* <= 1*e* − 04.

SARS-CoV-2 infected cells exhibited considerable heterogeneity in their temporal dynamics, with some 12-h cells exhibiting delayed expression of viral genes (i.e., they were resistant to viral infection) (**Fig. SI7A middle**). We hypothesize that such resistant cells with moderate viral infection should exhibit an overall lower acceleration for transcriptomic RNA, but higher acceleration of antiviral genes, than non-resistant cells. Intuitively, one may make an analogy that the virus is the force that changes the global transcriptome of the host cell (i.e., transcriptome velocity), whereas the antiviral and immunological response of host cells provide friction against the virus infection by attenuating the change in the transcriptome velocity vectors. We first identified a group of 12-h cells with viral loading >30% of its maximum and with global acceleration magnitudes larger than those of most other 12-h cells (**Fig. 7F**). On the UMAP embedding, those cells are far away from most other 12-h cells and interestingly locate to a gap in UMAP space near 4h cells (**Fig. 7F, Fig. SI7O, P**). Genes in these cells with higher acceleration values than those in other 12-h cells were enriched in cytokine-mediated or interferon-related pathways (**Fig. 7G, H**). We treated the cells collected at 4 h that were nearest neighbors to these putative resistant cells as the possible initial cell states of the latter. These 4h cells also exhibited considerably higher expression (**Fig. 7I, J**) and acceleration (**Fig. SI7Q**) of genes related to antiviral pathways than other 4-h cells. **Fig. SI7R** provides a plausible mechanistic explanation. In infected cells, antiviral responses race against virus proliferation, and the responses are likely sigmoidal, a pattern that appears ubiquitously in signal transduction and gene regulation (Alon 2019); this is supported by the existence of self-activation among antiviral genes (**Fig. SI7S**). A characteristic of these sigmoidal dynamics is that response time is sensitive to the initial conditions, i.e., the basal levels of the antiviral response genes, as much of the response time is spent at the initial accumulation stage (**Fig. SI7R**) (Zhang NPJ Syss Biol). Resistant cells with initially high antiviral-related gene expression can quickly turn on (with positive acceleration) and reach a plateau (reflected by close to zero or negative acceleration) to block or delay the effects of viral infection and virus accumulation, consistent with the gap between the 4-h cells with higher antiviral gene expression/acceleration and other cells, as well as the delayed progression of the putative resistant 12-h cells based on the streamline plot (**Fig. SI7A**).

In summary, our vector field approach allowed us to identify putative host–virus interactions and resistance to SARS-CoV-2 infection. Thus, this method has the potential to shed light on the mechanisms of many other diseases, including viral infections and cancers.

## DISCUSSION

In *The Strategy of the Genes*, Waddington explained the formation of the epigenetic landscape using the “guy-rope” model, in which genes and the interactions between them determine the topography of the landscape (Waddington 1957). Our vector field framework provides an important mathematical realization of this model, not only revealing the transitions between cell states but also providing testable hypotheses regarding the governing mechanism of biological processes.

Our analytical framework consists of three integral stages. First, we estimate genome-wide kinetic rate constants and RNA velocity vectors from single-cell data. Next, we use RNA abundance as well as velocity vectors to reconstruct the single-cell vector field functions. Finally, we apply single-cell differential geometry analyses made possible by the analytical vector field function, thereby obtaining biological insights. In the first stage, we explicitly model metabolic labeling while integrating it with splicing dynamics in metabolic labeling–based scRNA-seq or time-resolved scRNA-seq (tscRNA-seq) data to accurately infer the kinetic parameters governing each step in the RNA’s life cycle. Our estimated kinetic rate constants, which are more accurate than those obtained using the original RNA velocity method, provided multiple functional insights, including the high synthesis rates of mitochondrial genes and the global and proportional increase in RNA stability of human genes relative to their mouse orthologs. The latter observation expands on recent findings that biochemical reactions, especially protein degradation, are consistently slower in human cells than in their mouse counterparts during both embryonic segmentation (Matsuda et al. 2020) and motor neuron differentiation (Rayon et al. 2020). By jointly analyzing intron/exon and labeling information from tscRNA-seq experiments, we reconcile the kinetics obtained by metabolic labeling with those obtained with RNA splicing, unexpectedly revealing a subset of RNAs with slower splicing than degradation. Because our estimation approach implements a universal modeling system, the approach is compatible with all existing single-cell RNA metabolic labeling strategies, as well as new labeling protocols that may be developed, such as dual labeling with 4sU and 6-thioguanine (6-TG) to directly measure RNA acceleration (Kiefer, Schofield, and Simon 2018). This comprehensive approach to estimate absolute kinetic parameters will therefore have broad application in RNA biology. Finally, the total RNA velocity estimated from tscRNA-seq data using our framework significantly improves the consistency of RNA velocity flow between a cell and its neighbors while simultaneously providing absolute velocity vector quantification, thereby alleviating the limitations of conventional RNA velocity analyses.

In the second stage, we take the discrete, sparse, and noisy single-cell velocity vectors as input to robustly learn a continuous vector field function. Early efforts in pseudotime ordering, which recovers central trajectories of cell populations (Trapnell et al. 2014), RNA velocity, which estimates local velocity direction of sampled cells (La Manno et al. 2018), and sci-fate, which builds linkages that connect the most likely historical and current states sequentially over time (Cao, Zhou, et al. 2020), constitute key developments in dynamics inference. However, it has taken until now to reconstruct analytical and continuous vector field functions in transcriptomic space, as we have described in this work. Our vector field reconstruction takes advantage of the power of advanced machine learning (ML) to scalably and accurately learn the vector field functions even for datasets with hundreds of thousands of cells (e.g., murine hematopoiesis). With the reconstructed continuous vector field function, we can predict the cell states over an extended time period in the past or future, as evidenced by our analysis of sequential transcriptomic profiling and clone fate tracing for neutrophil differentiation or murine hematopoiesis. Our method is also capable of *in silico* tracing the transcriptomic dynamics of cell ensembles over time, which may provide an important complement to live-cell imaging (Baker 2010) or lineage tracing (McKenna et al. 2016; Frieda et al. 2017; Chan et al. 2019).

In the third stage, we enable the interpretation of our vector field function by applying predictive dynamical systems methods and differential geometry analyses that extract regulatory information from the vector field function. The differential geometry analyses reveal features, including fixed points, Jacobians, acceleration, curvature, and divergence, that are only made possible by the reconstructed differentiable vector field functions. These features, in turn, provide insights into regulatory mechanisms in various biological processes that are not otherwise transparent. For example, in the analysis of the zebrafish dataset, the stable and unstable fixed points corresponded to progenitors and terminal cells, respectively; the topranked acceleration genes pointed to a chondrocyte lineage origin that was not apparent from total RNA analysis; and the Jacobian analyses revealed a plausible interaction between the the pigment cell regulator *erbb3* and the chondrocyte marker gene *slc35c2*. Interestingly, our vector field analyses revealed that high initial expression levels of antiviral genes may lead to resistance to SARS-CoV-2 virus infection, reflecting the nonlinear dynamics nature of the innate immune responses. These findings in disparate systems point to our framework’s general ability to reveal regulatory mechanisms across various types of single-cell technologies, conditions, and biological processes.

Although this work establishes a general and powerful framework, important directions remain for further development. First, our vector field learning approach focuses on the deterministic part of the dynamics, i.e., the convection part of a convection–diffusion process (Cho and Rockne 2019) in transcriptome space. Biological systems are, however, intrinsically stochastic; accordingly, future work should seek to reconstruct the (stochastic) diffusion part of the model as well. Second, one limitation of single-cell data is the low RNA capture rate. As RNA capture sensitivity, sequencing depth, and sample size increase, our framework will further facilitate the discovery of regulatory mechanisms by decreasing the uncertainty from the measurements and estimation (Chapman et al. 2020). Third, the reconstructed vector field functions can be confounded by unobserved hidden variables, e.g., environmental cues from surrounding cells, their epigenetic states, or protein abundance. Incorporating datasets from the recent developments of single cell multi-omics (Cao 2020; S. Ma et al. 2020), spatial transcriptomics (Moffitt et al. 2018; Rodriques et al. 2019), or both (Liu et al. 2020) into our framework will provide the opportunity to address the hidden variable problem in our vector field analysis. Finally, in the current form of this approach, we use the Gaussian kernel in vector field learning because of its simplicity and effectiveness. An interesting future direction would be to express the vector field in a form with a direct biological interpretation, e.g., the sigmoid functions widely used in mathematical modeling of biological networks, which could further facilitate interpretation of the reconstructed vector field.

In summary, we have built a general framework for the interpretation of expression kinetics that can be applied to numerous systems compatible with single-cell genomic profiling. For example, vector field reconstruction and differential geometry analysis may help fully resolve the regulatory cascade associated with infections by viruses (Hein and Weissman 2021; Emanuel et al. 2020) including CMV, SARS-CoV-2, and others; dissect the complicated differentiation hierarchy of hematopoiesis; and pinpoint the precise regulatory mechanisms underlying normal or rare hematopoietic lineage bifurcation or switches, etc. With the rapid expansion of RNA metabolic labeling methods, especially those that can explore large numbers of cells (Q. Qiu et al. 2020), ***dynamo*** will be an increasingly important tool for using those technologies to study RNA kinetics specific to developmental stages or species, or relevant to particular diseases, as well as their coordination and regulatory logic, across a variety of labeling strategies and biological systems. More broadly, the general and scalable framework of kinetic parameter estimation and vector field function analysis developed in this study, especially when coupled with RNA metabolic labeling, lineage tracing (McKenna et al. 2016; Frieda et al. 2017; Alemany et al. 2018; Chan et al. 2019), RNA age (Rodriques et al. 2020), signal pathway recording (Sheth and Wang 2018), as well as genetic perturbations (Adamson et al. 2016; Dixit et al. 2016), will enable us to move towards holistic kinetic models and theories of the entire organism for cell atlas projects (Cao, O’Day, et al. 2020) comprising many cell types and encompassing many conditions or species.

## Supporting information

Supplementary table for SARS-CoV-2 data analysis

Supplementary animation

Supplementary animation

Supplementary animation

Supplementary animation

Supplementary animation

Supplementary animation

## ACKNOWLEDGEMENT

We thank Jiayi Ma and Tatsu Hashimoto for technical discussions on recovering vector fields and potential functions; Alex Ge and Luke Gilbert for help with culture of HL60 cells and implementation of scSLAM-seq; Jay Shendure, Cole Trapnell, and June Cao for sharing sci-fate datasets; Macro Hein, Sara Sunshine, and Joseph Min for estimating velocities for the SARS-CoV-2 viral genes/features; Vedran Franke and Emanuel Wyler for defining cells resistant to COVID-19; Laren Sanders for zebrafish dataset analyses; and Hao Wu, Marco Jost, Tomas Norman, Kara Mckinley, Richard She, Yuancheng (Ryan) Lu, Qi Qiu, and other members of the Weissman lab for discussion and comments on this work.

## AUTHOR CONTRIBUTIONS

X. Q., J. X., and J. S. W. conceived the project. X. Q., Y. Z., and J. X. developed the overall model and theory. Y. Z. carried out most of the derivations with suggestions from J.X. and X. Q. X. Q., S. H., D. Y., and J. S. W. designed the experiments. S. H., X. Q., D. Y., M. S., A. P., and J. M.R. performed the experiments. X. Q. and Y. Z. performed the analyses. All authors contributed to the writing of the manuscript.

---

**Supplementary Figure 1:**
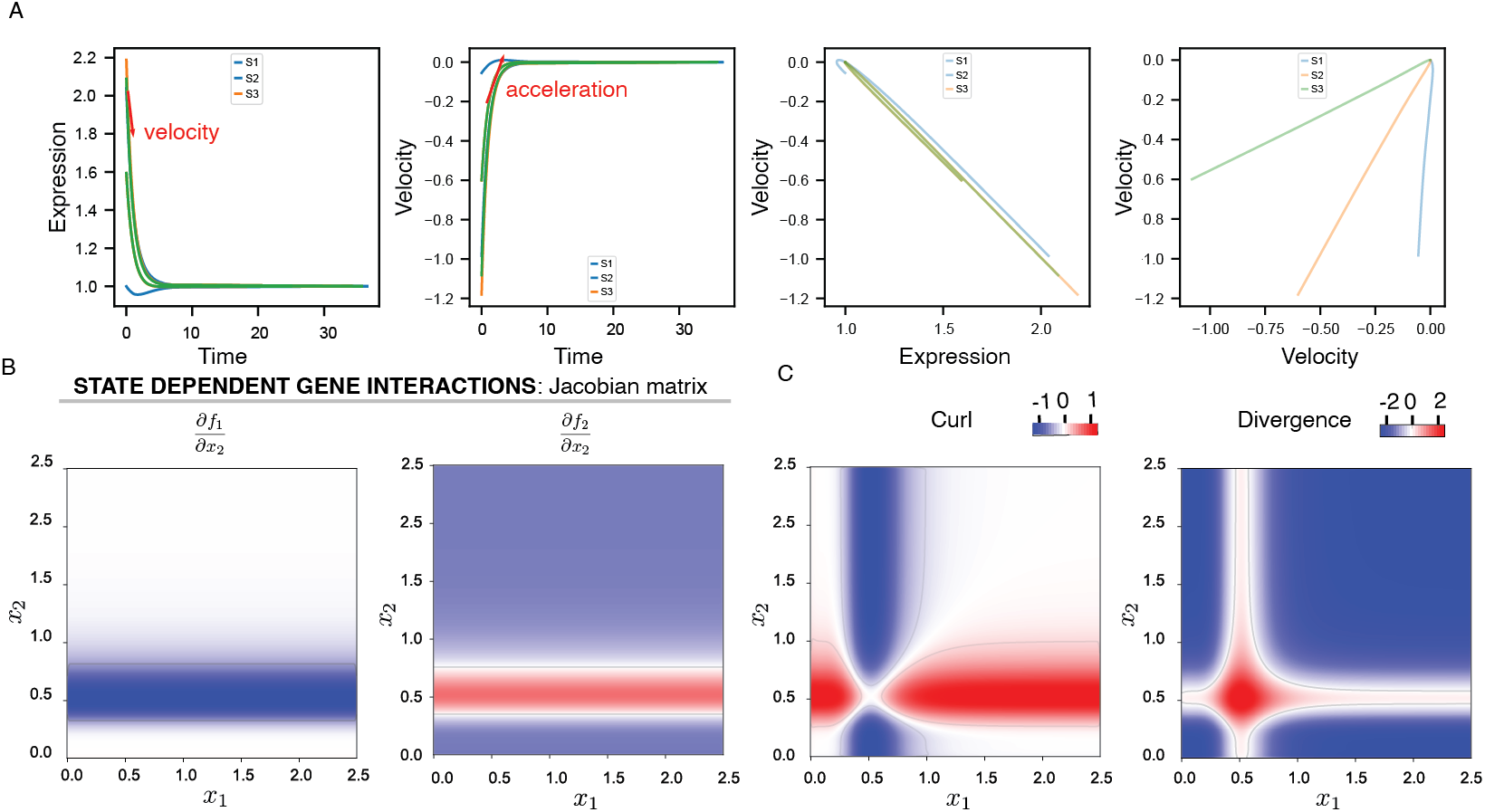
Various Ways to Quantify Expression Dynamics and additional Differential Geometry Analyses of Vector Fields. A. Gene expression dynamics along the indicated trajectories in **Fig. 1A (4)**. **1**) Gene expression quickly decreases, **2**) while velocity rapidly approaches 0 over time. Taking the derivative of the expression or velocity with respect to time along the indicated trajectory gives velocity **1**) or acceleration **2**), respectively, represented by red arrows. **3**) Increasing the expression of one gene linearly decreases the velocity of the other genes. **4**) The velocity of gene *x*_1_ positively correlates with that of *x*_2_, but with different strengths across the three trajectories. B. The Jacobian of *∂**f***_1_/*∂x*_2_ (**left**), *∂**f***_2_/*∂x*_2_ (**right**) **a**long the horizontal dashed line indicated in **Fig. 1 A4.** Two other symmetric Jacobian elements, *∂**f***_2_/*∂x*_1_ (left), *∂**f***_1_/*∂x*_1_, are shown in **Fig. 1C**. C. Map of the curl (defined only in two or three dimensions ∇ × ***f*** = *∂**f***_2_/*∂x*_1_ – *∂**f***_1_/*∂x*_2_) and divergence 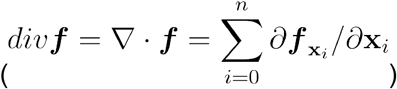 landscapes in the phase space of the two-gene system.

**Supplementary Figure 2:**
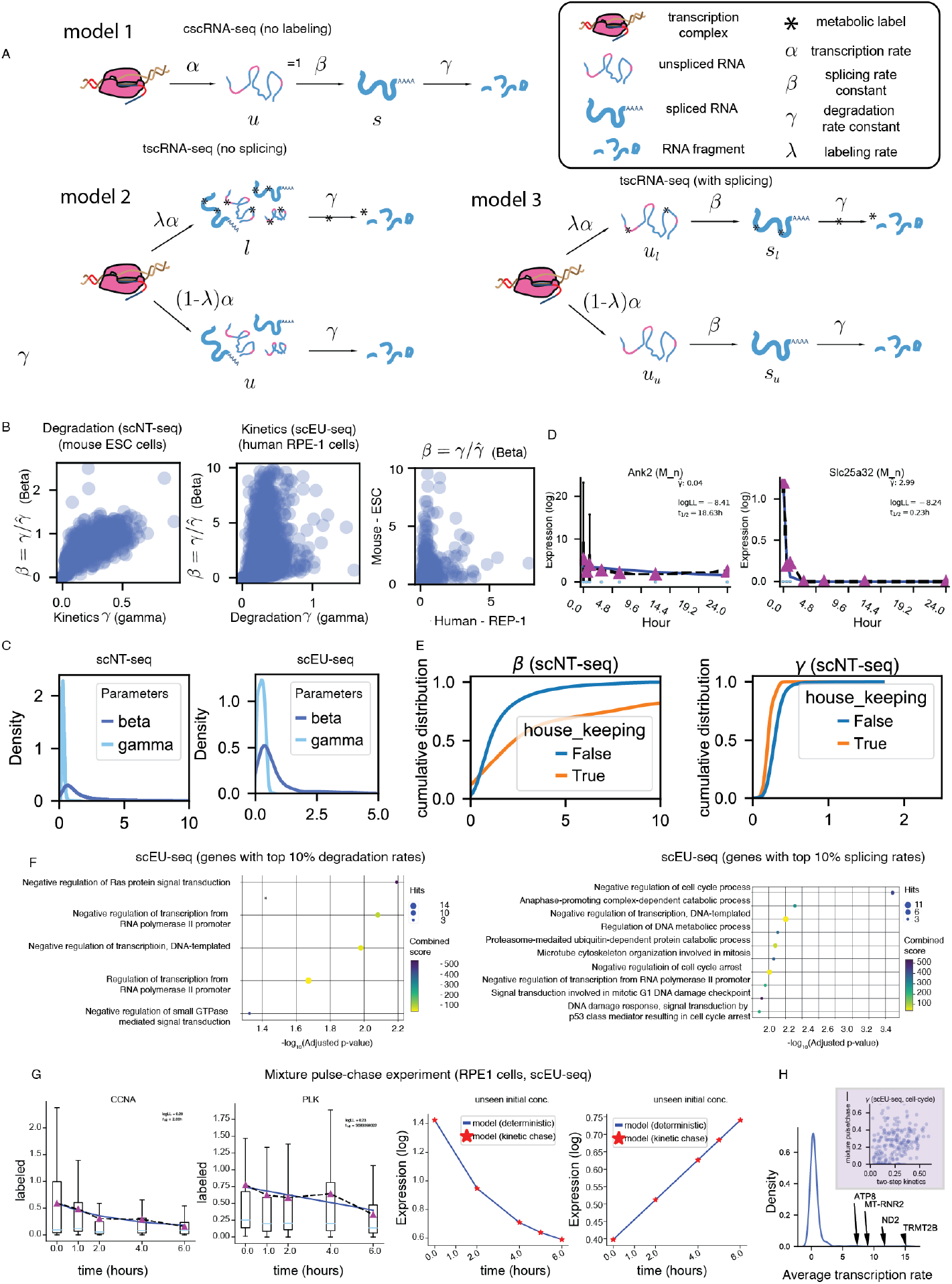
Comprehensive Expression Kinetics Estimation Framework in ***Dynamo*** and Global Analyses of Transcription, Splicing, and Degradation Rate Constants. A. Three main models of cscRNA-seq data (**Model 1**) and tscRNA-seq data that do not incorporate splicing (**Model 2**) or do (**Model 3**). B. Estimating RNA degradation and splicing rates with data from degradation or kinetics labeling tscRNA-seq experiments. Scatterplot of 1) degradation rates *γ* estimated from labeling data, and slopes of the unspliced–spliced plane 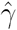 estimated from splicing data from the scNT-seq study with mouse ESC cells on the **left,** and 2) degradation rates *γ* and the splicing rate 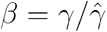 from the human RPE-1 cells from the scEU-seq study in the **middle**. The murine splicing rate constant (*β*) calculated based on scNT-seq data is generally higher than that for humans calculated based on scEU-seq data (**right**). C. Deterministic first-order decay model fitting of *Ank2* (slow degradation) and *Slc25a32* (fast degradation) chase data, using the ESC experiment data from the scNT-seq study (Q. Qiu et al. 2020). D. Splicing rate constants (*β*) are in general much larger than the degradation rate constants (*γ*) in both the scNT-seq (**left**) and scEU-seq (**right**) dataset analysis based on the density plot. E. Housekeeping genes tend to have faster splicing (**left**) but slower degradation (**right**) than other genes based on the cumulative distribution plot. F. The top 10% genes from the scEU-seq dataset with highest splicing (**left**) or degradation (**right**) are enriched in transcription and cell cycle–related pathways. G. Demonstration of estimating kinetic parameters from a mixture pulse-chase experiment from the scEU-seq study (Battich et al. 2020), using also its non-steady state model. H. Genes with highest transcription rates are all mitochondrially encoded. I. Degradation rates estimated from the non–steady-state model of the mixture pulse-chase experiment are consistent with those estimated from the degradation experiment.

**Supplementary Figure 3:**
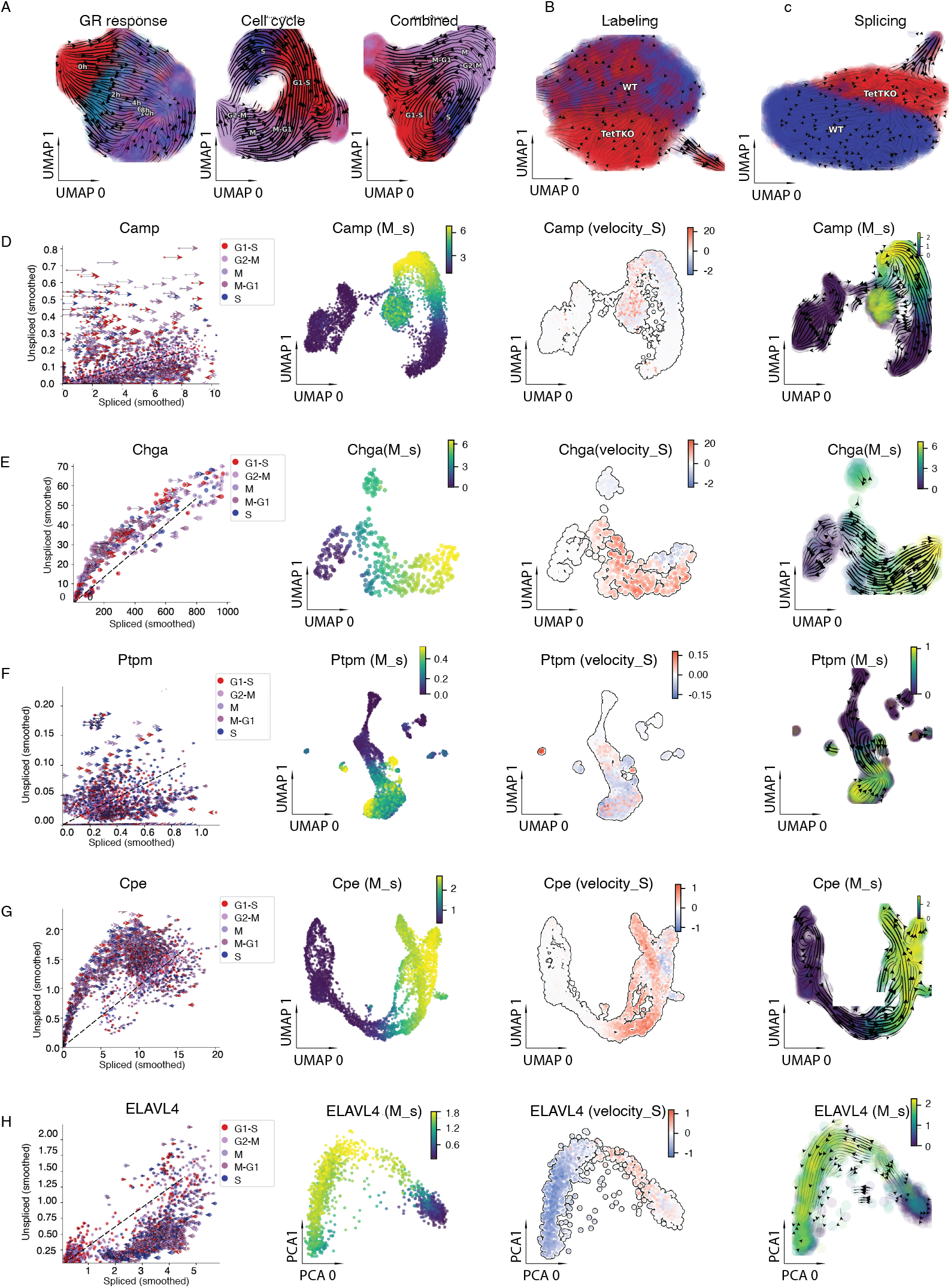
***Dynamo*** Estimates RNA Velocity Robustly and Accurately across a Variety of tscRNA-Seq or cscRNA-Seq Experiments. A. Streamline plots with only GR-related genes (**left**), cycle–related genes (**middle**), and a combination of GR and cell cycle–related genes (**right**) for the sci-fate dataset on UMAP embedding. The right panel is the same as the right panel in **Fig. 2C** but is annotated with inferred cell-cycle stages. B. Cells with triple KO of *Tet 1/2/3* (TetTKO) are biased to differentiate into 2C-like cells, based on the splicing RNA velocity streamline plot produced with ***dynamo***. C. Cells with TetTKO are biased to differentiate into 2C-like cells, based on the labeling RNA velocity streamline plot produced with ***dynamo***. Both **B** and **C** are the same as the middle and bottom panels from **Fig. 2E** but are annotated with cell genotype information. D. RNA velocity analyses of the BM dataset (Petukhov et al. 2018) with ***dynamo*.** First column: phase plot of the example gene *Camp;* second column: single-cell gene expression of *Camp* (normalized and locally smoothed) on UMAP embedding; third column: single-cell RNA velocity of *Camp* on the UMAP embedding; fourth column: streamline plot of the project RNA velocity on UMAP space. E. Same as above but for *Chga* in the chromaffin dataset (Furlan et al. 2017). F Same as above but for *Ptpm* in the dentate gyrus dataset (Hochgerner et al. 2018). G. Same as above but for *Cpe* in the pancreas dataset (Bastidas-Ponce et al. 2019). H. Same as above but for *ELAVL4* in the HG dataset (La Manno et al. 2018).

**Supplementary Figure 4:**
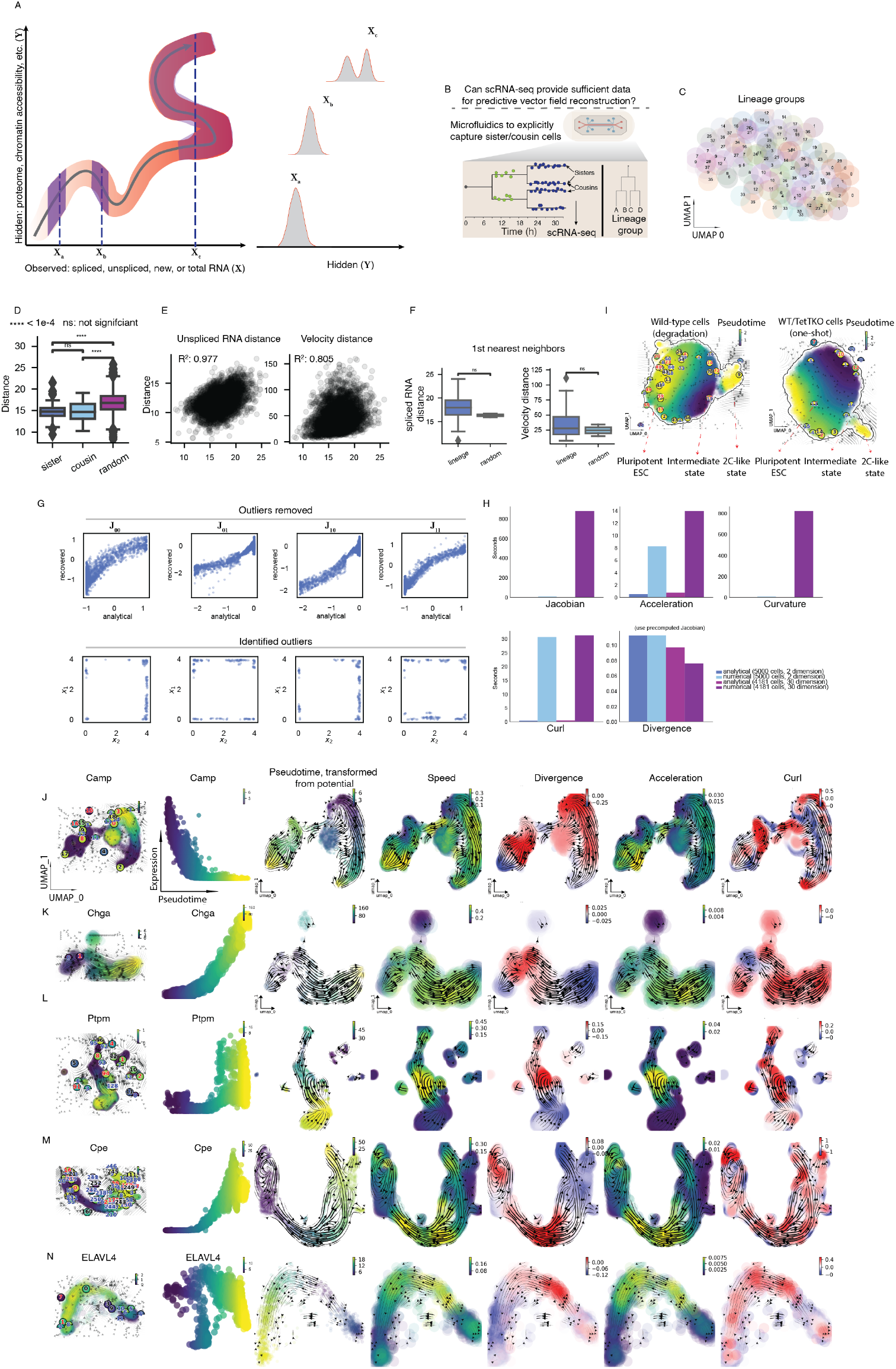
***Dynamo*** Enables scalable and accurate Reconstruction of Vector Field Functions and Characterization of Vector Field Topologies. A. Existence of hidden variables may confound vector field reconstruction. The averaging intervals (shaded box, **left**) at points **X**_*a*_ and **X**_*b*_ have single-peaked distributions along **X** as well as the unmeasured **Y** (shaded density plot, **right**), whereas the interval at point **X**_*c*_ has a two-peaked distribution for unmeasured **Y** (shaded density plot, **right**). Vector field reconstruction is expected to perform well when hidden variables from the system correlate with the observed variables (e.g. at **X**_*a*_, **X**_*b*_), but less well when they are loosely coupled (e.g. **X**_*c*_). B. Microfluidic platform design and experimental scheme to capture sisters/cousins from primary activated murine *CD8+* T-cell. Adapted from **Fig. 1** of (Kimmerling et al. 2016). C. RNA velocity streamline plot of cells on UMAP embedding. Cells are colored by lineage groups (i.e., sister or cousin cells) of single cells. D. Boxplot of the expression distance distribution (in the PCA space) of sister and cousin cell pairs, as well as that of random cell pairs. Mann–Whitney–Wilcoxon test (two-sided) was used to calculate the p-value between groups. ****: *p* <= 1*e* − 04. E. Scatterplots of the distance of spliced RNA expression states of single cells vs. the distance of unspliced RNA expression states, and vs. that of RNA velocity vectors of single cells show strong correlations. Distances were calculated in PCA space. R-squared value (*R*^2^) is shown for each panel. F. Distances between first-nearest neighbor cells show no difference among cells from the same or different linkages. Mann–Whitney–Wilcoxon test (two-sided) was used to calculate the p-value between groups. **: *p* <= 1*e* − 02. G. Pairwise scatterplots of estimated and analytical Jacobian elements (indicated by the equations for each column) from corresponding cells (**top**) and the identified outlier cells on the gene expression space of *x*_1_, *x*_2_ (**bottom**). Accuracy of estimated Jacobian deteriorates in boundary regions of sampled cells due to insufficient and biased sampling. H. Analytical differential geometric analyses enable nearly 1000x faster computation than state-of-the-art numeric algorithms ([numdifftools](https://pypi.org/project/numdifftools/)). I. Fixed points and vector field–based pseudotime for wild-type or the Tet1/2/3-KO cells on UMAP space. J. Reconstruction of vector field and characterization of the topology of the BM dataset (Petukhov et al. 2018) with ***dynamo***. First column: Topography plot of the system in UMAP space with cell colored by expression of *Camp* (normalized and locally smoothed); second column: gene expression of *Camp* vs. vector-field based pseudotime; third through seventh columns: single-cell vector-field based pseudotime, speed, divergence, acceleration, and curl on UMAP spaces. K. Same as above but for *Chga* in the chromaffin dataset (Furlan et al. 2017). L. Same as above but for *Ptpm* in the dentate gyrus dataset (Hochgerner et al. 2018). M. Same as above but for *Cpe* in the pancreas dataset (Bastidas-Ponce et al. 2019). N. Same as above but for *ELAVL4* in the HG dataset (La Manno et al. 2018).

**Supplementary Figure 5:**
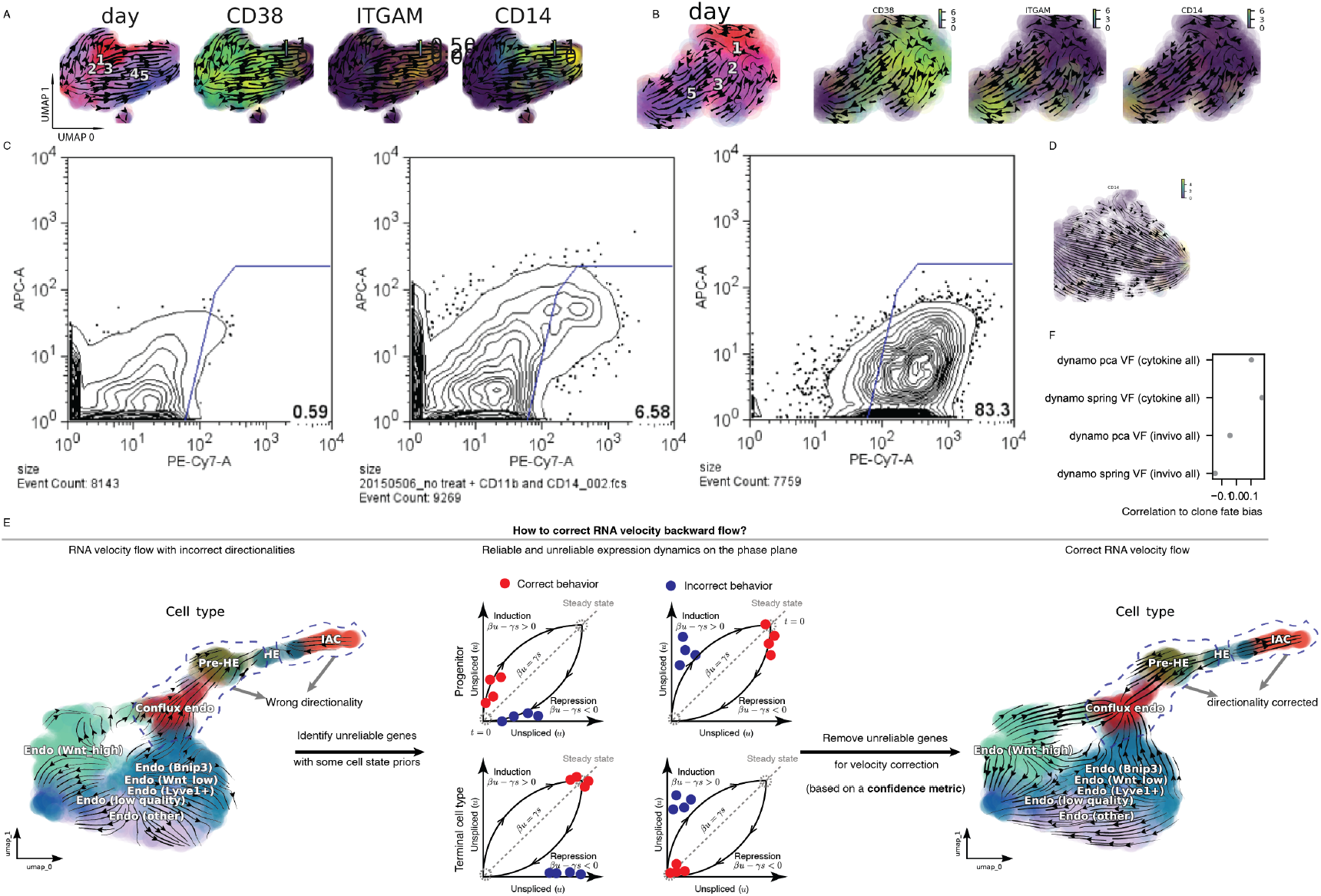
Vector Field Predicts long-range Cell Fate Commitments during Hematopoiesis. A. Expression of *CD38, ITGM*, and *CD14* over days 1, 2, 3, and 5 during HL60 neutrophil lineage commitment, and streamline plot in the UMAP space (same for **B, D**) from the 10x dataset. B. Same as above but with the splicing data from the clonal tracing scSLAM-seq experiment. C. Contour plot of APC vs. PE-Cy7 on days 1, 3, and 5. APC: *CD14* (monocyte marker), PE-Cy7: *CD11b* (neutrophil/monocyte marker). Gate was used to select CD11b+ positive cells to indicate maturation of the neutrophil-like lineage. D. Same as in **C** but for gene *CD14* and with the corresponding labeling data. E. Correcting RNA velocity flow with broad lineage hierarchy information in prehematopoietic stem cell formation (Zhu et al. 2020). See more discussion in **SUPPLEMENTARY METHODS**. F. Correlation of clone fate bias based on single cell fate predictions of the reconstructed vector field for the cytokine perturbation and in vivo datasets.

**Supplementary Figure 6:**
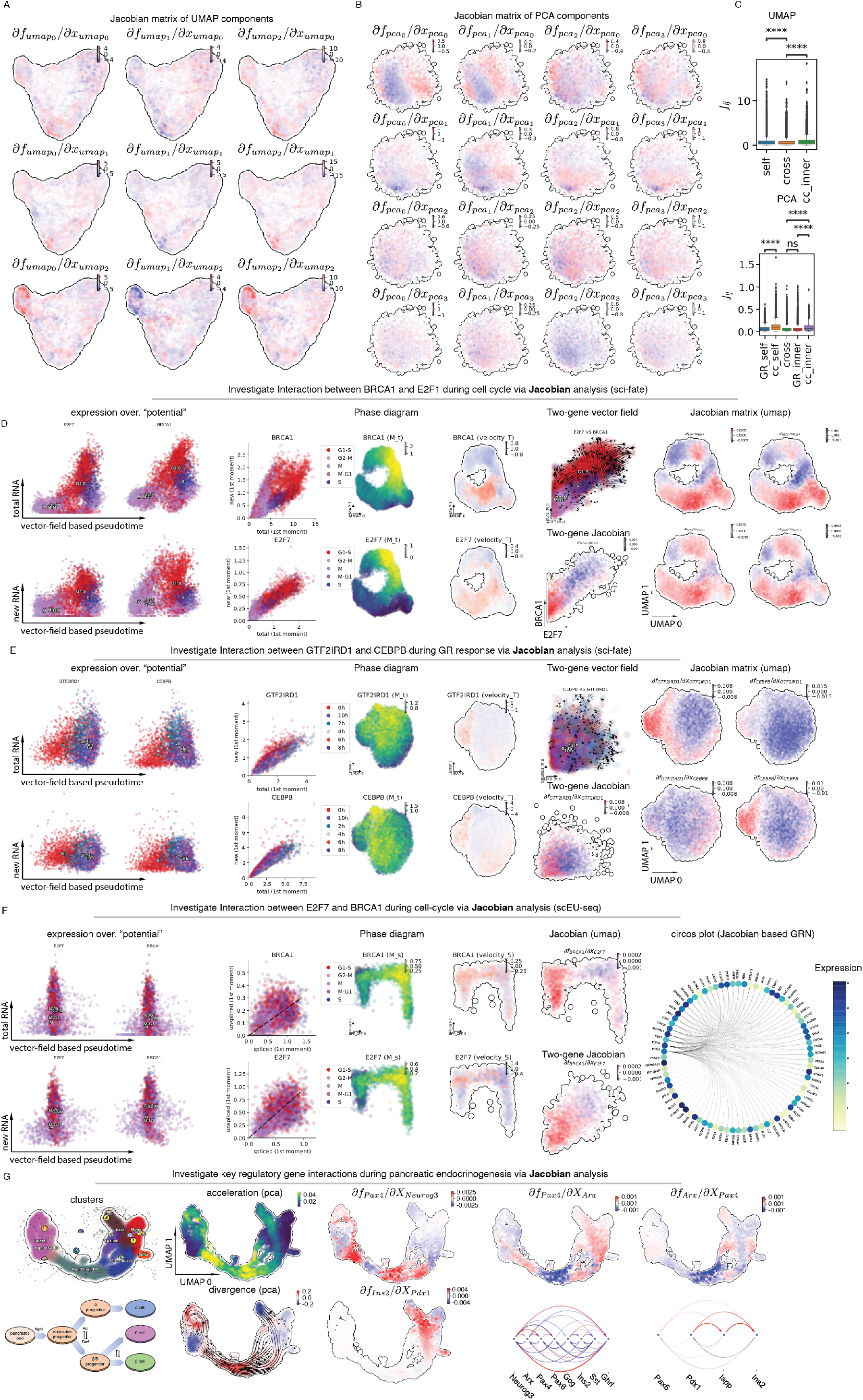
Differential Geometry Analyses of the Reconstructed Vector Field Function with ***Dynamo*** Predict potential Gene Regulation Mechanism across a Variety of tscRNA-Seq or cscRNA-Seq Experiments. A. Jacobian matrix analysis of UMAP components across cells based on the vector field reconstructed in a 3-dimensional UMAP space (data from sci-fate study, same as **B–C**). The first UMAP component corresponds to the GR response, whereas the second and third components correspond to the cell cycle. B. Jacobian matrix analysis of PCA components across cells based on the vector field reconstructed in 4-dimensional PCA space. The first two PCA components correspond to the GR response, whereas the second two components correspond to the cell cycle. C. Boxplots of Jacobian between cell cycle–related or GR response–related components and that between the cell cycle and GR response components, confirming that the cell cycle and GR response are independent of each other. Mann–Whitney–Wilcoxon test (two-sided) was used to calculate the p-value between groups. ****: *p* <= 1*e* − 04. D. Interaction between *BRCA1* and *E2F1* during the cell cycle (sci-fate dataset). **First column**: *BRCA1* and *E2F1* expression over vector field–based pseudotime. **Second column**: Phase-plot, single-cell gene expression in UMAP embedding, and single-cell velocity in UMAP embedding. **Third column**: top, vector field of *BRCA1, E2F1* gene state space; bottom, single-cell Jacobian of *E2F1* to *BRCA1* in *BRCA1, E2F1* gene expression space. **Fourth column**: Jacobian matrix across single cells visualized in UMAP embedding. E. Same as above but for *GTF2IRD1* and *CEBPB* and the GR response (sci-fate dataset). F. Same as above but for *GTF2IRD1* and *CEBPB* and the GR response (scEU-seq dataset). The Jacobian matrix plots in the rightmost column were replaced with the circosPlot of putative genes associated with the cell cycle. G. Dynamics and key gene interactions during pancreatic endocrinogenesis, revealed by differential geometry analyses. **First column**: 2D UMAP vector field topology shows stable fixed points (attractors) in alpha, beta, and ductal cells, and a saddle point at the branching point between ductal cells and early progenitors. The lower diagram illustrates the differentiation process of pancreatic endocrine cells and key regulatory genes/motifs. **Second column**: High acceleration is observed at the interface between early and late progenitors (magnitude change in velocity), and the bifurcation point where progenitors differentiate into stable cell types (direction change in velocity). Negative divergence is observed at the saddle point and attractors, and positive divergence at the bifurcation point and cell cycle of pancreatic buds. Third column: Jacobian analyses suggest that 1) *Ngn3* activates *Pax4* in progenitors, initiating pancreatic endocrinogenesis; and 2) *Pdx1* activates *Ins2* in beta cells. **Last four figures**: Jacobian analyses reveal mutual inhibition of *Pax4* and *Arx* at the bifurcation point in progenitors. Arcplot shows gene regulatory networks in progenitors and beta cells.

**Supplementary Figure 7:**
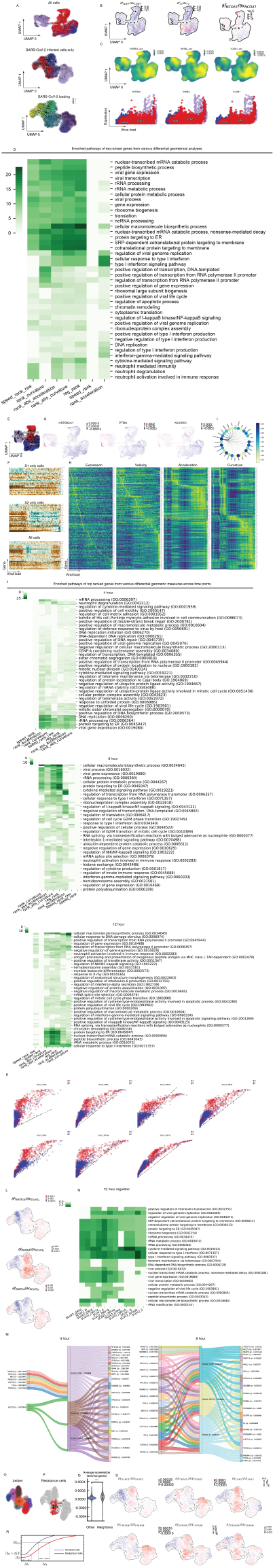
Vector Field Analyses Reveal Virus–Host Interaction and Resistance to SARS-CoV-2 Infection. A. Streamline plots of all collected cells (**top**), colored by viral infection conditions (s1: SARS-CoV-1 infection; s2: SARS-CoV-2 infection; mock: no viral infection), and SARS-CoV-2– infected cells, colored according to viral infection time (**middle**) or SARS-CoV-2 virus loading (**bottom**). The embedding from (Emanuel et al. 2020) was used for all cells (**top**), whereas the UMAP embedding produced from ***dynamo*** was used for the other two plots. Group names are placed at the median coordinates of cells belonging to a particular group, and those cells are also depicted in the same color, as below. B. Cell-wise Jacobian values of representative top-ranked self-interaction genes *CEBPE, F3*, and *TOP2A* in the “basal” cell cluster (cell groups enriched for mock cells and cells located at the infection stage, see **Fig. SIA**) on the UMAP embedding. C. Cell-wise acceleration values of representative top-ranked positive acceleration genes *NFKB1A, INHBA* and *ICAM1*, in the “basal” cell cluster on the UMAP embedding (**top**). **Bottom**: Expression (y-axis) dynamics are plotted as a function of viral loading (x-axis). D. Various differential geometrical analyses reveal commonly enriched GO pathways related to viral infections. E. Basal cluster, SARS-CoV-1 or SARS-CoV-2 branch, on UMAP embedding. F. Expression dynamics of antiviral response genes as a function of viral loading in cells infected with either SARS-CoV-1 or SARS-CoV-2, or in all cells. G. Cell-wise acceleration values of representative top-ranked negative acceleration genes *HSP90AA1, PTMA*, and *NUCKS1* in the “basal” cell cluster on the UMAP embedding. H. Kinetic heatmap of all host genes corresponding to proteins previously reported to bind to SARS-CoV-2 virus proteins (Gordon et al. 2020) as a function of viral loading. Genes are row-scaled and ordered based on the position of their maximal value on the x-axis (viral loading) across all cells from left to right, and independently for each quantity. I. Arcplot of the network between the top regulators for all host genes corresponding to proteins previously reported to bind to the representative SARS-CoV-2 virus protein *Orf9c*. J. Enriched pathways of top-ranked genes from various differential geometric measures across each time point (4, 8, and 12 h). K. The “spliced-unspliced” RNA phase plot of virus nonstructural genes after treating 3’ UTR, an indicator of viral load, as the “unspliced RNA”, and each nonstructural gene (*orf1ab, orf3a, orf6, orf7a, orf7b, orf8, orf10*) as “spliced RNA” for each structural gene. See phase plots for structural genes in **Fig. 7E**. L. Cell-wise Jacobian values of representative top positive host effectors *TNFSF10, NFKBIA*, and *PARP14*, from 12-h cells for SARS-CoV-2 gene *S* on the UMAP embedding. M. Sankey diagram to visualize the putative host–virus gene interactions in 4- and 8-h cells. Interactions with average Jacobian values less than 1.05e-3 were filtered out to simplify the diagram. Numbers after gene names indicate the computed average Jacobian values for each associated interaction or sum of all relevant interactions. “i”: input or regulator; “o”: output or effector based on the Jacobian analysis. N. Top regulators of each SARS-CoV-2 feature/gene from 12-h cells share commonly enriched GO pathways related to viral infections. O. Leiden clustering of SARS-CoV-2–infected cells. 300 cells with lowest global acceleration magnitude that overlap with clusters 0, 2, 4, 5, 6 were used to define putative resistant cells (see also **Fig. 7F**). P. Distribution of putative resistant cells (colored with red) on UMAP embedding of all cells. Q. Distribution of mean acceleration values of genes related to antiviral related pathways is up-shifted in 4-h neighbors of putative resistant cells. The same test as in **Fig. 7H** is used for two different groups: other (non-neighboring cells) and neighbors (neighbors of resistant cells among 4-h cells). ****: *p* < = 1*e* – 04. R. Cell-wise self-interaction Jacobian values of genes related to antiviral pathways, *NLRC5, TRIM21, JAK2, TRIM38, IFNGR1, TRIM26*, from 12-h cells for SARS-CoV-2 gene *S* on the UMAP embedding. S. A sigmoidal shape of the antiviral response that is sensitive to basal expression level may explain the putative mechanism of resistance to SARS-CoV-2 infection. [*X*_0_]: initial antiviral gene expression of background cells. [*X*_0_] + [*X*]: initial antiviral gene expression of resistant cells with higher-than-background expression. [*X*_1/2_]: half-maximal of antiviral gene expression. Δ*t*_1_: time delay between background and resistant cells because of initial difference in antiviral gene expression. Δ*t*_2_: time delay between background and resistant cells when expression achieves half-maximum.

## MATERIAL AND METHODS

### Cell Culture

HL60 cells (ATCCẋ CCL-240™) were grown in RPMI 1640 medium (Gibco), with 20% FBS + 5% Penicillin-Streptomycin) at 37°*C* under 5% CO2, supplemented with 10% fetal bovine serum (Sigma) and 1% Penicillin/Streptomycin (HyClone). Cells were maintained below a density of 10^6^ *cells/ml*. On the first day of the differentiation experiment, cells were seeded at 200, 000 *cells/ml* in 12well plates (unless stated otherwise) and treated with 1*μM* ATRA (all-trans-retinoic acid, Cat#R2625-100MG) to differentiate into either the neutrophil-like cells. Cell differentiation status was confirmed by flow cytometry analysis of *CD14* (Biolegend, Cat#367117) and *CD11b* (Biolegend, Cat#301309).

### scRNA-seq with 10x Chromium

HL60 differentiations were initialized on different days so that all samples could be harvested in a single scRNA-seq reaction to minimize batch effects. Cells were treated with 1 μM ATRA and differentiated for 0 (no ATRA treatment), 1, 2, 3, 4, or 5 days, with all differentiations performed in biological replicates. Samples were tagged or “cell hashed” (Stoeckius et al. 2018) with distinct BD sample tags (BD Bioscience, cat#PN 633780) to enable demultiplexing of cells, and then pooled for scRNA-seq. scRNA-seq was performed on one lane of the 10x Chromium™ Single Cell 3’ v2 system following the standard library prep protocol (10x Genomics Single Cell 3’ Reagent Kits v2 User Guide, CG00052). Libraries were amplified with 10 cycles of cDNA amplification and 15 cycles of Sample Index PCR. BD Sample Tags were size-separated by SPRI selection after cDNA amplification and amplified according to standard protocols (BD User-Demonstrated Protocol: BD Single-Cell Multiplexing Kit—Human Doc ID: 179682 Rev. 1.0). Final cDNA and sample tag libraries were sequenced on a NovaSeq 6000 (Illumina).

### scSLAM-seq

Our scSLAM-seq protocol was adapted from (Erhard et al. 2019; Hendriks et al. 2019). Before proceeding with the protocol using cells collected on particular days (see **below**), HL60 cells were labeled in medium with 100*mM* 4sU (Lexogen) for about 60 minutes at 37°*C* and sorted into lysis buffer (4*μl* 0.5 U/*μ*L Recombinant RNase Inhibitor (Takara Bio, 2313B), 0.0625% Triton X-100 (Sigma, 93443-100ML) in 96-well PCR plates. All plates were frozen at −80°*C* until use. After thawing the plates to room temperature, to the lysed cells, 0.4 *μL* of 10x PBS and 4.4*μL* of alkylation mix (20*mM* IAA in 100% DMSO) was added for a final concentration of 10mM IAA, DMSO. Alkylation was stopped by addition of 1.3 *μL* of 100*mM* DTT and incubating for 5 minutes at room temperature. Alkylated RNA was purified with 1.1 volume of Ampure XP beads and two washes with fresh 80% ethanol, and eluted into an RNA elution buffer (4*μ*L, 3.125 *mM* dNTP mix (Thermo Fisher, R0193), 3.125 *μM* Oligo-dT30VN (Integrated DNA Technologies, 5’AAGCAGTGGTATCAACGCAGAGTACT30VN-3’), 0.5 U/*μ*L Recombinant RNase Inhibitor, 1:24million ERCC RNA spike-in mix (Thermo Fisher, 4456740)). cDNA and the remaining library preparation was performed according to a modified version of the protocol for Smart-seq2 (Tabula Muris Consortium 2020). The prepared libraries were sequenced on MiSeq and NovaSeq5000 platform (Illumina), generating paired-end reads with 100 PCR-cycle.

### scSLAM-seq with sequential lineage tracing

To facilitate lineage tracing in scSLAM-seq libraries, cellular barcodes (GBCs) were introduced using a lentiviral transduction strategy (Adamson et al. 2016). Given that the success of this experiment critically depended on the uniqueness of barcode sequence to each cell at the start of the experiment, i.e. low barcode collision rate, and the capture of clone cells (clones with the same barcodes) across different days, we used an experimental scheme in which the starting population of the HL60 cells were infected at a low (2%) multiplicity of infection (MOI). This scheme has two benefits: first, we obtained a small number of barcoded single cells (~2000 in 1 ml of media in each well of a 24-well plate) so that we could capture clone cells via plate-based SLAM-seq (scRNA-seq augmented by metabolic labeling) characterized of low throughput; second, co-culturing the small number of infected cells with a large population of uninfected cells enabled us to differentiate infected cells more conveniently, as a small number of cells are difficult to grow and differentiate. Single cells carrying barcodes and expressing the blue fluorescent protein (BFP) reporter were sorted (Sony SH800) at five timepoints, days 0, 1, 2, 3, and 5, during differentiation in the presence of ATRA. cDNA from single cells was prepared in a 96-well format as previously described (Tabula Muris Consortium 2020) following alkylation and RNA cleanup (Erhard et al. 2019; Hendriks et al. 2019). Sequencing libraries were either reformatted into a 384-well format and prepared using TTP Mosquito automated liquid handlers, or in a 96-well format using a multichannel pipette. GBC sequencing libraries were prepared by dual PCR amplification to enrich for GBC cDNA and to add Illumina adapters and dual indexes complimentary to that cell’s transcriptome sequencing library indexes. GBC sequencing libraries were spiked into transcriptome libraries at 1:10 and sequenced on the NextSeq or MiSeq platform (Illumina). Transcriptome libraries were sequenced separately using a NovaSeq5000 S2 300-cycle kit.

### Dynamo: from velocity vector samples to continuous vector field functions and differential geometry analysis

Our analytical framework, ***dynamo***, consists of three integral stages: estimation of genomewide kinetic rate constants and velocity vectors, single-cell vector field functions with the resultant cell state and velocity samples, and various differential geometry analyses.

As the core of the first stage, we develop a comprehensive parameter estimation framework that includes all key steps involved in expression dynamics. This complete model assumes that the promoter of a gene stochastically switches with switching rate between an active state (*A*, with a high transcription rate *α_A_*) and an inactive state (*I*, with a much lower transcription rate *α_I_*) (Golding et al. 2005) Next, we explicitly model the accumulation or decay of 4sU-labeled RNAs (**Fig. 2A, B**, also see below), which are subsequently captured by scRNA-seq augmented with RNA metabolic labeling. We denote the ratio between the true 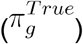 and estimated (*π_g_*) fraction of labeled reads for gene *g* as the **labeling correction coefficient**. Our model further incorporates RNA splicing dynamics with the splicing rate constant *β*. The degradation of the spliced RNA is captured by the degradation rate constant *γ_s_*. The protein translation rate constant *η* and degradation rate constant *γ_p_* are also modeled in ***dynamo*** for possible datasets from single-cell transcriptomic–proteomic coassays. For the purpose of simplicity, this work mainly focuses on RNA transcription, splicing, degradation, and metabolic labeling. We analyze various types of scRNA-seq data with and without metabolic labeling. For the former, we consider four possible experimental scenarios (**Fig. 2B**); for each case, one may or may not consider RNA splicing. We use three groups of models (**Fig. SI2C**) to describe these various types of scRNA-seq data. Details on how to estimate the RNA turnover rates and RNA velocities for each case are given below.

#### Limitations of conventional RNA velocity methods for scRNA-seq experiments without metabolic labeling

Most existing pseudotime ordering methods merely reveal the central trend of a population of cells. By contrast, RNA velocity (La Manno et al. 2018), an important recent development in inferring dynamics of single cells, explicitly models the RNA kinetics to offer a local extrapolation, for a period up to a few hours, of cell fate transitions of individual cells by exploring the intron or exon reads incidentally captured by most scRNA-seq platforms. The conventional RNA velocity method (La Manno et al. 2018) from the original paper exploits the kinetics of RNA transcription, splicing, and degradation with corresponding ODEs (ordinary differential equations) as follows:

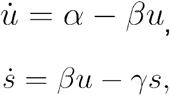

where *u* and *s* are the copies of unspliced and spliced RNA for a particular gene in a cell, respectively; *α, β*, and *γ* are the rate constants for transcriptional, splicing, and degradation (see **SUPPLEMENTARY METHODS** for a discussion of “rate” and “rate constant”, as well as their dimensions), respectively. In this study, we classify such a model system as **Model 1**. If we can estimate the kinetic parameters (*α, β, γ*), together with *u, s* measured by scRNA-seq, we can derive a measure of “RNA velocity” of unspliced 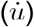 or spliced RNA 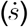 that reveals the direction and magnitude of rate of change of gene expression of each gene in each cell. Because in general *α* is not constant, but rather a function of the cell state and other variables (e.g., abundance of transcription factors, extrinsic signals, etc.), it is difficult to obtain the unspliced RNA velocity. On the other hand, splicing and degradation rate constants (*β, γ*) can in most cases be approximated as constants for certain cell types. The question, then, is how to estimate those kinetic parameters. Assuming pseudo-steady state 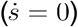 for cells with extreme high unspliced and spliced RNA expressions (top right corner of the phase plane), one reaches the following linear relation between the spliced and unspliced RNA

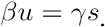

Let 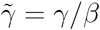, the above relation can be rewritten as:

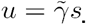

A linear regression of cells at steady states can be performed to obtain 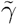. Thus, the conventional RNA velocity as defined in the original study is given by:

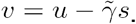

Note that *υ* is equal to 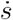 up to the splicing rate constant *β*, which is in general gene-specific as revealed in **Fig. 2D**. Because velocity can be estimated for each gene in each cell, velocities of all genes in any cell form a high-dimensional vector, with each dimension corresponding to a gene. This high-dimensional velocity vector is often projected into a low-dimensional space for visualization using either pearson or cosine kernels (La Manno et al. 2018; Bergen et al. 2020; Li et al. 2020) to reveal the direction of cell fate transitions in low-dimensional space via projected velocities.

Although conventional RNA velocity has been successfully applied to a variety of studies, it has several limitations:

1. Because the intron reads are generated through mis-priming on polyA- or polyT-enriched intronic regions of nascent pre-RNA, conventional RNA velocity can be difficult to apply to most transcription factors, which are typically expressed at low levels, and genes with no polyA/T-enriched intron regions;
2. The linear regression methods used by conventional RNA velocity ignores the distribution of unspliced and spliced RNA, which can be used to improve the estimation of kinetic parameters;
3. For systems far away from the pseudo-steady state, using cells with extreme RNA expression levels for linear regression may lead to inaccurate velocity calculations for most cells;
4. The time scale for conventional RNA velocity 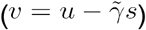 is scaled by *β* (since 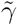 is the ratio of *β* and *γ*), which makes velocity a relative quantity.
5. Conventional RNA velocity only estimates velocity for observed cells. Thus, it is a discrete, sparse, and local measure of cell dynamics and often merely used as a descriptive instead of a predictive tool.

A great deal of effort has been devoted to improving conventional RNA velocity estimation (La Manno et al. 2018) in regard to challenges 2) and 3) and extend the concept to “protein velocity” (Gorin, Svensson, and Pachter 2020), but 1) and 4) are fundamental limitations that cannot be resolved at the computational level without additional experimental information. In this section, we introduce our methods for analyzing conventional scRNA-seq data, addressing some of the issues with existing RNA velocity methods. In the next section we focus on computational methods for computing RNA velocity for metabolic labeling data, which reconciles the splicing- and labeling-based kinetics and overcomes other drawbacks of conventional RNA velocity methods. Finally, to address 5), we go beyond RNA velocity samples of single cells to map the continuous vector field functions in transcriptomic space and perform sophisticated differential geometry analyses to gain various functional vector field predictions and biological insights.

#### Generalized method of moments (stochastic splicing and negative binomial distribution method) improves RNA velocity estimation for conventional scRNA-seq experiments

Current scRNA-seq methods have low RNA capture rates that lead to frequent “dropouts,” in which individual RNA levels are not observed. In order to alleviate dropout effects and measurement noises as well as to improve the robustness of the estimation, the original RNA velocity method (La Manno et al. 2018) utilizes the mean expression (first moment) of each gene across cells, calculated based on the *k*-nearest neighbor graph of cells, instead of the raw expression:

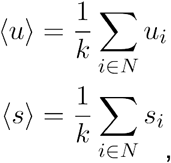

where *N* (30 by default in ***dynamo***) is the set of *k*-nearest neighbors of each individual cell, often constructed in the space of the top PCs (principal components) (e.g., 30 PCs), reduced from the original gene expression space of highly variable genes. These can be considered as estimators of the first moments of the distribution of unspliced and spliced RNAs. RNA velocity calculations performed on the first moments lead to a cleaner phase plane and therefore smoother velocity vectors (La Manno et al. 2018). However, higher moments of the distribution are ignored in the original linear regression method.

Second moments (uncentered variances and covariances) provide information additional to first moments on the shape of the underlying distribution. It is thus desirable to also take advantage of the second moments to improve the estimation robustness and accuracy of the kinetic parameters, and thus that of the RNA velocity measurements. The second moments of unspliced and spliced RNA, as well as their mixed moments, also rely on the *k*-nearest neighbor graph of cells, and can be computed as follows:

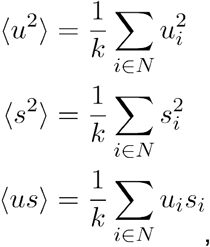

With the first, second, and mixed moments of unspliced and spliced RNAs for each gene across cells, one can apply the generalized method of moments (GMM) to improve the estimation of kinetic parameters *θ* (e.g. *α, β*, and *γ*), in lieu of the linear regression on mean expressions as used in the original RNA velocity method. Instead of directly fitting the distribution, GMM seeks to solve the following equations of moments for *θ*, also known as *moment conditions*:

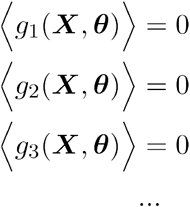

where *g*_1_, *g*_2_, *g*_3_,… are functions of the random variables ***X*** (e.g. the copies of spliced and unspliced RNA across cells) and parameters *θ*. The optimal *θ* can be found by minimizing the Euclidean norm of the above expectations:

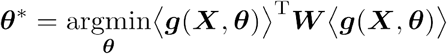

where ***g*** is a vector-valued function consisting of the moment conditions, and ***W*** is a positive definite weighting matrix.

Specifically, to apply GMM in the context of RNA velocity, one needs to find the moment conditions for first and second moments. By deriving the ODEs for first and second moments from master equations of **Model 1** (**Fig. SI2B**), Berger et al. showed that the moment conditions are (Bergen et al. 2020):

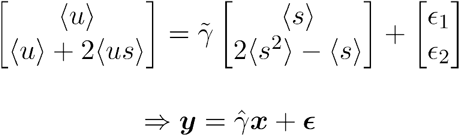

where 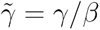. Given vector pairs 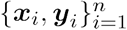 of the first and second moments computed from the conventional scRNA-seq data in *n* cells at pseudo-steady state, the optimal 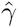 is obtained by minimizing the following least squares:

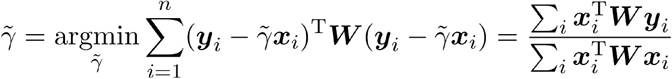

where ***W*** is the covariance matrix and is computed as follows:

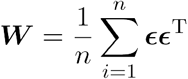

We name this procedure as the *stochastic splicing method*, which has been proven to be more accurate and robust than the original linear regression method used in the conventional RNA velocity, possibly due to the inclusion of the additional moments (Bergen et al. 2020). Another major improvement to the RNA velocity methods from (Bergen et al. 2020) is the *dynamical model*, where Berger et al. derived the solutions for *u* and *s* under the assumption that the promoter has only two states: active and inactive. This assumption is reasonable and proven to be effective but not necessarily true; see above discussion of transcription rates. An EM algorithm is used to iteratively infer the state of the promoter and the latent time for each gene in each cell, and then the solutions are fit to the resulting pseudo-time course of unspliced and spliced RNAs to obtain the kinetic parameters. No steady state assumption is required in this method other than providing a reasonable guess about the initial values for kinetic parameters.

We also developed an alternative procedure, the *negative binomial (NB) distribution method*, based on an observation that in most cases total RNA counts at steady state follow the NB distribution (Grün, Kester, and van Oudenaarden 2014). With this distribution the variance *σ*^2^ (second central moment) and the mean *μ* satisfy the following relationship:

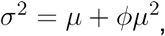

where *ϕ* is the reciprocal of the dispersion parameter of NB distribution. Because at the pseudo-steady state, the first moment of spliced and unspliced RNA is related by 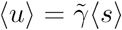 so the spliced RNA is proportional to the total RNA, making it an NB-distributed variable. The variance of spliced RNA satisfies:

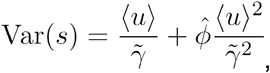

where 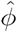 is the estimator of *ϕ* and is computed from:

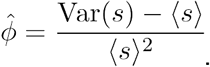

Put all together, these give the moment conditions for the first and second moments:

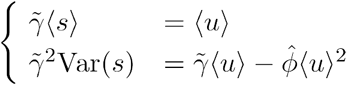

A nonlinear least squares optimizer can then be used to solve for 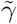 with the above two equations. Note that the two assumptions applied here are: 1) there is a linear relationship between two random variables, which are not limited to the unspliced and spliced RNA, but can also be generalized to labeled or new and total RNA, and 2) one of the variables follows the NB distribution. Therefore, it is straightforward to generalize this method to one-shot labeling data, as will be detailed later.

#### Estimating absolute RNA velocity for metabolic labeling-based scRNA-seq experiments across various labeling strategies

Because metabolic labeling–based scRNA-seq (time-resolved RNA-seq or tscRNA-seq) measures the synthesis or degradation of labeled RNA within a known period of time in an experimentally programmable manner, it offers a more direct measurement of the kinetics of gene expression than cscRNA-seq. Thus, in principle, it also provides an opportunity to overcome some of the challenges of the cscRNA-seq in RNA velocity estimation. However, it is nontrivial to properly estimate kinetic parameters and compute RNA velocity for tscRNA-seq data with various metabolic labeling approaches, including three general labeling strategies given in **Fig. 2C**: one-shot (the simplest labeling strategy with a single RNA labeling period), kinetics or pulse (a time-series of 4sU or other nucleotide analog treatment to observe the accumulation of metabolically labeled RNA over time), and degradation or chase (a time-series after an extended 4sU or other nucleotide analog treatment period, followed by chase at multiple time points after the wash-out to observe the decay of metabolic labeled RNA over time). Although the exact details of the resultant data vary across different labeling strategies, we found they can be uniformly treated with two different models, **Model 2,** which explicitly considers RNA labeling but not splicing, and **Model 3**, which considers both labeling and splicing (**Fig. SI2B**). In the following, we will first briefly introduce these two models, then provide the respective estimation procedures of the three general labeling strategies based on the corresponding models.

In **Model 2**, we take into account labeling (with a labeling correction coefficient A) but not splicing. The total RNA has a synthesis rate constant *α* and a degradation rate constant *γ*. The labeled RNA has a reduced synthesis rate constant λ*α* but the same degradation rate constant. The ODEs for describing the dynamics of labeled (*l*) and total (*r* = *l* + *o*) are,

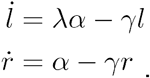

The general solution for the total RNA *r* over time *t* is:

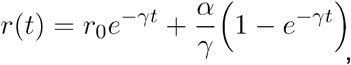

where *r*_0_ is the initial concentration of the total RNA *r*. For the labeled RNA, the solution is:

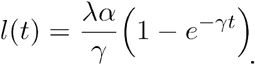

Note that in this study we rely on a binomial mixture distribution model of background or 4sU-introduced mutation rates, otherwise stated, to quantify the labeled or unlabeled RNA from the observed T-to-C mutation in the final sequencing reads (Jürges, Dölken, and Erhard 2018). Therefore, assuming labeled RNA (*l*) is well corrected with the binomial mixture model (Jürges, Dölken, and Erhard 2018), A will is effectively 1 (same as in the following). Also see **SUPPLEMENTARY METHODS** for a detailed discussion on labeling correction coefficient.

In **Model 3**, we consider both the labeling and the splicing processes. The solutions for labeled, unspliced RNA (*u_l_*) and labeled, spliced RNA (*s_l_*) are equivalent to those for unspliced and spliced RNA in **Model 1**, with an additional A modifying the effective transcriptional rate of the labeled RNA:

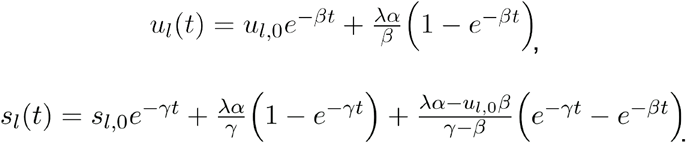

When *β* = *γ*, the solution for *s_l_* is instead:

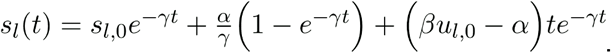

We will omit this special scenario for simplicity in the following sections, although it is included in ***dynamo*** for the sake of completeness and robustness for kinetic parameter estimations.

Below, we detail the respective estimation procedures of the four labeling scenarios given in **Fig. 2B** based on the corresponding models.

**Table 1.** Available estimation algorithms for each labeling strategy.

| Labeling strategy | One-shot | Kinetics (pulse) | Degradation |
| --- | --- | --- | --- |
| Model | Model 2/3 | Model 2/3 | Model 2/3 |
| Has splicing | With or without | With or without | With or without |
| Time points | Single time point | Multiple time points | Multiple time points |
| Steady state assumption | Yes | Yes or No | Yes or No |
| Estimation | “One-shot” method (without splicing); NB method (with or without splicing); | “Two-step” method (without splicing); NB method (with or without splicing); curve fitting (with or without); | Curve fitting (with or without splicing) |

**Table 1: Available estimation algorithms for each labeling strategy.**
| Velocity | Velocity_N/T/S/U if integrated with conventional RNA velocity, Velocity_N/T otherwise | Velocity_N/T/S/U if integrated with conventional RNA velocity, Velocity_N/T otherwise | Velocity_S if splicing is considered, none otherwise |
| --- | --- | --- | --- |

Now we will introduce the respective estimation procedures and the corresponding models for each of the three general labeling strategies given in **Fig. 2B**.

#### One-shot experiment

In “one-shot” experiments, there is only one labeling time point, and the splicing process is not explicitly considered. The solution for new RNA in **Model 2** is:

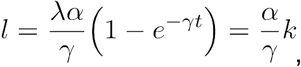

where *t* is the labeling time and we denote *k* = λ(1 − *e*^−*γt*^). When the dynamics of total RNA is at steady state 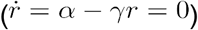,

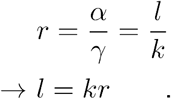

Then the parameter *k* can be obtained through a simple linear regression with zero intercept of the first moments of labeled and total RNAs (*l, r*), for cells with extreme high expressions of both *l, r* (top right corner of the phase plane). This approach effectively replaces *u, s* in the original RNA velocity method with *l, r*, and was previously reported as the “NTR” (New to Total Ratio) velocity method (Erhard et al. 2019). Although not fully explained in the literature, the NTR velocity can be calculated as:

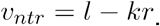

Because we used corrected labeling RNAs, i.e. λ ~ 1, the degradation parameter *γ* can be calculated from *k* and the labeling duration *t*:

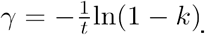

Because we obtain *γ*, not the relative 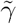 as in the original velocity of spliced RNA, we can calculate the velocity of total RNA with a physical time unit (Q. Qiu et al. 2020):

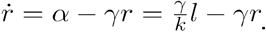

Note that the NTR velocity proposed in (Erhard et al. 2019) is very similar to this method, but scaled by *γ/k*, a factor that can differ for individual genes and cancels the unit of time, so it only approximates the true kinetics.

Because in one-shot experiments the labeled and total RNAs are linearly correlated with a slope of *k* = λ(1 − *e*^−*γt*^), and at steady state the total RNA follows the negative binomial distribution, one can easily incorporate second moments using the negative binomial method:

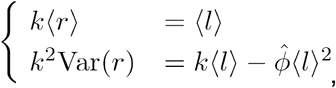

where,

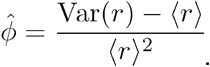

Then one obtains a more accurate slope *k*, and can be used to compute the velocity of total RNA.

#### Kinetics (pulse) experiment

Two approaches were developed to estimate the RNA turnover rates for the datasets obtained from the kinetics experiment. The first method is a generalization of the “one-shot” method to multiple time points, whereas the second uses a curve fitting strategy which can be also applied to datasets obtained for the degradation experiment. We introduce these two approaches in order:

##### 1) The “two-step” approach (**Fig. 2C** Case 2-4, multi labeling time points/with or without splicing)

With data collected at multiple labeling time points in a kinetics (pulse) experiment, on the phase plane of labeled and total RNA, we find that cells from the same labeling period are distributed on a line whose slope increases as the labeling period increases. We realize that this phenomenon can be explained by the fact that the slope *k* is a monotonically increasing function of the labeling time *t* (see the **“one-shot” method**):

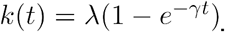

We then take advantage of this discovery and develop the “two-step” approach, which relies on two consecutive linear regressions to estimate the degradation rate constant *γ* based on **Model 2 (Fig. SI2B)**, and the steady state assumption that 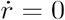. The first step computes the slope *k* for the labeled (*l*) and total (*r*) RNA for different labeling time *t*, based on the linear relationship (see the **“one-shot” method**):

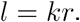

When labeling correction coefficient is close to one, from *k*(*t*) = 1 − *e*^−*γt*^, it is apparent that the slope increases with longer labeling time and asymptotically approaches one. Rearranging this equation, we have:

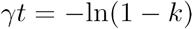

A linear relationship exists between the labeling time *t* and the quantity −ln(1 − *k*). In the second step, we then estimate the parameter *γ* using a simple linear regression of *t* and −ln(1 − *k*). The total RNA velocity is again:

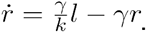

Note that the “two-step” approach can be regarded as a generalization of the above “one-shot” method for one-shot labeling experiments to kinetics experiments with multiple labeling time points. The negative binomial method can also be applied here in the first step to achieve a more robust estimation of the slope *k*. We note that not every single gene in the dataset may follow this kinetics, and in general we use R-square of the “two-step” model fitting to select genes with confident fittings for downstream analysis.

##### 2). Curve fitting methods (**Fig. 2C** Case 2-4, multi labeling time points/with or without splicing)

When single-cell kinetics (pulse) or degradation (chase) data using RNA metabolic labeling [e.g., scEU-seq or scNT-seq (Battich et al. 2020; Q. Qiu et al. 2020)]), at multiple time points are available (**Fig. 2B Case 2**), it is possible to estimate the kinetic parameters (i.e. *α, β, γ*) for each gene using nonlinear least-squares methods. In general, given *m* experimental data points *y*^(1)^, *y*^(2)^,…, *y*^(*m*)^, at time points *t*^(1)^, *t*^(2)^,…, *t*^(*m*)^, the least-squares fitting method finds a set of parameters *θ* that minimize the following loss function:

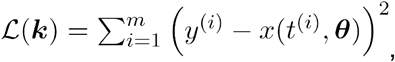

where *x*(*t*, ***θ***) is the solution of the ODEs at the time point *t*, given parameters *θ*. When there are multiple species (i.e., unspliced labeled *u_l_*, spliced labeled *s_l_*, unspliced unlabeled *u_u_*, or spliced unlabeled *s_u_* RNAs) quantified from the experiment, we cast the ODEs into a matrix form while the composite loss function is the summation for loss function of all species, and weights can be added to the loss function to adjust the importance of each species (e.g., a higher weight is assigned to the labeled than the unlabeled species (default is 2:1) for the kinetics experiment because the unlabeled species does not strictly follow the degradation kinetics due to imperfect labeling):

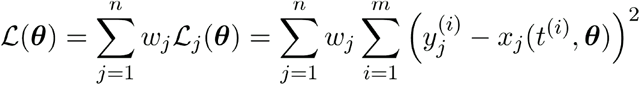

This general procedure is applied to all following curve-fitting methods; the key is to find solutions of each species for various RNA labeling strategies.

We used Latin hypercube sampling to randomly initialize a set of values of *θ* in a predetermined range (see **SUPPLEMENTARY METHODS**) as the initial guesses for the parameters *θ* required by the nonlinear least squares optimizer.

In kinetics experiments, the samples are collected after a short period of 4sU (or other nucleotide analogs) labeling. At the beginning of the experiment, the concentrations for labeled RNA, unspliced labeled and spliced labeled RNA, are zero (*l*_0_ = *l*(0) = 0, *u*_*l*,0_ = *u_l_*(0) = 0, and *s*_*l*,0_ = *s_l_*(0) = 0). During the labeling process, because we assume that the labeling period is much shorter than the time scale of the biological process of interest, transcriptional rates are treated as constant in all cells. Therefore, based on the solutions of **Model 3,** the abundance of labeled, unspliced labeled and spliced labeled RNA increase over time:

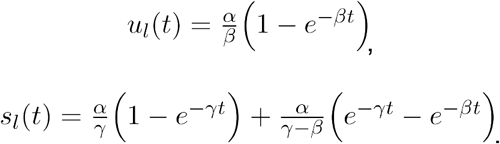

With sufficient sampling of the labeling time points (at least three), all three kinetic parameters can be estimated in theory. Because cells at different states may have different transcription rates, clustering can be performed first and the fitting is done for each cluster to derive cluster or cell-type specific kinetic rates (Battich et al. 2020; Q. Qiu et al. 2020). The above solutions are often insensitive to variations in 7, and the read counts for the unspliced RNA are unreliable for genes with fast splicing rates, so it is optional to provide further constraints by including the kinetics of unlabeled or old, unlabeled spliced and unlabeled unspliced RNA, in the curve-fitting procedure. The unlabeled RNA in kinetics experiments mostly follow the degradation kinetics, if the labeling efficiency is close to 1 (see **SUPPLEMENTARY METHODS**), and the solutions are more sensitive to *β* and *γ* than those of the labeled species:

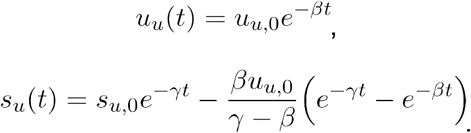

The spliced RNA velocity can be computed as before:

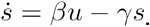

This also allows us to compute the velocity for unspliced RNA in individual cells:

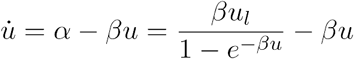

If no splicing data are available, the solution for **Model 2** can be used:

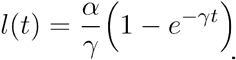

The total RNA velocity can be computed either for each cluster, where *α_c_* denotes the transcription rate constant of cluster *c*:

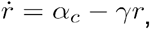

or for individual cells:

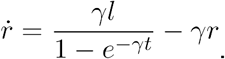

The velocity for new RNA can be computed in a similar way:

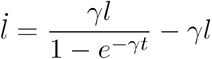

There is, however, a practical issue when using curve-fitting methods with **Model 1** for data obtained from the kinetics experiments. Because the current labeling time of a tscRNA-seq kinetics experiment typically requires at least 1 hour (because of the low sensitivity of singlecell methods), which is much longer than the time scale of RNA splicing (usually on the scale of minutes), the labeling kinetics do not have sufficient time resolution for reliable estimation of the splicing rate constant *β*. We can circumvent this by first computing 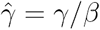 from the total unspliced 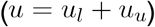 and spliced RNA (*s* = *s_l_* + *s_u_*) using the conventional RNA velocity method. Then, we can use either model to estimate the actual degradation rate constant *γ*, and the splicing rate constant is simply given by:

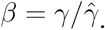

With this, we can then estimate absolute RNA velocities for total, spliced, unspliced, and new RNAs according to the model and data available. Note that a similar procedure can also be applied to relative kinetic parameters estimated with the dynamical method from (Bergen et al. 2020) that generalize to non–steady-state assumption, and used to scale them to absolute values.

#### Degradation (chase) experiments

In degradation experiments (**Case 3 in Fig. 1B**), samples are chased after an extended 4sU (or other nucleotide analog) labeling period and the wash-out to observe the decay of the abundance of the (labeled) unspliced *u_l_* and spliced *s_l_* RNA decay over time. The process can be formulated as below (the zero in the subscript indicates the initial condition):

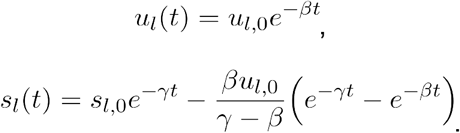

These two equations can be substituted into the loss function, and we obtain splicing rate constant *β* and degradation rate constant *γ* using the nonlinear least squares. The (labeled and unlabeled) spliced RNA velocity is then given by:

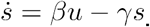

Although the unlabeled RNAs (*u_u_, s_u_*) indeed increase over time due to transcription, cell-wise transcriptional rates *α* cannot be directly estimated from such experiments because each cell has different transcription activity. However, with a promoter two-state stochastic expression model, we can assume a universal *α*_on_ and *α*_off_ for all cells, similar to the dynamical model (Bergen et al. 2020).

For degradation experiments without splicing data, the solution of **Model 2** is used. The abundance of labeled RNA (*l*) follows the first-order decay kinetics (Q. Qiu et al. 2020):

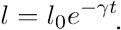

Note that this method has the same drawback as the curve-fitting method for experimental kinetics data, i.e., the estimation of *β* can be unreliable if the chasing time resolution is much larger than the time scale of splicing. Again, one may combine the curve fitting with the conventional RNA velocity method and obtain a more accurate splicing rate constant *β* and RNA velocities.

#### Robust reconstruction of continuous velocity vector field functions from sparse single cell transcriptomic measurements

In the second and third stages of our ***dynamo*** model framework, we robustly learn a continuous vector field function of single cells from the input discrete, sparse, and noisy single-cell velocity vector samples. We also bring in predictive dynamical system methods and differential geometry analyses to improve the interpretability of the “black box” machine learning powered vector field functions, thus marrying the power of advanced machine learning (ML) approaches in functional approximation with the interpretability of dynamical systems formulations.

##### Vector field of expression space in single cells

In classical physics, including astronomy, fluidics and aerodynamics, velocity and acceleration vector fields are used as fundamental tools to describe motion or external force of objects, respectively. In general, a vector field can be defined as a vector-valued function ***f*** that maps any point (i.e. expression state of a cell) ***x*** in a (subset of) *d* dimensional (gene expression) space to a vector ***υ*** (e.g. the RNA velocity vectors) in the same space, i.e., ***υ*** = ***f***(***x***). Thus, RNA velocity estimates (La Manno et al. 2018; Bergen et al. 2020) from single cells can be formally treated as samples in the velocity vector field. In two or three dimensions, a vector field is often visualized as a quiver plot, where a collection of arrows with a given magnitude and direction is drawn. Assuming an asymptotic deterministic system, the trajectory of the cells travelling in the gene expression space follows the vector field and can be calculated using numerical integration methods, e.g., the Runge–Kutta algorithm. In two or three dimensions, a streamline plot can be used to visualize those integration paths. For high-dimensional vector fields, it is challenging to present all information at once, and multiple quantities are required to reveal different features of the vector field. As we will show later, differential geometry offers many such quantities, each allowing us to capture some but not all dynamical features of the vector field.

##### Vector field reconstruction from sparse, noisy single-cell expression and velocity samples

With csc- or tscRNA-seq data and the computational framework mentioned above, in principle we can obtain vector field samples in either the unspliced, spliced, new, or total RNA space, depending on the exact experiment, labeling strategy, and estimation method. High-dimensional velocity vectors are often projected onto top PCA (principal component analysis) space or two- or three-dimensional UMAP (Uniform Manifold Approximation and Projection) space (La Manno et al. 2018; Bergen et al. 2020). In order to go beyond sparse velocity samples to continuous vector field functions in full gene expression space, we build on some recent advances in vector valued function approximation to scalably, efficiently, and robustly learn the transcriptomic vector field (see **Box 2** and **below**) from noisy and sparse samples of single-cell states and velocity estimates. Our reconstruction works in projected PCA or UMAP space, or even in the full geneexpression space. When it is reconstructed in low-dimensional space, the learned vector field can be projected back to the original transcriptomic space for gene-specific velocity and differential geometry analyses.

##### Vector Field Reconstruction in the Reproducing Kernel Hilbert Space

To formally introduce the problem of velocity vector field learning in the context of scRNA-seq, we consider a set of pairs of cell expression states 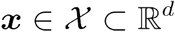 and RNA velocities 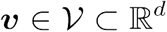, i.e. 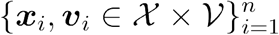, where *n* is the number of cells, and *d* is the dimension (i.e. number of genes or number of principal components) of the cell state space. We suppose that the measured single-cell RNA velocity is sampled from a smooth, differentiable vector field that assigns each cell expression state ***x*** with a RNA velocity vector ***υ***. Normally, single-cell RNA velocity measurements are results of biased, noisy, and sparse sampling of the cell expression state space. Therefore, the goal of velocity vector field reconstruction is to robustly learn a mapping function ***f***, which outputs an RNA velocity vector ***υ*** given any cell expression state ***x***, based on the observed data 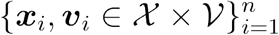, under certain smoothness constraints (J. Ma et al. 2013). Ideally, the mapping function ***f*** should recover the true velocity vector field on the entire domain 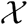 and can be used to predict the true dynamics in regions of expression space that are not sampled. The discussion introduced above is based on velocity vector field but it can be similarly extended into any general vector field, e.g., an acceleration vector field (Gorin, Svensson, and Pachter 2020).

Intuitively, the loss function for the search of an optimal vector field function ***f***\* can be written in a least-squares fashion:

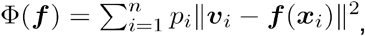

where *p_i_* is a weight deciding the importance of the *i*-th data point in the loss function. However, It is not a trivial task to minimize the above loss function with respect to a function ***f***. Approximating vector-valued functions in a sparse reproducing kernel Hilbert space (RKHS) has been shown to be effective in learning vector field functions for 2D applications, and can be easily generalized to high dimensional data (J. Ma et al. 2013). For a function in the RKHS space, i.e. 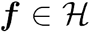, The function can be evaluated at any point in 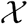, as a summation of Gaussian kernels centered on the so-called “control points”:

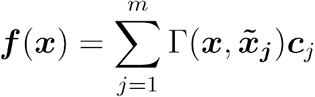

where *m* is the number of control points and 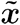 is the coordinate of the control point. ***c***’s are coefficient vectors in 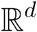, where *d* is the dimension of the vector field. The reproducing kernel is chosen to be a Gaussian function:

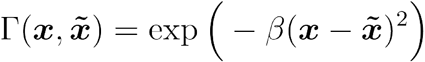

where *β* is a width parameter. In addition, a norm of functions can be computed on 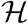 (J. Ma et al. 2013):

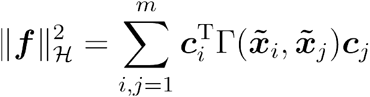

In this representation, the loss function can be optimized with respect to the coefficient vectors ***c***, and a vector-valued *L*_2_ regularization term can be introduced to it:

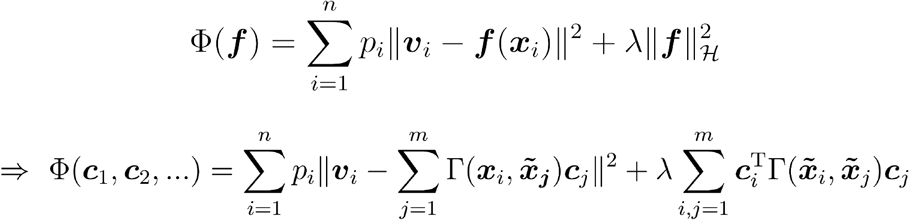

The sparseVFC (sparse vector field consensus) algorithm (J. Ma et al. 2013) improves this loss function for better outlier identification and rejection by formulating the weight *p_i_* as a likelihood function (See details in **SUPPLEMENTARY METHODS**). The final loss function has an additional parameter *σ* accounting for inlier noise:

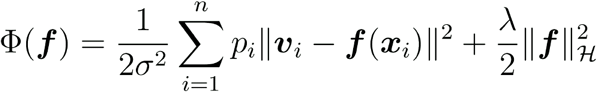

If we let ***C*** = [***c***_1_, ***c***_2_,…, ***c**_m_*]^*T*^, it can be shown that the solution ***C***\* to the following linear equation contains the coefficient vectors for the optimal vector field function ***f***\*:

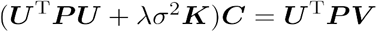

where *U* is an *n*-by-*m* matrix whose elements are 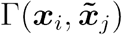 and ***K*** an *m*-by-*m* Gram matrix consisting of 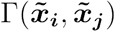. The ***P*** matrix is a diagonal matrix of the weights *p_i_*, and ***V*** = [***υ***_1_, ***υ***_2_,…, ***υ***_*n*_]^T^.

The sparseVFC algorithm (J. Ma et al. 2013) consists of 1) an **E-step**: calculation of the diagonal matrix ***P*** based on the likelihood function for outlier rejection (see **SUPPLEMENTARY METHODS**), and 2) an **M-step**: Solving the above linear system for ***C***, and updating the vector field function evaluations at sample points ***f***(***x***) with the optimal ***c**_i_*’s. Other parameters, for example *σ*, are also updated accordingly in this step. The algorithm finishes when the loss function converges or the number of optimization steps surpasses the designated maximum iterations.

##### Topological analysis of single-cell vector field

In this study, we focus on calculating fixed points and nullclines in our topological analysis of vector fields. The fixed points are defined as points where the value of the vector field function is zero:

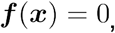

and the solution can be obtained using any nonlinear equation solver (scipy fsolve is used in our case). Because the solver can only find fixed points closest to an initial guess ***x***_0_, we simply randomize *n* such initial points in a domain containing all data points. We used Latin hypercube sampling technique (Iman, Helton, and Campbell 1981) to sample initial points effectively. To characterize the stability of a fixed point, the Jacobian is evaluated at the point and we simply categorize fixed points into three types, based on the signs of its Jacobian’s eigenvalues:

1. Stable fixed point (attractor): all eigenvalues are negative;
2. Unstable fixed point (repulsor): all eigenvalues are positive;
3. Saddle point: The eigenvalues are a mixture of positive, negative values, or even zero.

If one is interested in fixed points of a specific order (i.e., with a given number of positive eigenvalues), one may use a quasi-Newton conditional root-finding algorithm developed by Wang et al. (P. Wang et al. 2014).

Nullclines are lines (in 2D) or surfaces (in higher dimensions) when at least one component of the vector field is zero. For example, for a 2D vector field ***f***(*x, y*) = [*f_x_*(*x, y*), *f_y_*(*x, y*)]^T^, the x-nullcline consists of points where:

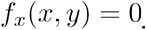

Because it is computationally expensive to compute nullclines in higher dimensions, in our study we limit to 2D vector fields. In the 2D case, fixed points are intersections of x- and y-nullclines, so we compute nullclines using a pseudo-arclength continuation method (Seydel 1988) starting at a certain fixed point. As an example, to find the next point ***p***_1_ on the x-nullcline, given a known point ***p***_0_ and a tangent vector of the nullcline ***υ***_0_, one simply finds the initial guess for ***p***_1_ by:

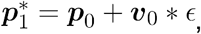

where *ϵ* is an incremental increase in the arclength. ***p***_1_ can then be found using a nonlinear equation solver for:

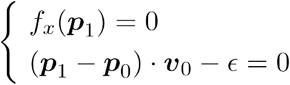

This guarantees that the solution ***p***_1_ is *ϵ* away from the known point ***p***_0_ on the nullcline. The tangent vector for the next iteration is approximated as ***υ***_1_ = (***p***_1_ – ***p***_0_)/|*p*_1_ – ***p***_0_|, and the first tangent vector at the fixed point is a random normalized vector.

##### Differential geometry analysis of the reconstructed single-cell vector field

We derive the analytical formula of Jacobian of the vector field which improves the computational efficiency tremendously than numerical approaches. The vector field function obtained from the sparseVFC algorithm has the following form (See Box 2 for details):

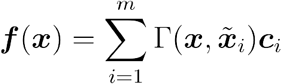

where Γ is the Gaussian kernel, 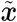 are the control points, and ***c*** are the combination coefficient vectors. Because the vector field is a linear combination of Gaussian kernels, whose derivative is:

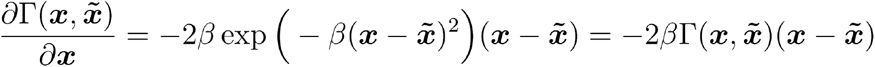

The Jacobian of the vector field function is then:

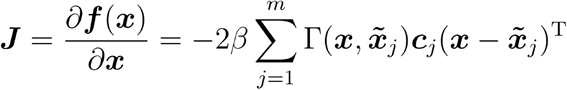

Let:

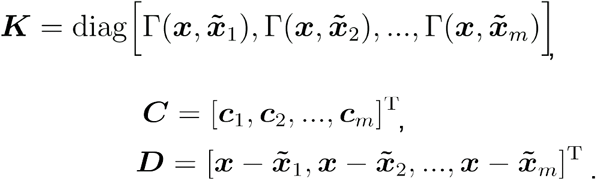

Then, the above analytical form of the Jacobian can be vectorized into:

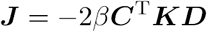

The divergence, acceleration, and curvature are calculated as follows, respectively:

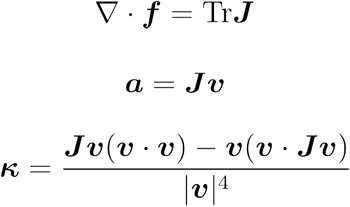

In 2D, the curvature formula has an equivalent but simpler form:

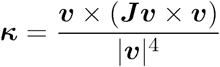

where ***υ*** is the velocity vector. The curl is only computable in 2D or 3D, and is computed as follows for a 3D system:

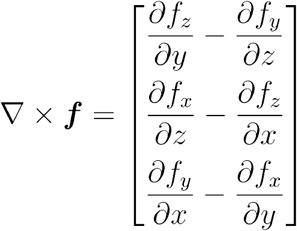

Because the vector field function is often learned in a PCA-reduced space, and to acquire genespecific information, a transformation of the Jacobian from the PCA space to the original gene expression space is needed. Suppose the first *k* principal components form a *d-by-k* matrix ***Q***, where *d* is the dimension of the original gene expression space, then the gene-specific Jacobian ***G*** is:

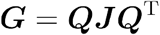

Thus, the *ij*-th element of ***G*** is the partial derivative of the velocity of gene *i* with respect to the expression level of gene *j*. The obtained Jacobian ***G*** here is only an approximation of the true gene-specific Jacobian, as only *k* < *d* principal components are used.

#### Vector field simulation and benchmark of the two-gene bifurcation system

We use the simple canonical self-activating and mutual-inhibiting two-gene motif that frequently appears in a variety of cell fate bifurcation systems to introduce key concepts in dynamical systems and differential geometry employed in this study (**Fig. 1A**). The vector field function of this system is adapted from Qiu et al. (X. Qiu, Ding, and Shi 2012):

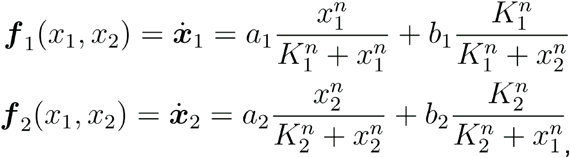

where *K*_1_ = *K*_2_ = *a*_1_ = *a*_2_ = *b*_1_ = *b*_2_ = 1, *n* = 4. In the following two subsections, we will describe how the demonstration of the vector field analysis and the benchmarking of our vector field reconstruction with this two-gene system are performed.

##### Mapping the topological and geometry feature of the two-gene system

To make the quiver plot of the two-gene system, we first set the expression range of *x*_1_, *x*_2_ to be [0,2.5] and plot the velocity values calculated with the above vector field function on a 25-by-25 grid with even spacing in this space. The velocity values on the grid are also used to create the streamline plot. Individual trajectories associated with states 1, 2, 3 are obtained via numerical integration of the vector field function. Attractor and saddle points are directly solved from the vector field function. To obtain the separatrices, we start with unstable fixed points, calculate the eigenvalues at those points, and then move forward a tiny step along the direction of the negative of the eigenvalues. In the end, we integrate the vector field function backwards in time to generate the separatrices. Next we calculate the analytical Jacobian of this system:

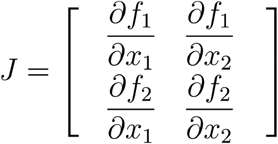

Because our model system is symmetric, we only need to calculate, for example, the elements from the first column, denoted in the following:

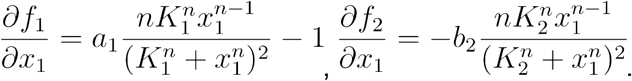

With the analytical Jacobian, we can then also obtain the analytical formula for the acceleration, curvature, divergence, and curl. We use heatmaps to plot the four elements of Jacobian, divergence, and curl on the same space used in the quiver or streamline plot. For acceleration and curvature, because they are vectors like velocity, we thus plot their magnitudes together with the corresponding vector fields (i.e., acceleration or curvature vector field). To enhance the presentation of the plottings for the above differential geometry quantities, higher than 25×25 grids, 2-D gaussian kernel smoothing on those grid values as well as different colormaps are used as needed.

##### Benchmarking the reconstruction of the vector field and the calculation of differential geometry quantities

To generate the benchmark dataset, we randomly select 5000 points within the same domain used in the above plots and calculate the corresponding velocity vectors for those points. Those cell states and velocity vector pairs are then used as inputs to reconstruct vector field function with ***dynamo*** using default parameters. Attractor, saddle points, and nullclines are estimated with the reconstructed vector field function and plot with the streamline plot that is also based on the reconstructed vector field function via ***dynamo***. We used the reconstructed vector field function to calculate analytical Jacobian, acceleration, curvature, curl, and divergence with ***dynamo***. Scatterplots from ***dynamo***, including a frontier showing the boundary of all those cells, are used to plot the 5000 sampled cells, colored according to either four elements of the Jacobian, divergence, or curl at those points. ***Dynamo*** is also used to estimate the acceleration and curvature for those sampled cells, and then plot their magnitudes together with the corresponding vector fields (i.e. acceleration or curvature vector field). We calculate the analytical Jacobian, acceleration, curvature, divergence, and curl with the true vector field function at those sampled data points and compare the corresponding values estimated from ***dynamo*** with scatterplots (**Fig. 4C–E**).

To demonstrate the efficiency of our vector field reconstruction, we compare the time used either by the numeral approaches that build upon the [numdifftools](https://pypi.org/project/numdifftools/) or by the analytical approaches, both implemented in ***dynamo***. Note that numerical approaches for those differential geometry quantities are only possible with the analytical vector field function we learned, especially in the high-dimensional gene expression space.

#### Estimating RNA velocity for SARS-CoV-2 genes

SARS-CoV-2 is a positive-sense, single-stranded RNA virus (D. Kim et al. 2020). As an RNA virus, SARS-CoV-2 does not have introns; thus, we cannot rely on RNA splicing and degradation to estimate RNA velocity for SARS-CoV-2 viral genes. Upon cell entry, two open reading frames (ORFs), *ORF1* and *ORF2*, are directly translated from the viral genomic RNA (gRNA) to respectively produce 11 and 15 nonstructural proteins (nsps) through protein cleavage (D. Kim et al. 2020). Then, RNA-dependent RNA polymerase (RdRP), mediated by the translated *nsp12*, uses the gRNA as a template for virus replication and transcription (D. Kim et al. 2020). During this process, the negative-sense RNA intermediates are generated; these then serve as templates for synthesis of both gRNA and subgenomic RNAs (sgRNAs), including those encoding spike protein (*S*), envelope protein (*E*), membrane protein (*M*), and nucleocapsid protein (*N*), as well as a few accessory proteins through a mechanism called discontinuous translation (D. Kim et al. 2020). The gRNA and structural proteins are then packaged in Golgi apparatus to form virion progeny and be released through exocytosis to induce further infection (D. Kim et al. 2020). Because of their characteristic discontinuous translation, all sgRNAs share the same leading sequence as well as the 3’UTR sequence, which are also shared with the initial and replicated genomic RNAs (D. Kim et al. 2020). On the other hand, the 3’ single-cell RNA capture protocol used in (Emanuel et al. 2020) always captures the 3’UTR sequence as well as the other regions enriched with poly-A/T motifs.

Therefore, we hypothesize that the 3’UTR, which indicates the total viral load, can be used as a surrogate of viral RNA progeny to obtain an analog of RNA velocity for SARS-CoV-2 viral genes.

Once the precursor for each SARS-CoV-2 gene is defined, a composite vector field that includes both SARS-CoV-2 genes and host genes can be reconstructed by considering both virus and host genes as a whole. Similarly, all differential geometry qualities (e.g. Jacobian, acceleration, curvature, divergence and curl) can be calculated for both virus and host genes. In particular, we can identify the top regulators or effectors from the host for each SARS-CoV-2 gene to produce host–virus gene interaction maps.

## CODE AVAILABILITY

***Dynamo*** (version: 1.0) is implemented as a Python package and is available through GitHub (https://github.com/aristoteleo/dynamo-release). Notebooks, tutorials for reproducing all figures in this study, and tutorials of ***dynamo*** usage cases are also available through GitHub (https://github.com/aristoteleo/dynamo-notebooks, https://github.com/aristoteleo/dynamo-tutorials).

## DATA AVAILABILITY

The following public cscRNA-seq datasets are used in this study: the bone marrow (Petukhov et al. 2018), chromaffin (Furlan et al. 2017), dentate gyrus (Hochgerner et al. 2018), and fetal forebrain (La Manno et al. 2018) datasets; the scSLAM-seq study’s 10x dataset (Erhard et al. 2019), the pancreatic endogenesis dataset (Bastidas-Ponce et al. 2019) and the hematopoiesis clone tracing dataset (Weinreb et al. 2020). The following public tscRNA-seq datasets are used in this study: scSLAM-seq (Erhard et al. 2019), scNT-seq (Q. Qiu et al. 2020), sci-fate (Cao, Zhou, et al. 2020), and scEU-seq (Battich et al. 2020). All datasets can be directly downloaded with ***dynamo***. The raw and processed data for the 10x scRNA-seq and the scSLAM-seq clone tracing experiment will be accessible via GEO upon publication of this study.

