## Supplementary method for "Mapping Transcriptomic Vector Fields of Single Cells"

### THEORY Supplementary Methods

*[1]Whitehead Institute for Biomedical Research Cambridge, MA, USA. [2]Howard Hughes Medical Institute, Massachusetts Institute of Technology, Cambridge, MA, USA. [3]Department of Computational and Systems Biology, University of Pittsburgh, Pittsburgh, PA, USA. [4]Joint CMU-Pitt Ph.D. Program in Computational Biology, University of Pittsburgh, Pittsburgh, PA, USA. [5]Department of Molecular and Cell Biology, University of California, Berkeley, CA, USA. [6]Department of Mathematics, University of Texas at Arlington, Arlington, TX, USA. [7]Lycia Therapeutics, South San Francisco, San Francisco, CA, USA. [8]California Institute for Quantitative Biosciences, University of California, Berkeley, CA, USA. [9]Microsoft, Redmond, WA, USA. [10]Halıcıoğlu Data Science Institute, University of California San Diego, San Diego, CA, USA. [11]Medical Scientist Training Program, University of California, San Francisco, CA, USA. [12]Genentech Inc., South San Francisco, CA, USA. [13]UPMC-Hillman Cancer Center, University of Pittsburgh, Pittsburgh, PA, USA. [14]Department of Physics and Astronomy, University of Pittsburgh, Pittsburgh, PA, USA. [$]Equal contribution*

**Single-cell RNA-seq, together with RNA velocity and metabolic labeling, reveals cellular states and transitions at unprecedented resolution. Fully exploiting these data, however, requires dynamical models capable of predicting cell fate and unveiling the governing regulatory mechanisms. Here, we introduce *dynamo*, an analytical framework that reconciles intrinsic splicing and labeling kinetics to estimate absolute RNA velocities, reconstructs continuous velocity vector fields that predict future cell fates, and finally employs differential geometry analyses to elucidate the underlying regulatory networks. We applied *dynamo* to a wide range of disparate biological processes including prediction of future states of differentiating hematopoietic stem cell lineages, deconvolution of glucocorticoid responses from orthogonal cell-cycle progression, characterization of regulatory networks driving zebrafish pigmentation, and identification of possible routes of resistance to SARS-CoV-2 infection. Our work thus represents an important step in going from qualitative, metaphorical conceptualizations of differentiation, as exemplified by Waddington's epigenetic landscape, to quantitative and predictive theories.**

*Corresponding authors*:

*X.Q.* ( - @Xiaojie_Qiu - https://www.devo-evo.com/)

*J.X.* ( - @jhxing001 - https://www.csb.pitt.edu/Faculty/xing/)

*J.S.W* ( - @JswLab - https://biology.mit.edu/profile/jonathan-weissman/)

**KEYWORDS**: dynamo, RNA metabolic labeling, vector field reconstruction, differential geometry analysis, RNA Jacobian, acceleration, curvature, divergence and curl, cell fate transitions, whole-cell kinetic models, mathematical modeling, dynamical systems theory, systems biology

INTRODUCTION

### Hidden variables of single-cell transcriptomic datasets affect cell state and dynamics quantification, and vector field reconstruction

There are two fundamental assumptions in reconstructing the vector field from single-cell genomics data: first, the transcriptome is complete (or sufficient) to specify a cell state; and second, the dynamics can be described by a set of memoryless equations, i.e., the temporal propagation of the system depends only on the present state, but not those at prior times. Here, we provide some justifications for those assumptions and discuss the limitations of the vector field reconstruction.

Generically, one can represent the internal state of a cell by the expression levels (and even spatial distributions) of intracellular molecular species, e.g., spliced or unspliced RNAs. Mathematically, one represents the cell state as a vector $\boldsymbol{z} = \{\boldsymbol{x}, \boldsymbol{y}\}$, where $\boldsymbol{x}$ represents the measured spliced and unspliced transcripts (and/or labelled and total RNA in the case of labeling-based scRNA-seq experiments), and $\boldsymbol{y}$ represents all other unmeasured species such as the proteome and epigenome. Let us assume that one can describe the dynamics of a cell by a set of stochastic differential equations (or other forms such as discrete dynamics, for which the following discussions still hold),

$$\frac{\mathrm{d}\boldsymbol{x}}{\mathrm{d}t} = \boldsymbol{F}(\boldsymbol{x}, \boldsymbol{y}, \mu(t)) + \zeta_{\boldsymbol{x}}(\boldsymbol{x}, \boldsymbol{y}, t) \ ,$$
$$\frac{\mathrm{d}\boldsymbol{y}}{\mathrm{d}t} = \boldsymbol{G}(\boldsymbol{x}, \boldsymbol{y}, \mu(t)) + \zeta_{\boldsymbol{y}}(\boldsymbol{x}, \boldsymbol{y}, t) \ .$$

The functions $\boldsymbol{F}$ and $\boldsymbol{G}$ form a vector field in the full space that describes interactions among intracellular species, influence from extracellular environmental factors ($\mu$) including external stimuli and extracellular secretome, and direct interactions with neighboring cells. Biologically, we expect that different layers of gene regulation, e.g., the proteome and transcriptome, are coupled. The extracellular factors $\mu$ are in general explicitly time-dependent. The terms $\zeta_{\boldsymbol{x}}$ and $\zeta_{\boldsymbol{y}}$ refer to random noise, and we assume them to be white noise with zero means. Under this formulation, there are two theoretical issues that must be considered when reconstructing the vector field from single-cell transcriptome data alone. First, in a typical scRNA-seq experiment, only $\boldsymbol{x}$ is measured, and the other variables are hidden. Second, a cell is generally subject to a time-varying extracellular environment.

For simplicity, we restrict ourselves to the case that external stimuli are constant and spatially uniform, whereas in a more general situation the vector field is time-dependent. We also treat direct and indirect cell–cell interactions in a mean-field sense instead of treating the many-body cell–cell interaction problem explicitly. With single-cell multi-modality co-assays that are also augmented with spatial and temporal resolution [1], our framework will allow us to explicitly account for "hidden variables".

If the system dynamics are deterministic, i.e., $\zeta_{\boldsymbol{x}} = \zeta_{\boldsymbol{y}} = 0$, cells evolve along a manifold embedded in the state space of $(\boldsymbol{x}, \boldsymbol{y})$. If one wants to define the metaphorical Waddington's epigenetic landscape, it should be defined on this manifold. In the case that $\boldsymbol{x}$ and $\boldsymbol{y}$ are tightly coupled, and the manifold can be parameterized solely by $\boldsymbol{x}$ , i.e., $\boldsymbol{y} = \boldsymbol{H}(\boldsymbol{x})$, then a cell state can be well represented by the transcriptome alone (but see discussions on **Fig. SI4A** below). Mathematically, this means that we assume that the manifold in the full space and its projection to the $\boldsymbol{x}$ space are homeomorphic.

The presence of stochasticity loosens the coupling. Mathematically, a transcriptome-based quantity $O(\boldsymbol{x})$, e.g., $\mathrm{d}\boldsymbol{x}/\mathrm{d}t$, should be understood as being projected to the subspace of $\boldsymbol{x}$ , i. e., averaged over the hidden variables,

$$\overline{O(\boldsymbol{x})} = \int \mathrm{d}\boldsymbol{x}'\mathrm{d}\boldsymbol{y}' O(\boldsymbol{x}', \boldsymbol{y}')\rho(\boldsymbol{x}', \boldsymbol{y}', t)\delta(\boldsymbol{x}' - \boldsymbol{x}) \ ,$$

where $\rho(\boldsymbol{x}, \boldsymbol{y}, t)$ is the probability density function in the overall state space, and $\delta(x)$ is Dirac's delta function. In the case of time scale separation between transcription and other slower processes (translation, epigenetic modification, etc.), one may further assume that $\boldsymbol{x}$ reaches quasi-steady-state for a given set of $\boldsymbol{y}$, and one can expect that $\rho(\boldsymbol{x}, \boldsymbol{y}, t) \approx \rho_1(\boldsymbol{x}_{ss}(\boldsymbol{y}))\rho_2(\boldsymbol{y}, t)$ also varies slowly in time.

In practice, the above average is typically performed by averaging cells neighboring in the state space, weighted with a specific kernel function [2, 3, 4],

$$\overline{O(\boldsymbol{x})} = \sum_i O(\boldsymbol{x}_i', \boldsymbol{y}_i')K(|\boldsymbol{x}_i' - \boldsymbol{x}|) \ ,$$

with the data set sampled from $\rho(\boldsymbol{x}, \boldsymbol{y}, t)$, and $\sum_i K(|\boldsymbol{x}_i' - \boldsymbol{x}|) = 1$. Notice that $K(|\boldsymbol{x}' - \boldsymbol{x}|)$ is typically chosen as a fast-decaying function to $|\boldsymbol{x}' - \boldsymbol{x}|$, compared to $\rho(\boldsymbol{x}, \boldsymbol{y}, t)$, as well as $\boldsymbol{F}$ and $\boldsymbol{G}$. Due to coupling between $\boldsymbol{x}$ and $\boldsymbol{y}$, one expects that for a given $\boldsymbol{x}$, the data set $O(\boldsymbol{x}_i', \boldsymbol{y}_i')$ that contributes to the average also varies slowly along the hidden variable $\boldsymbol{y}$. Consequently, the average is insensitive to the exact form of $\rho(\boldsymbol{x}, \boldsymbol{y}, t)$ and thus to the experimental sampling, provided that the sampling size is sufficient for statistical convergence. That is, one can understand the reconstructed vector field physically as the bare vector field with local average along the manifold in the full state space which is then projected to the reduced transcriptome space when it is learned in the reduced PCA or UMAP space, or mathematically,

$$\begin{aligned}
\overline{O(\boldsymbol{x})} =& O(\boldsymbol{x}, \bar{\boldsymbol{y}}) + \sum_i \Big[O(\boldsymbol{x}_i', \boldsymbol{y}_i') - O(\boldsymbol{x}, \bar{\boldsymbol{y}})\Big] K\Big(|\boldsymbol{x}_i' - \boldsymbol{x}|\Big) \\
=& O(\boldsymbol{x}, \bar{\boldsymbol{y}}) + \sum_i \left(\frac{\partial O(\boldsymbol{x}, \bar{\boldsymbol{y}})}{\partial \boldsymbol{x}} \cdot (\boldsymbol{x}_i' - \boldsymbol{x})K(|\boldsymbol{x}_i' - \boldsymbol{x}|) + \mathcal{O}(|\boldsymbol{x}_i' - \boldsymbol{x}|^2)\right) \\
&+ \sum_i \left(\frac{\partial O(\boldsymbol{x}, \bar{\boldsymbol{y}})}{\partial \boldsymbol{y}} \cdot (\boldsymbol{y} - \bar{\boldsymbol{y}})K(|\boldsymbol{x}_i' - \boldsymbol{x}|) + \mathcal{O}(|\boldsymbol{y}_i' - \boldsymbol{y}|^2)\right) \ ,
\end{aligned}$$

where $\bar{\boldsymbol{y}}$ is the mean of the contributing data set points, and the data points are expected to have a slowly varying distribution of $\boldsymbol{y}$ peaked around $\bar{\boldsymbol{y}}$.

**Fig. SI4A** schematically shows three typical situations. At points $\boldsymbol{x}_a$ and $\boldsymbol{x}_b$ we assume that the data points within the averaging box have single-peaked distributions along $\boldsymbol{x}$ , as well as the unmeasured $\boldsymbol{y}$. There are situations (e.g., at $\boldsymbol{x}_c$) that the assumption is not valid. The existence of some hidden variables may manifest as a broad and even multi-peaked distribution for the value of the measured quantity (e.g., $\mathrm{d}\boldsymbol{x}_i'/\mathrm{d}t$) in the contributing data set $O(\boldsymbol{x}_i', \boldsymbol{y}_i')$, which may render neighbor cells with similar transcriptomic states with vastly different velocity vectors (See **below**).

#### Caveats on vector field reconstruction

Note that the vector field, defined as $\dot{\boldsymbol{x}}(t) = \boldsymbol{f}\big(\boldsymbol{x}(t)\big)$, does not allow two trajectories to cross each other. Therefore, the input velocity vectors for vector field reconstruction should not have many cells with very similar gene expression states but inconsistent velocity vectors. This can happen either when the data have strong hidden variable effects (case $c$ in **Fig. SI4A**), or when there are potential strong batch effects between different batches of datasets. We expect that the hidden variable issue can be alleviated by single cell multi-omics to capture a more holistic view of cell states, improvements in RNA capture rate, and a reduction in sequencing cost. Further efforts by our group or others will be needed to address the second issue so that we can correct batch effects while performing RNA velocity and vector field reconstruction.

Note that our vector field reconstruction is applicable to both the cscRNA-seq and the tscRNA-seq data. Because the RNA velocity from cscRNA-seq data is relative and scaled by the splicing rate constant $\beta$ for each gene, we explore whether the velocity directionality would be affected by this scaling with relative RNA velocity, especially in the UMAP space. Randomly scaling the velocity vector by a positive value on a few cscRNA-seq data, sampled from the uniform distribution (0, 10) for each gene, however, does not change the velocity vector directionality in UMAP (data not shown), indicating that the sign of velocity is the most important information for revealing the directionality of RNA velocity, especially when projected to a lower dimension. This result may explain why the conventional RNA velocity method, although relative, still proves useful in revealing the directionality of cell fate transitions. When the RNA velocity estimates are relative, the resultant vector field and differential geometry quantities are also relative. In this study, we specifically choose the zebrafish cscRNA-seq dataset to demonstrate the differential geometry analyses, as the dataset has high quality while the associated system involves a complicated differentiation process with many cell types, making it suitable for deep investigation. In future work, it would be interesting to formally test whether the absolute RNA velocity can be used to improve the differential geometry analysis when applied to an interesting biological system with a carefully designed tscRNA-seq experiment.

### Impacts of dimensions of rate constants on RNA velocity

The rate law connects the rate of a reaction and concentrations of involved (bio)chemical species. As an example, the rate law for a first-order reaction that generates a product $A$ is:

$$v = k[A] \ ,$$

where $v$ is the reaction rate, $k$ the first-order rate constant, and $[A]$ is the concentration of product $A$. Because the time scale of a first-order reaction is often characterized by the reciprocal of the rate constant (also known as the "time constant", or "half-life" up to a factor of $\ln 2$ for first-order degradation, i.e., $t_{1/2} = \ln 2/k$), "rate" and "rate constant" are often used interchangeably in certain contexts [2, 3]. They are, however, quantities with different dimensions, and this often leads to confusion, especially for RNA velocity methods. For scRNA-seq data, we assume a constant cell volume (see **below** for more discussions on impacts of the cell volume and others), and the concentration, whose dimension is usually the quantity (or the copies of RNA species) of gene $A$ per unit volume, is replaced by the copy number of $A$. Therefore, the dimension of the reaction rate $v$ is copy number of molecules per unit time (denoted as $N/T$), and that of the first-order rate constant is one per unit time ($1/T$). For a zeroth-order reaction, the rate constant is also the reaction rate, and therefore they share the same dimension.

In the context of RNA velocity, the velocities of unspliced and spliced RNA for a gene are essentially net reaction rates for the production and depletion of unspliced and spliced RNA, with the dimension of $N/T$. Because RNA splicing and degradation are first-order reactions, $\beta$ and $\gamma$ are first-order rate constants with dimension $1/T$. In the original RNA velocity method [2], the degradation rate constant $\gamma$ is scaled by the splicing rate constant $\beta$, so the relative rate constant $\hat{\gamma}$ is dimensionless. The resulting RNA velocity $v = u - \hat{\gamma} s$ does not have the dimension of reaction rates $N/T$, but rather only the number of molecules ($N$), and thus the "velocity" is relative to the splicing rate constant $\beta$. Consequently, suppose that one obtains a small relative RNA velocity for a gene, the actual change in the copy number of spliced RNA per unit time can be large if the splicing rate is fast.

The transcription of unspliced RNA is assumed to be a zeroth-order reaction, so $\alpha$ is a zeroth-order rate constant with the dimension $N/T$. Note that RNA transcription is not an elementary reaction in which products are formed in a single step, but instead a complex reaction with multiple steps involving various trans- and cis-elements. The zeroth-order rate constant $\alpha$ is thus an apparent rate constant under a reduced reaction scheme that lumps many intermediate steps, which are in fact regulated by a variety of internal and external signals. As a result, the transcription rate constant $\alpha$ is a function of cell state in the gene expression space. This has also been shown to be the case for splicing and degradation rate constants [5] although it is reasonable to assume those are constants as we and others did [2, 3].

Here, we would like to provide some thoughts on cell volume. Because typical scRNA-seq data contain no cell volume information, as a zeroth-order approximation we assume a constant cell volume for all cells. This approximation does not affect the sign of estimated RNA velocity because all RNA species in one cell are affected equally. With cell volume information available together with the expression state (e.g., from imaging based methods), it is straightforward to incorporate cell-specific volume information in our parameter estimation procedure.

We also want to comment that in practice, we additionally assume all cells share the same total RNA content, this is, in our preprocessing steps, we scale the total UMI counts in each cell to be 1e4 molecules, similar to many other scRNA-seq analysis toolkits. The normalized gene expression in each cell can be regarded as the fraction of total RNA content occupied by each gene. This normalization scheme is believed to help remove library size differences happen during library construction and sequencing [6].

### Binomial mixture model to quantify labeled and unlabeled RNA

We use the binomial mixture model first described in the GRAM-SLAM study [7] to estimate the fraction of labeled ($\pi_g$) and unlabeled reads of each gene $g$ for the scSLAM-seq data that we produced. The probability of $y$ T-to-C mutations in a read that contains $n$ possible mutation sites can be defined with the following equation:

$$P\left(y; p_{\mathrm{e}}, p_{\mathrm{c}}, n, \pi_{\mathrm{g}}\right) = \left(1-\pi_{\mathrm{g}}\right) B\left(y, n, p_{\mathrm{e}}\right) + \pi_{\mathrm{g}} B\left(y, n, p_{\mathrm{c}}\right) \ ,$$

where $p_e$ is the background T-to-C mutation rate that is independent of the mutations introduced by metabolic labeling , and $p_c$ is the T-to-C mutation rate introduced by metabolic labeling. $B(y, n, p)$ is the binomial probability mass function. Estimation of $p_c$, $p_e$, and $\pi_g$ is performed using the pipeline from [8] with a few custom adaptations. We define the ratio between the true ($\pi_g^{\mathrm{True}}$) and estimated ($\pi_g$) fraction of labeled reads as the labeling correction coefficient, denoted as $\lambda = \pi_g/\pi_g^{\mathrm{True}}$. When the fraction of labeled RNA is overestimated, $\lambda$ is larger than 1 and vice versa.

### Attaining tscRNA-seq dataset used in this study

The scSLAM-seq study [9] used the same binomial mixture model to estimate the labeled and unlabeled RNA. We retrieved the scSLAM-seq data deposited on zenodo (doi: 10.5281/zenodo.1299119). The data deposited by the scEU-seq [5] study provided four species, namely unspliced unlabeled, unspliced labeled, spliced unlabeled and spliced labeled RNA ($u_u$, $u_l$, $s_u$, $s_l$), and were retrieved via the GEO access ID GSE128365. Because scEU-seq manually separated labeled and unlabeled RNA, there is no need for a statistical estimation. However, the manual separation of labeled and unlabeled RNA may introduce potential cross-contamination, and the preparation of two libraries may lead to batch effects. Correction of those possible cross-contamination and batch effects represents an interesting future direction. Data for the sci-fate [10] and scNT-seq [11] studies were obtained through direct communication with the authors before their publication. Custom statistical corrections, as reported in the original studies, were applied to the obtained datasets. Datasets for those studies can now also be downloaded via GEO access IDs GSE131351 and GSE141851, respectively.

### Effects of under and overestimation of labeled RNA fraction on tscRNA-seq kinetic parameter estimation and velocity calculation

For one-shot experiments, the slope of the linear relationship between labeled RNA $l$ and total RNA $r$ is proportional to the labeling correction coefficient:

$$k = \lambda(1 - e^{-\gamma t}) \ .$$

Therefore, an overestimated labeling correction coefficient amounts to a high NTR (new to total RNA ratio) at steady state. For one-shot experiment, we assume that the labeling data have been statistically well corrected with the mixture binomial model, and thus the labeling correction coefficient is effectively close to 1. Then the slope is approximately $k = 1 - e^{-\gamma t}$, allowing us to obtain the degradation rate constant $\gamma$ from the NTR slope. We can evaluate the error between the estimated $\gamma$ under this assumption, and the true degradation rate constant $\gamma^{\text{true}}$:

$$\begin{aligned}\gamma - \gamma^{\text{true}} &= -\frac{1}{t}\Big(\ln(1 - \frac{k}{\lambda}) - \ln(1 - k)\Big) \\ &= -\frac{1}{t}\ln\Big(\frac{1 - \frac{k}{\lambda}}{1 - k}\Big) \ .\end{aligned}$$

When $\lambda < 1$, $\gamma < \gamma^{\text{true}}$, $\gamma$ is underestimated. Consequently, the velocity of total RNA, considering only its magnitude, differs from the true velocity by,

$$\begin{aligned}|\dot{r}| - |\dot{r}^{\text{true}}| &= \Big|\frac{\gamma n}{k} - \gamma r\Big| - \Big|\frac{\gamma^{\text{true}} n}{k} - \gamma^{\text{true}} r\Big| \\ &= (\gamma - \gamma^{\text{true}})\Big|\frac{n}{k} - r\Big| \ .\end{aligned}$$

Note that $n/k - r$ determines the sign of the velocity, i.e., $n/k - r > 0$ amounts to a positive velocity, and vice versa. Therefore, under-correction of labeled RNA leads to underestimation of the velocity. It is also apparent that a labeling correction coefficient higher than one leads to overestimation of both the degradation rate constant $\gamma$ and the velocity of total RNA. Labeling correction coefficient, which is assumed to be constant across all time points, has minimal impacts on curve-fitting methods for degradation rate constants because the time scale of a first order degradation is independent of initial concentrations. However for kinetics experiments, the labeling efficiency affects the curve-fitting of both the synthesis of labeled RNAs, and the degradation of unlabeled RNAs. The transcription rate $\alpha$ is under-estimated when $\lambda < 1$, and overestimated when $\lambda > 1$. The kinetics of unlabeled RNA are not merely a degradation process when $\lambda < 1$, as there are artificial increases of new RNA due to underestimation of labeled RNA, and the degradation appears slower than the true rate. By contrast, when $\lambda > 1$, the degradation of the unlabeled RNA is unaffected for similar reasons as in degradation experiments. As a result, the cluster-wise velocity for kinetic experiments is underestimated when $\lambda < 1$ and overestimated when $\lambda > 1$. In the extreme cases, an underestimated labeling RNA fraction can lead to a sign change in the velocity. On the other hand, because the cell-wise velocity is:

$$\dot{r} = \frac{\gamma l}{1 - e^{-\gamma t}} - \gamma r$$

It is unaffected by the inaccurate estimation of $\alpha$, and an inaccurately estimated $\gamma$ alters its magnitude but not the sign.

### Estimation of parameter ranges for curve-fitting methods

To overcome the local optima of the cost function and speed up parameter estimation, we need to have good guesses of parameters and the valid ranges of those parameters. A set of parameter ranges are used for initial parameter value sampling and providing upper and lower bounds for optimizers to avoid unrealistic results. The "guesstimated" values $\theta_0$ for specific parameters are first determined, according to the specific labeling strategy used. The range of the parameters is then simply set to be $(0, 100\ \theta_0)$. The methods for obtaining "guesstimations" are different for each parameter:

1) Kinetics experiments

If the RNA dynamics are far from steady state and degradation is negligible, then the amount of newly synthesized RNA is proportional to the labeling time:

$$l_t \sim \alpha t\ ,$$

where $l_t = n(t)$, i.e. the number of copies of new RNA at labeling time $t$. Thus, the guesstimated $\alpha$ is simply the averaged ratio of new RNA and labeling time. The degradation rate constant can be roughly estimated from the old RNA:

$$\gamma \sim \frac{1}{t} \ln \frac{o_0}{o_t}\ .$$

The splicing rate constant is estimated in a similar manner:

$$\beta \sim \frac{1}{t} \ln \frac{u_u(0)}{u_u(t)}\ .$$

2) Degradation experiments

The guesstimated values for the initial conditions, including $l_0$, $u_{l,0}$, and $s_{l,0}$, are simply the average abundance of labeled RNAs across all cells belong to the initial labeling time point. The degradation rate constant is guesstimated with the labeled RNA, using a equation similar to the one for kinetics experiments:

$$\gamma \sim \frac{1}{t} \ln \frac{l_t}{l_0} \ .$$

The splicing rate constant is estimated with:

$$\beta \sim \frac{1}{t} \ln \frac{u_l(t)}{u_l(0)} \ .$$

#### Goodness of fit for linear regression and curve-fitting methods during kinetic parameters estimation

For linear regression models, given the data and model predictions $\{x_i, y_i\}_{i=1}^{n}$ for $n$ cells, the goodness of fit is determined using the standard R-squared:

$$R^2 = 1 - \frac{\sum_{i=1}^{n}(x_i - y_i)^2}{\sum_{i=1}^{n}(x_i - \bar{x})^2} \ ,$$

where $\bar{x}$ is the mean of data. For curve-fitting methods, the Gaussian log-likelihood is used as a measure for goodness of fit. Given the data and model predictions $\{\boldsymbol{x}_i, \boldsymbol{y}_i\}_{i=1}^{k}$ of $k$ species, where each $\boldsymbol{x}_i$ and $\boldsymbol{y}_i$ is a vector of model predictions for $m$ time points, the Gaussian log-likelihood is:

$$\ln G(\boldsymbol{x}_1, \boldsymbol{x}_2, \ldots, \boldsymbol{x}_k | \boldsymbol{y}_1, \boldsymbol{y}_2, \ldots, \boldsymbol{y}_k) = -\frac{n}{2} \ln(2\pi) - \sum_{i=1}^{k} \ln\big(\sigma(\boldsymbol{x}_i)\big) - \sum_{i=1}^{k} \frac{1}{2} \|\bar{\boldsymbol{x}}_i - \boldsymbol{y}_i\|^2$$

where $\bar{\boldsymbol{x}}$ and $\sigma(\boldsymbol{x})$ are the mean and standard deviation of $\boldsymbol{x}$, respectively. To balance the numerical difference between species, the data and model predictions can be normalized by the maximal value of data for each species.

#### Outlier detection in vector field reconstruction

Outlier detection is vital for robust vector field function reconstruction from noisy RNA velocity data. The sparseVFC algorithm [12] models noise in velocities $\boldsymbol{v}$ of inliers with a Gaussian distribution, i.e.:

$$P(\boldsymbol{v} | z = 1, \boldsymbol{x}, \boldsymbol{\theta}) = \frac{1}{(2\pi\sigma^2)^{d/2}} e^{-\frac{\|\boldsymbol{v} - \boldsymbol{f}(\boldsymbol{x})\|^2}{2\sigma^2}} \ ,$$

where $z$ is an indicator variable, such that $z = 1$ when the cell is an inlier, and $z = 0$ otherwise. $\boldsymbol{\theta}$ contains all parameters, including the variance of the Gaussian distribution $\sigma^2$, the vector field function $\boldsymbol{f}$, and the prior probability $q$ mentioned below. The probability distribution of outliers is modeled with a uniform distribution:

$$P(\boldsymbol{v} | z = 0, \boldsymbol{x}, \boldsymbol{\theta}) = \frac{1}{a} \ ,$$

where $a$ is the volume of the domain for velocity vectors. Empirically, this is a parameter used for adjusting the aggressiveness of the outlier detection. Denote the fraction of inliers as $q$:

$$q = P(z = 1|\boldsymbol{x}, \boldsymbol{\theta}) \ .$$

Then, this is essentially a mixture model where the likelihood is:

$$\begin{aligned} P(\boldsymbol{v}|\boldsymbol{x}, \boldsymbol{\theta}) &= qP(\boldsymbol{v}|z = 1, \boldsymbol{x}, \boldsymbol{\theta}) + (1-q)P(\boldsymbol{v}|z = 0, \boldsymbol{x}, \boldsymbol{\theta}) \\ &= \frac{q}{(2\pi\sigma^2)^{d/2}} e^{-\frac{\|\boldsymbol{v}-\boldsymbol{f}(\boldsymbol{x})\|^2}{2\sigma^2}} + \frac{1-q}{a} \ , \end{aligned}$$

and the posterior probability can be derived from Bayes' theorem (notice that the following corrects an error in [12]):

$$\begin{aligned} P(z = 1|\boldsymbol{v}, \boldsymbol{x}, \boldsymbol{\theta}) &= \frac{qP(\boldsymbol{v}|z = 1, \boldsymbol{x}, \boldsymbol{\theta})}{P(\boldsymbol{v}|\boldsymbol{x}, \boldsymbol{\theta})} \\ &= \frac{e^{-\frac{\|\boldsymbol{v}-\boldsymbol{f}(\boldsymbol{x})\|^2}{2\sigma^2}}}{e^{-\frac{\|\boldsymbol{v}-\boldsymbol{f}(\boldsymbol{x})\|^2}{2\sigma^2}} + \frac{1-q}{q}\frac{(2\pi\sigma^2)^{d/2}}{a}} \end{aligned}$$

For $n$ such independent and identically distributed (i.i.d.) RNA velocity data samples, one can construct a diagonal matrix $\boldsymbol{P} = \mathrm{diag}(p_1, p_2, \ldots, p_n)$, where $p_i = P(z_i = 1|\boldsymbol{v}_i, \boldsymbol{x}_i, \theta)$. The E-step of the EM algorithm evaluates this matrix, which is used as the weight in the loss function for sparseVFC (see **MATERIAL AND METHODS**).

To update $\sigma$ and $q$ at the M-step of each EM iteration following standard EM algorithm procedure, the updated parameters are the solutions of the following optimization problem [12]:

$$\boldsymbol{\theta}^{\mathrm{new}} = \mathrm{argmax}_{\boldsymbol{\theta}} \mathcal{Q}(\boldsymbol{\theta}, \boldsymbol{\theta}^{\mathrm{old}}) \ ,$$

where $\mathcal{Q}(\theta, \theta^{\mathrm{old}})$ is a conditional expectation of the complete-data log-likelihood function:

$$\mathcal{Q}(\boldsymbol{\theta}, \boldsymbol{\theta}^{\mathrm{old}}) = \sum_{\boldsymbol{z}} P(\boldsymbol{z}|\boldsymbol{V}, \boldsymbol{X}, \boldsymbol{\theta}^{\mathrm{old}}) \ln P(\boldsymbol{V}, \boldsymbol{z}|\boldsymbol{X}, \boldsymbol{\theta}) \ ,$$

with

$$\begin{aligned} P(\boldsymbol{z}|\boldsymbol{V}, \boldsymbol{X}, \boldsymbol{\theta}^{\mathrm{old}}) &= \prod_{i=1}^{n} P(z_i|\boldsymbol{v}_i, \boldsymbol{x}_i, \boldsymbol{\theta}^{\mathrm{old}}) \ , \\ P(\boldsymbol{V}, \boldsymbol{z}|\boldsymbol{X}, \boldsymbol{\theta}) &= \prod_{i=1}^{n} P(\boldsymbol{v}_i, z_i|\boldsymbol{x}_i, \boldsymbol{\theta}) \ . \end{aligned}$$

With i.i.d. samples, one can show that [12]:

$$\begin{aligned}
\mathcal{Q}(\boldsymbol{\theta}, \boldsymbol{\theta}^{\text{old}}) &= \sum_{i=1}^{n} \sum_{z_i=0}^{1} P(z_i | \boldsymbol{v}_i, \boldsymbol{x}_i, \boldsymbol{\theta}^{\text{old}}) \ln P(\boldsymbol{v}_i, z_i | \boldsymbol{x}_i, \boldsymbol{\theta}) \\
&= \sum_{i=1}^{n} \sum_{z_i=0}^{1} P(z_i | \boldsymbol{v}_i, \boldsymbol{x}_i, \boldsymbol{\theta}^{\text{old}}) \ln \Big( P(\boldsymbol{v}_i | z_i, \boldsymbol{x}_i, \boldsymbol{\theta}) P(z_i | \boldsymbol{x}_i, \boldsymbol{\theta}) \Big) \\
&= \sum_{i=1}^{n} \left\{ p_i \Big( \ln q - \frac{d}{2} \ln(2\pi\sigma^2) - \frac{\|\boldsymbol{v}_i - \boldsymbol{f}(\boldsymbol{x}_i)\|^2}{2\sigma^2} \Big) + (1 - p_i) \ln \frac{1-q}{a} \right\}
\end{aligned}$$

By taking derivatives of $\mathcal{Q}(\theta, \theta^{\text{old}})$ w.r.t. $\sigma^2$ and $q$ and equating them to zero, one obtains the solutions for updating the parameters:

$$\begin{aligned}
\sigma^2 &= \frac{(\boldsymbol{V} - \boldsymbol{F})^{\mathrm{T}} \boldsymbol{P} (\boldsymbol{V} - \boldsymbol{F})}{d \times \mathrm{tr} \boldsymbol{P}} \ , \\
\gamma &= \mathrm{tr} \boldsymbol{P} / n \ ,
\end{aligned}$$

where $\boldsymbol{F} = [\boldsymbol{f}(\boldsymbol{x}_1), \boldsymbol{f}(\boldsymbol{x}_2), ..., \boldsymbol{f}(\boldsymbol{x}_n)]^{\mathrm{T}}$.

#### Effects of parameters in vector field function reconstruction

The sparseVFC algorithm with an isotropic Gaussian kernel has four main parameters: the number of control points $m$, the regularization parameter $\lambda$, the inverse bandwidth of the Gaussian kernel $\beta$, and the maximal number of iterations $N_{max}$. Their default values and effects of changes in these values on the resultant vector field function are summarized in the following table:

| | Default | Effects |
|---|---|---|
| $m$ | 5% of the number of cells, with a minimum of 50 control points | Too small: the approximation of the vector field in RKHS is too sparse (underfitting); Too large: the optimization of the loss function is memory- and time-consuming. |
| $\lambda$ | 3 | Too small: overfitting; Too large: underfitting. |
| $\beta$ | determined by the distribution of the data (see **below**) | Too small: large bandwidth means all control points have approximately equal contributions to all surrounding states in the vector field, and the vector field function becomes linear; Too large small bandwidth means control points have insufficient influence over distant states, resulting in zero velocities evaluated for distant cells. |
| $N_{max}$ | 500 | Too small: the algorithm is terminated before reasonable convergence (underfitting); Too large: when convergence is hard to achieve, the algorithm takes too long with negligible improvements in minimizing the loss function. |

The inverse bandwidth $\beta$ is determined in the following way:

1) Find k-nearest-neighbors for each cell ($k$ is by default 20% of the number of cells);
2) Compute the mean distance of each cell to its neighbors $d_m$;
3) The inverse bandwidth $\beta = 1.5/(\sqrt{2}d_m)$, so that the standard deviation of the Gaussian kernel is $\sigma = d_m/1.5$.

### Ranking genes based on differential geometrical quantities

Generally, given some quantity (expression, velocity, acceleration, curvature, etc.) calculated for each gene in each cell, i.e. a $n \times m$ matrix $\boldsymbol{Q}$, where $n$ is the number of cells, and $m$ the number of genes, one can obtain a gene-wise vector of such quantities by averaging over cells:

$$q_j = \langle Q_{\cdot,j} \rangle = \frac{1}{n} \sum_{i=1}^{n} Q_{i,j} \ .$$

Suppose that cells are divided into several clusters, e.g., distinct cell types, the above average can be calculated for each cluster:

$$q_j^c = \langle Q_{\cdot,j} \rangle_c = \frac{1}{n_c} \sum_{i \in C} Q_{i,j} \ ,$$

where $C$ is the set of cells in cluster $c$, and $n_c$ the number of cells in $C$. When one is interested in the absolute values of the quantities, the average is calculated with $|\boldsymbol{Q}|$. Then, genes can be ranked based on $q_j^c$ for each cluster. For the ranking of the Jacobian, since each cell is associated with an $m \times m$ Jacobian matrix, the whole data is an $m \times m \times n$ 3D matrix. The same averaging method is applied to all cells or each cluster:

$$\langle J_{\zeta,\xi,\cdot} \rangle = \frac{1}{n} \sum_{i=1}^{n} J_{\zeta,\xi,i} \ ,$$
$$\langle J_{\zeta,\xi,\cdot} \rangle_c = \frac{1}{n_c} \sum_{i \in C} J_{\zeta,\xi,i}$$

Because for each cell or cluster, the Jacobian or average Jacobian is an $m \times m$ matrix, and the ranking can be performed in various ways:

1) Top interactions: because each element in the averaged Jacobian indicates the change in the velocity of the putative effector with respect to the change in the expression of the putative regulator, top elements suggest strong gene–gene interactions in each cell or cell type, as below.

2) Top putative regulators for each effector: we rank each row of the averaged Jacobian matrix, so that for each effector, one obtains the top genes that potentially regulate the effector.

3) Top putative effectors for each regulator: we rank each column of the averaged Jacobian matrix, so that for each regulator, one obtains top genes potentially regulated by the regulator.

4) Top putative regulators: For effector $\zeta$, we sum up its averaged Jacobian elements with respect to all possible regulators:

$$R_\zeta = \sum_{\xi=1}^{n} \langle J_{\zeta,\xi,\cdot} \rangle = \sum_{\xi=1}^{n} \left\langle \frac{\partial f_\zeta}{\partial x_\xi} \right\rangle \ ,$$

and rank all $R_\zeta$, which shows the top genes potentially involved in the regulation of others;

5) Top putative effectors: For regulator $\xi$, a summation is taken across all effectors:

$$E_\xi = \sum_{\zeta=1}^{n} \langle J_{\zeta,\xi,\cdot} \rangle = \sum_{\zeta=1}^{n} \left\langle \frac{\partial f_\zeta}{\partial x_\xi} \right\rangle \ .$$

The ranking of all $E_\xi$ reveals putative top regulated genes.

### Analysis details for the scNT-seq dataset

The wild-type and *Tet1/2/3* triple-knockout (TetTKO) datasets for studying the bidirectional transition between mESC pluripotent and totipotent state from [11] were used in this study. The wild-type experiment used the degradation metabolic labeling scheme, whereas the TetTKO experiment used the one-shot metabolic labeling scheme. From both experiments, we obtained unspliced, spliced, labeled, and total RNA data for each gene in each cell. To estimate the absolute degradation rates for each gene in the wild-type dataset, we used the labelled and total RNA data and apply a curve-fitting estimation approach that builds on **Model 2** (**Fig. SI2A**), which does not consider splicing, and assumes a first-order decay for the RNA. We used the labeling and splicing kinetics integration method to estimate the absolute splicing rate constants by dividing the absolute degradation rate constants by the corresponding relative degradation rate constants that are estimated with spliced and unspliced RNA data from the same dataset. Absolute splicing and degradation rate constants for each gene were then used for absolute RNA velocity calculation, velocity projection to 2D UMAP space of spliced RNAs, vector field reconstruction (in the top 30 PC space), differential geometry analyses (e.g., Jacobian calculation), etc.

For the TetTKO dataset, we used the labeled and total RNA data to estimate absolute transcription and degradation rate constants with the "one-shot" method, which explicitly considers the time of the RNA metabolic labeling. Absolute transcription and degradation rate constants then gave us absolute total RNA velocity. Note that the transcription rates calculated here were cell- and gene-dependent (i.e., they corresponded to a cell-by-gene matrix like the expression matrix). On the other hand, the spliced and unspliced RNA were used to estimate the relative degradation rate constants. By integrating with the absolute degradation rate constants from the "one-shot" method, we obtained absolute splicing and degradation rate constants, which gave us the absolute spliced RNA velocity. The absolute total RNA velocity or spliced RNA velocity was then projected to the total RNA-based or spliced RNA–based 2D UMAP and used for vector field reconstructions (in the top 30 PC space), differential geometry analyses (e.g. Jacobian calculation), etc.

#### Analysis details for the scEU-seq dataset

Both the kinetics and mixture labeling experiment datasets of the cell cycle study using human RPE-1 cell line from [5] were used in this study. The degradation labeling experiment dataset of the intestinal organoid study from [5] was also used. We retrieved unspliced unlabeled, unspliced labeled, spliced unlabeled, and spliced labeled RNA data $(u_u, u_l, s_u, s_l)$ for each gene in each cell from all experiments which then gave us unspliced $(U)$, spliced $(S)$, labeled $(L)$ and total $(T)$ RNA data (because $U = u_u + u_l$, $S = s_u + s_l$, $L = u_l + s_l$, $T = U + S$). We mainly focused on analyzing the kinetics and degradation labeling experiment, while demonstrating the generalizability of our estimation framework and revealing the high transcription rates for mitochondrial genes with the mixture labeling experiment. For the kinetics experiment, we used the labeled and total RNA data and the "two-step" method to estimate the absolute transcription (cell- and gene-dependent, as above) and degradation rate constants. Next we used the integration method to estimate absolute splicing rate counts with the relative degradation estimated from the unspliced and spliced RNA data. With the absolute transcription, splicing, and degradation rate constants, we can obtain absolute unspliced, spliced, labeled (or new), and total RNA velocities. The absolute total RNA velocity or spliced RNA velocity was then projected to the total RNA–based or spliced RNA–based 2D UMAP, and are used for vector field reconstructions (in the top 30 PC space), differential geometry analyses (e.g., Jacobian calculation), etc. For the mixture experiment, which had a fixed time period that includes a variable initial kinetics experiment and later accompanying degradation experiment [Fig. SI7 from [5]], we used a curve fitting strategy under **Model 2** (**Fig. SI2A**) to estimate the transcription and degradation rate constants. For the degradation experiment, we used the same strategy as mentioned above for the degradation experiment data from scNT-seq.

#### Functional analysis of kinetic rates calculated from scNT-seq or scEU-seq studies

Recent studies showed that degradation is slower for human proteins than their mouse counterparts during both embryonic segmentation [13] and motor neuron differentiation [14]. Because we calculated the degradation and splicing rate constants in the mESCs cells and hRPE-1 cells with data from the scNT-seq [11] and scEU-seq studies [5] respectively, we can compare the degradation and splicing rate constants between human and mouse ortholog genes. The database of human and mouse ortholog genes was retrieved from from ensembl bioMart [15].

We also tested whether genes with high or low splicing and transcription rate constants are enriched for particular biological pathways. For the mESC degradation study, we compared the cumulative distribution of the degradation and splicing rate constants from housekeeping genes and other genes. The database of housekeeping genes was retrieved from `https://www.genomics-online.com/resources/16/5049/housekeeping-genes/`. For the hRPE-1 kinetics study, we took the top 10% of genes with the fastest splicing and degradation rate constants, and then subject them to GO pathway enrichment analysis.

#### Analysis details for the scSLAM-seq dataset

Both the 10x and the scSLAM-seq datasets from [9] were analyzed in this study. We used the stochastic splicing model, equivalent to the stochastic model from [3], to estimate the relative spliced RNA velocity from both the full dataset and the randomly downsampled datasets that have the same number of cells as in the plate-based scSLAM-seq dataset. The "one-shot" method was applied to the labeled (new) and total RNA data from the scSLAM-seq study to estimate absolute total RNA velocity. The stochastic splicing model was used to estimate the relative spliced RNA velocity for the unspliced and spliced RNA data from the scSLAM-seq study. We calculated the average Pearson correlation between each cell's velocity vector and its neighbor cells' velocity vectors and used this as a metric to quantify the consistency of RNA velocity flow as well as the optimality of data types for velocity vector field reconstructions.

#### Analysis details for the sci-fate dataset

The new and total RNA data from [10] were analyzed in this study. The absolute transcription and degradation rate constants, as well as the associated absolute total RNA velocity were estimated with the "one-shot" model. Genes from the original study reported to be associated with cell-cycle and glucocorticoid receptor (GR) response (Supplementary Table 2 of [10]) were used for the separated and combined RNA velocity analyses. To formally test whether the cell-cycle progression is independent of GR response, we first reconstructed the vector field on the 4D PCA space or the 3D UMAP space that was reduced from the combined expression space with cell-cycle and glucocorticoid receptor (GR) response genes, using the corresponding projected cell state and velocity vector pairs. We then calculated the Jacobian between those UMAP or PCA components in each cell. Overall high-magnitude Jacobian values across cells indicate a strong coupling between those processes related to those components and vice versa. The first and second principal components were related to linear GR response, whereas the third and fourth principal components were related to the cell cycle process. The first UMAP space is related to the GR response, whereas the second and third were related to the cell cycle process.

#### Analysis details for the zebrafish dataset

We used *velocyto* [2] (`http://velocyto.org/`) to parse the bam files generated by running the cellranger (`https://github.com/10XGenomics/cellranger`) pipeline on the zebrafish 10x dataset from [16], to retrieve the unspliced and spliced RNA count matrices. The default preprocessing, dimension reduction, relative RNA velocity estimation and projection, and vector field reconstruction implemented in ***dynamo*** were applied. The fixed points were also identified for the 2D UMAP space based vector field, using default parameters in ***dynamo***. Scatterplots were used to visualize marker gene expression kinetics over vector field based pseudotime, estimated with the *ddhodge* algorithm [17]. The RNA speed, divergence, acceleration and curvature were calculated with the PCA-based vector field function. The magnitude (length of the vector) of the acceleration and curvature in each cell were also calculated. Then, scatterplots of cells on UMAP space were colored according to the RNA speed, divergence, acceleration magnitude, and curvature magnitude with either a viridis or blue-white-red (bwr) colormap, together with the streamlines based on the projected RNA velocity on the UMAP space. RNA acceleration on PCA space was then projected back to the original gene expression space and used to obtain the top-ranked acceleration genes (based on magnitude of acceleration values) for each cell type. The top 10% ranked acceleration in the previously unknown cell type was then subjected to GO pathway enrichment analysis. Genes appeared in the enriched chondrocyte lineages were further used for Jacobian analysis to identify putative interaction between *erbb3b* and other chondrocyte marker genes. Those putative interactions were then visualized with the Arcplot. We also identified the top interactions involved in cells belonging to the iriphore lineage and visualized those interactions with a CircosPlot. The interaction between *tfec* and *pnp4a* was also visualized either on the UMAP space or the gene expression space of *tfec* and *pnp4a*, with cells colored according to cell-wise Jacobian values.

### Analysis details for the pancreatic endocrinogenesis dataset

To discover gene–gene interactions related to pancreatic endocrinogenesis [18], we included as many relevant genes as possible in the data set. A total of 4000 genes were selected using the standard feature selection method implemented in ***dynamo***. A 2D vector field based on UMAP embedding from the original study [18] was first reconstructed and next used to identify fixed points. Then, a 30-dimensional (30-D) vector field was reconstructed using the first 30 PCs along with the projected RNA velocity vectors. The divergence and acceleration were calculated with the 30D vector field. The gene-wise Jacobian matrix was calculated for relevant genes (*Pax4, Neurog3, Ins2, Pdx1*, etc.), which shows their interactions in different cell types (**Fig. SI 5G**). Cell type–specific networks were constructed based on the Jacobians averaged across all cells from a cell-type (see the above section) and illustrated using Arcplots (**Fig. SI 5G**).

### Analysis details for other cscRNA-seq datasets used in this study

To demonstrate ***dynamo***'s general applicability and robustness in relative RNA velocity estimation, vector field reconstruction, and the differential geometry analyses, we analyzed a variety of disparate biological systems profiled with cscRNA-seq, including bone marrow [19], chromaffin [20], dentate gyrus [21], and fetal forebrain [2] datasets in addition to our HL60 10x dataset, the scSLAM-seq study's 10x dataset, zebrafish dataset [16] and the pancreatic endogenesis dataset [18]. We used default parameters to preprocess those datasets, followed by performing dimension reduction, estimating and projecting relative RNA velocity to the first two PCA components for the fetal forebrain dataset (as used in the original study) or UMAP space for all other datasets (The same selection of the PCA or UMAP space for each dataset was taken in the following). To demonstrate the generality of relative velocity estimation, phase plots, scatterplots of gene expression, and velocity of marker genes were applied to all datasets. We then reconstructed the velocity vector fields and identified the fixed points on the first two PCA components or UMAP space. Vector field–based pseudotime was also estimated for all datasets, and scatterplots of expression kinetics were used to visualize the corresponding marker gene's expression over pseudotime. RNA speed (length of velocity vector), divergence, and acceleration magnitude (length of acceleration vector) were calculated in the PCA based vector field; curl, which is defined on two- or three-dimensional space, was calculated using the first two PCA components or the UMAP space–based vector field. Finally, embedding scatterplots of cells on the first two PCA components or UMAP space were colored according to the RNA speed, divergence, acceleration magnitude, and curl with either a viridis or blue-white-red (bwr) colormap, together with the streamlines based on the projected RNA velocity on the same space.

### Analysis details of the HL60 cell differentiation datasets

#### Process clone barcode and build "cell linkages"

Based on the conserved sequences flanking the cellular barcodes (GBCs), we retrieved the GBCs sequence for all reads in each cell from the scSLAM-seq clone tracing experiment and formed a cell by barcode matrix in which each element corresponds to the number of reads for that barcode observed in that cell. After removing barcodes with low reads across cells, we calculated the Leivenstein distance between all pairs of the remaining barcodes and applied affinity propagation clustering algorithm to group barcodes into 666 clone clusters and identify a barcode exemplar for each cluster. Because the clustering algorithm itself does not incorporate a hard distance threshold between barcodes belonging to this barcode cluster and the exemplar of this cluster, we used a custom script to iteratively search for barcodes that had a Levenstein distance ¿ 3 from the cluster exemplar or any newly identified exemplars, and appended those as new barcode cluster exemplars in addition to the existing ones. This approach yielded 764 uniquely identified barcodes. On the cell level, most cells had only one barcode, but a few that had two or more. In order to identify only confident cell linkages in which two or more cells shared the same barcode and to avoid spurious linkages, we explicitly ignored cells processed at nearby wells of a 384-well plate that had the same barcode as clone cells (**Fig. 5E**). Because the wells in those plates were extremely small, cross-contamination between nearby wells can occur, leading to spurious cell linkage. This was less an issue for transcriptome qualification because the amount of leakage relative to the entire transcriptome was relatively small.

#### Analyze the 10x and scSLAM-seq datasets

We used default parameters to preprocess the 10x data and the unspliced and spliced RNA data from the scSLAM-seq experiment, and then performed dimension reduction and estimated and projected relative RNA velocity to the UMAP space for both datasets. For the one-shot labeling data from the scSLAM-seq experiment, the "one-shot" method was used with default parameters to estimate the absolute transcription rate and degradation rate constants, which were then integrated with the splicing data to obtain absolute splicing rate constants, as well as absolute spliced and total RNA velocity. Scatterplots of marker gene expression of progenitors and neutrophil lineages, as well as streamline plots with cells colored by sample collection time points on UMAP space across all datasets (10x, splicing data, and labeling data from the scSLAM-seq clone tracing experiment) were used to visualize neutrophil lineage commitment.

### Analysis details for the hematopoiesis dataset

We used hematopoietic datasets from [22], which included three major experiments: an *in vitro* experiment in which HSCs were cultured in competent differentiation media; a cytokine perturbation experiment in which HSCs in different plates received different differentiation factors, such as MPO or EPO; and an *in vivo* experiment in which barcoded HSCs were first allowed to proliferate *in vitro* for 2 days and then transplanted into 10 irradiated host mice whose blood cells were later harvested at week 1 and 2. Both of the first two experiments were subject to clone tracking on days 0, 2, 4, and 6, and all experiments were sequenced via inDrop-seq. Although the sequencing depth was not high (only 600 genes on average), roughly 100,000 cells are sequenced in each experiment. We used kb-python (https://github.com/pachterlab/kb_python) to reprocess the data to obtain unspliced and spliced counts for each cell.

We first performed velocity analysis on those datasets using ***dynamo*** with default parameters; however, for all three datasets, this resulted in unexpected backward velocity flows from terminal cell types to undifferentiated cells, based on cell-type assignments from the original study [22]. After carefully ruling out issues with RNA velocity estimation, we reached the conclusion that the shallow sequencing of this study was the culprit of the backward velocity flow. We noticed that such biologically conflicting results have been observed by others and circulated online. In fact, RNA velocity estimation is prone to be problematic if the intron capture is insufficient or biased, as in the case of shallow sequencing. Hence, we were motivated to develop a heuristic method that uses some prior (of broad cell lineage hierarchy) to filter genes whose expression kinetics does not follow clockwise dynamics on the spliced–unspliced RNA phase plane. This supervised method (see details below) was used to correct the relative RNA velocity estimation and vector field reconstructions for all three datasets (in vitro, cytokine perturbation, and the in vivo experiment).

#### Stable limit cycle detection and redundant trajectory removal for numerical integration of vector field functions

Stable limit cycles cause redundant sampling for trajectories integrated using vector field functions. In this study, we focus on detecting limit cycles for a 2D vector field, but the method can be easily generalized to higher dimensions. Suppose we have a trajectory of in total $n$ points $\{(x_i, y_i)\}_{i=1}^n$, and we divide it into $k$ ($k = 4$ by default) intervals, each of which contains $m$ points $\{\boldsymbol{X}_j, \boldsymbol{Y}_j\} = \{(x_i^j, y_i^j)\}_{i=1}^m$, where $j = 1, 2, \ldots, k$. If a portion of the trajectory enters a stable limit cycle and orbits around the corresponding fixed point, the $x$- and $y$-coordinates of the points are periodic. We use the fast Fourier transform to obtain the frequency spectra for the $x$- and $y$- coordinates of points in the last two intervals:

$$\boldsymbol{f}_x^j = \mathrm{FFT}(\boldsymbol{X}_j) \ ,$$
$$\boldsymbol{f}_y^j = \mathrm{FFT}(\boldsymbol{Y}_j) \ ,$$

where $j = k-1, k$. Let $\boldsymbol{f}^j = [\boldsymbol{f}_x^j, \boldsymbol{f}_y^j]$ be the concatenated frequency spectrum. If the spectral difference $\frac{1}{m}\|\boldsymbol{f}^{k-1} - \boldsymbol{f}^k\|$ is smaller than a certain threshold (0.05 by default), the last interval is considered redundant and thus removed. This procedure is performed iteratively, until the redundancy criterion is not met.

#### Prediction of cell fate via integration of vector field given initial cell states, and fate probability estimation

Once the vector field function was learned, either in reduced UMAP space, top PCA space or even the original gene expression space, we can use it to predict the historical and future cell expression states over arbitrary time scales given any initial cell state. This can be achieved by integrating the continuous vector field function from one or a set of initial cell states forward or backward in time. When the integration was performed for the early cells of a particular clone, the integration paths can be used to calculate the minimal distances from clone cells at later time points to the paths, as well as the fate bias (see **below**), to validate the accuracy and single cell trajectory predictivity of the reconstructed vector field function, as demonstrated in the HL60 or the hematopoiesis clone tracing datasets analyses (**Fig. 5**).

Fate probability is currently calculated as the percentage of points along the predicted cell fate trajectory whose first-nearest neighbors of the observed cells belong to any particular cell group, e.g., cell type. The distances to the first-nearest neighbors are required to be small, and are determined by median diffused first-nearest neighbors of all observed cells and a distance threshold, see details below:

Cell fate trajectories predicted by our vector field method sometimes end up in regions where few or no cells were actually measured (see the above section). A heuristic method is thus used to iteratively walk backward the integration path to assign cell fate. We first identify the regions with small velocity in the tail of the integration path, which is determined by a threshold of speed, and then check whether the distance of first-nearest points on the observed data (cells) to all those points are far from the observed data, as determined by a distance threshold. If they are not all close to the data, we then walk backwards along the trajectory by one time step while keeping the number of points of interest on the integration fixed to the initially identified ones. This walking-back procedure stops when all currently visited integration path's data points' first-nearest points in the observed cells are sufficiently close.

To calculate the cell fate probability, we diffuse one step further of the identified nearest neighbors from the integration path (by identifying their first-nearest neighbor cells) to identify more nearest observed cells, especially those from terminal cell types. This is done to avoid the case in which the nearby cells first identified are all close to some random cells. We then use group information of those observed cells to define the fate probability. Fate probability for a particular cell group is then defined as:

$$1 - (sum(distances > "distancethreshold" \times "mediandistance") + "walkbacksteps")/ \\ (len(indices) + "walkbacksteps") \ .$$

The "*distance*" is the distances between currently visited integration path's data points and their first-nearest points in the observed cells belonging to a particular cell group. "*Median distance*" is the median distance of their diffused first- nearest cell distance of all observed cells. "*Walkback steps*" is the number of steps walked backward along the integration path until the distance between all currently visited integration points' first-nearest points to the observed cells satisfy the distance threshold. "*indices*" are the time indices of integration points that are regarded as the regions with small velocities. Note when walking backward, those corresponding points do not necessarily have small velocity anymore.

### Animating the single-cell trajectories on two-dimensional vector field space

Animating cell fate commitments relies on initial value integration with reconstructed vector field functions, as in the above section. Note that this two-dimensional space can be either 2D UMAP, any two dimensions from PCA, or any two genes of interest, as long as we first reconstruct the vector field on this two-dimensional space. A vector field animation, in many ways similar to molecular dynamics simulation, visualizes the exact speed of a set of cells at each time point, its movement in gene expression space, and the long-range trajectory predicted by the reconstructed vector field. Thus, it provides intuitive visual understandings of the RNA velocity, curvature, acceleration, and cell fate commitment in action.

### Methods for defining confidence of genes in the context of phase plane, velocity vectors, and fixed points

#### Correcting RNA velocity flow by removing genes with low gene-wise confidence in the phase plane

In some scenarios, we may find unexpected wrong velocity backflow from your RNA velocity analysis. To diagnose those cases, we can identify genes showing up in the wrong phase portrait position that may contribute to the wrong flow direction. We can then remove those genes to correct velocity vectors. This requires us to give some priors about the progenitor and terminal cell types in our system. The underlying rationale boils down to understanding the following two typical scenarios (**Fig. SI6F**):

1) If the expression of a particular gene in the progenitor is low, starting from time point 0, this gene's expression should start to increase as cells differentiate from progenitor to terminal cell states. There must be progenitors that are above the steady-state fitting line in the phase plane. However, if most of the progenitor cells are located below the line, we will have negative velocity, leading to reversed vector flow.

2) If the expression of a particular gene in progenitors is high, starting from time point 0, its expression should start to decrease. There must be progenitors that are below the steady-state fitting line. However, if most of the progenitors are located above the steady-state line, we will have positive velocity, again leading to reversed vector flow. Similar rationale can be applied to the mature cell states.

Thus, we design a heuristic algorithm to quantify the confidence of each gene by assessing whether it obeys the above constraints:

- We first assess whether, when each progenitor state differentiates into each terminal cell state, a gene is in the induction or repression phase based on the shift of the median gene expression between these two states. If it is in the induction phase, cells should mostly have positive or close to zero velocity (e.g. a small negative velocity threshold) and vice versa. Those thresholds can be provided by the users or inherited from the default values provided by ***dynamo***.
- 1 - fraction of cells having velocity passing those thresholds in each state is then used as a measure of velocity confidence.

Note that this heuristic method requires one to provide meaningful progenitor groups and mature cell groups, and the thresholds of velocity. In particular, the progenitor groups should in principle have cells going out (transcriptomically), whereas mature groups should end up in a different expression state, and there are intermediate cells going to the dead end cells in each terminal group (or most terminal groups).

#### Cell-wise confidence of RNA velocity vectors

Several confidence metrics for cell-wise velocity vectors are implemented in ***dynamo***. By default it uses the Jaccard index, which measures how well each velocity vector meets the geometric constraints defined by the local neighborhood structure [23]. The Jaccard index is calculated as the fraction of the number of the intersected set of nearest neighbors from each cell at the current expression state ($\boldsymbol{x}$) and that from the future expression state ($\boldsymbol{x} + \boldsymbol{v}$) over the number of the union of these two sets, namely:

$$\boldsymbol{J} = \frac{S(x_i) \cap S(x_i + v_i)}{S(x_i) \cup S(x_i + v_i)} \ .$$

$x_i, v_i, S(x_i)$, and $S(x_i + v_i)$ are respectively the current expression state for cell $i$, the current velocity vector for cell $i$, the set of nearest neighbor cells for cell $i$ based on the current expression states ($\boldsymbol{x}$), and the set for nearest neighbor cells for cell $i$ based on the future expression states ($\boldsymbol{x} + \boldsymbol{v}$). $\cap, \cup$ indicates the intersection or the union of two sets.

The cosine or correlation method is similar to that used by *scVelo* [3] and is used in **Fig. 3A** to quantify the local consistency of the velocity flow for each cell.

##### Confidence of identified fixed points

We notice that some identified fixed points are far away from regions populated with data points, where the reconstructed vector field may be less reliable. We quantify the confidence of the fixed points based on how far they are from domains populated with cells, and use the filled color of each node (corresponds to the fixed points) to represent the confidence of those fixed points when creating the topography plot in ***dynamo***.

#### Statistical tests used in this study

Mann–Whitney–Wilcoxon two-sided tests with Bonferroni correction are used to compare the distribution differences in **Fig. 5G**, **L**, **Fig. 7N**, **F**, **Fig. SI4D**, **Fig. SI5C** as well as **Fig. SI7Q** to identify the differentially expressed or accelerated gene. The default hypergeometric test, from gseapy for GO enrichment analysis is used in **Fig. 6H**, **Fig. 7I**, **K**, **Fig. SI2F**, **Fig. SI7D**, **J**, **N**.

[4] Tiejun Li, Jifan Shi, Yichong Wu, and Peijie Zhou. On the mathematics of RNA velocity i: Theoretical analysis. September 2020.

[15] Damian Smedley, Syed Haider, Steffen Durinck, Luca Pandini, Paolo Provero, James Allen, Olivier Arnaiz, Mohammad Hamza Awedh, Richard Baldock, Giulia Barbiera, Philippe Bardou, Tim Beck, Andrew Blake, Merideth Bonierbale, Anthony J Brookes, Gabriele Bucci, Iwan Buetti, Sarah Burge, Cédric Cabau, Joseph W Carlson, Claude Chelala, Charalambos Chrysostomou, Davide Cittaro, Olivier Collin, Raul Cordova, Rosalind J Cutts, Erik Dassi, Alex Di Genova, Anis Djari, Anthony Esposito, Heather Estrella, Eduardo Eyras, Julio Fernandez-Banet, Simon Forbes, Robert C Free, Takatomo Fujisawa, Emanuela Gadaleta, Jose M Garcia-Manteiga, David Goodstein, Kristian Gray, José Afonso Guerra-Assunção, Bernard Haggarty, Dong-Jin Han, Byung Woo Han, Todd Harris, Jayson Harshbarger, Robert K Hastings, Richard D Hayes, Claire Hoede, Shen Hu, Zhi-Liang Hu, Lucie Hutchins, Zhengyan Kan, Hideya Kawaji, Aminah Keliet, Arnaud Kerhornou, Sunghoon Kim, Rhoda Kinsella, Christophe Klopp, Lei Kong, Daniel Lawson, Dejan Lazarevic, Ji-Hyun Lee, Thomas Letellier, Chuan-Yun Li, Pietro Lio, Chu-Jun Liu, Jie Luo, Alejandro Maass, Jerome Mariette, Thomas Maurel, Stefania Merella, Azza Mostafa Mohamed, Francois Moreews, Ibounyamine Nabihoudine, Nelson Ndegwa, Céline Noirot, Cristian Perez-Llamas, Michael Primig, Alessandro Quattrone, Hadi Quesneville, Davide Rambaldi, James Reecy, Michela Riba, Steven Rosanoff, Amna Ali Saddiq, Elisa Salas, Olivier Sallou, Rebecca Shepherd, Reinhard Simon, Linda Sperling, William Spooner, Daniel M Staines, Delphine Steinbach, Kevin Stone, Elia Stupka, Jon W Teague, Abu Z Dayem Ullah, Jun Wang, Doreen Ware, Marie Wong-Erasmus, Ken Youens-Clark, Amonida Zadissa, Shi-Jian Zhang, and Arek Kasprzyk. The

BioMart community portal: an innovative alternative to large, centralized data repositories. Nucleic Acids Res., 43(W1):W589–98, July 2015.

[16] Lauren M Saunders, Abhishek K Mishra, Andrew J Aman, Victor M Lewis, Matthew B Toomey, Jonathan S Packer, Xiaojie Qiu, Jose L McFaline-Figueroa, Joseph C Corbo, Cole Trapnell, and David M Parichy. Thyroid hormone regulates distinct paths to maturation in pigment cell lineages. Elife, 8, May 2019.

[17] Kazumitsu Maehara and Yasuyuki Ohkawa. Modeling latent flows on single-cell data using the hodge decomposition. March 2019.
