## Supplementary figures and images for "Mapping Transcriptomic Vector Fields of Single Cells"

### Supplementary animation

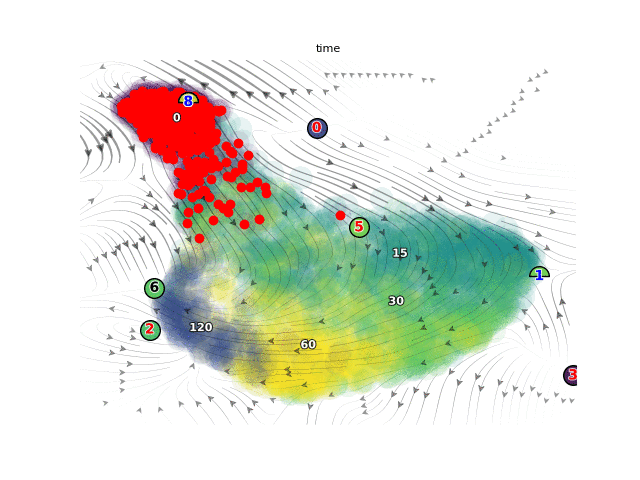

### Supplementary animation

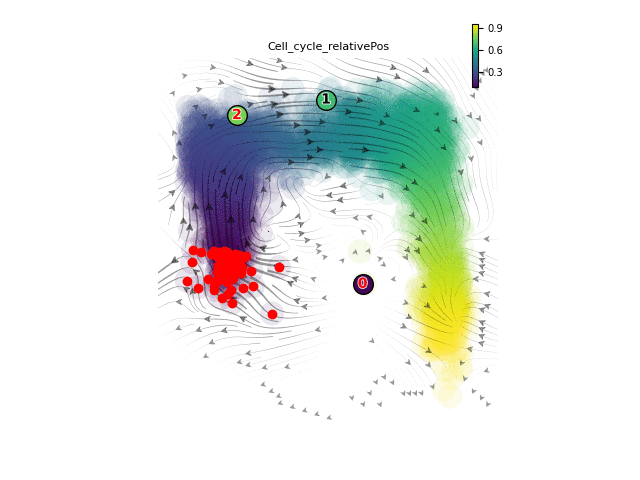

### Supplementary animation

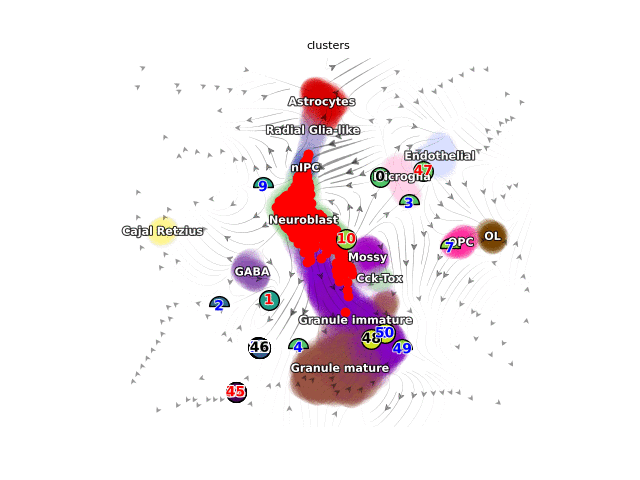

### Supplementary animation

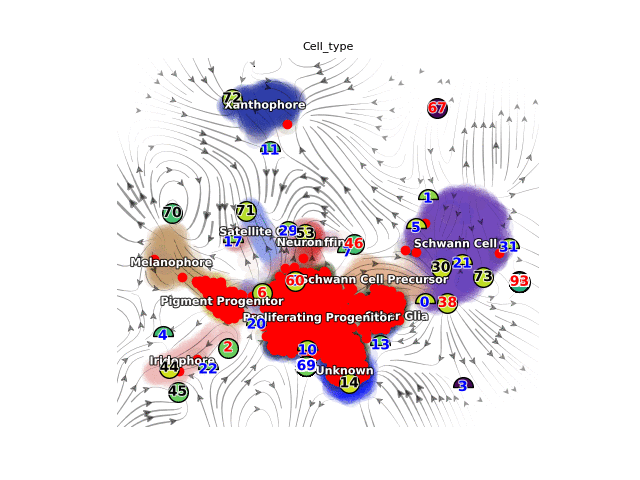

### Supplementary animation

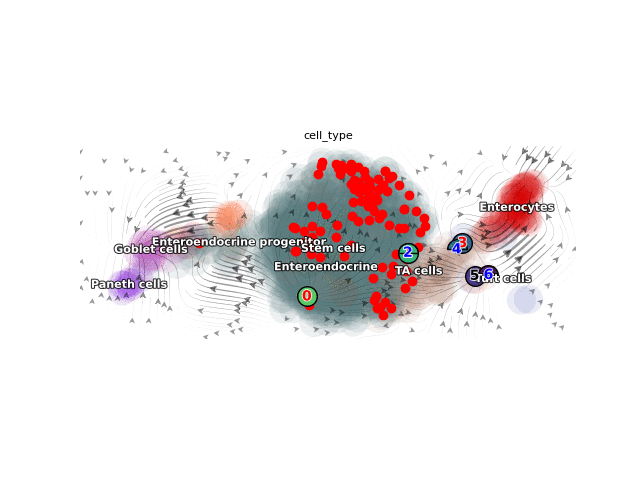

### Supplementary animation

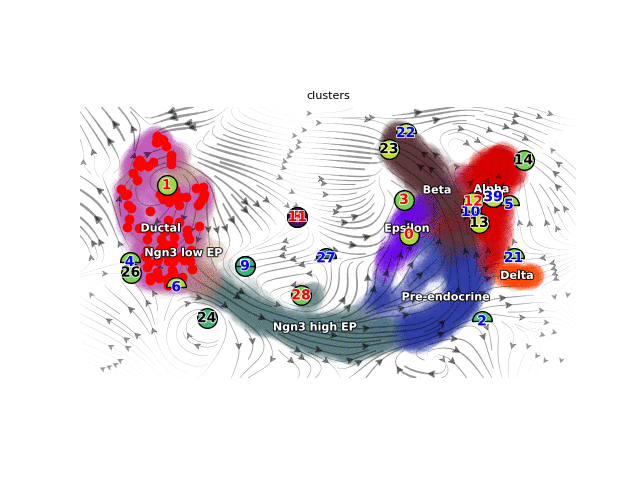
